# Sort-purification of human CD34^+^CD90^+^ cells reduces target cell population and improves lentiviral transduction for gene therapy

**DOI:** 10.1101/850479

**Authors:** Stefan Radtke, Dnyanada Pande, Margaret Cui, Anai M. Perez, Yan-Yi Chan, Mark Enstrom, Stefanie Schmuck, Andrew Berger, Tom Eunson, Jennifer E. Adair, Hans-Peter Kiem

## Abstract

Hematopoietic stem cell (HSC) gene therapy has the potential to cure many genetic, malignant and infectious diseases. We have shown in a nonhuman primate (NHP) HSC gene therapy and transplantation model that the CD34^+^CD90^+^ cell fraction was exclusively responsible for multilineage engraftment and hematopoietic reconstitution. Here we show the translational potential of this HSC-enriched CD34 subset for lentivirus-mediated gene therapy. Alternative HSC-enrichment strategies include the purification of CD133^+^ cells or CD38^low/-^ subsets of CD34^+^ cells from human blood products. We directly compared these strategies to the isolation of CD90^+^ cells using a GMP-grade flow-sorting protocol with clinical applicability. We show that CD90^+^ cell selection results in 40-fold fewer target cells in comparison to CD133^+^ or CD38^low/-^ CD34 subsets without compromising the engraftment potential *in vivo*. Single cell RNA sequencing confirmed nearly complete depletion of lineage committed progenitor cells in CD90^+^ fractions compared to alternative selections. Importantly, lentiviral transduction efficiency in purified CD90^+^ cells resulted in up to 3-fold higher levels of engrafted gene-modified blood cells. These studies should have important implications for the manufacturing of patient-specific HSC gene therapy and genome editing products.

## INTRODUCTION

HSC-mediated gene therapy or editing for the treatment of hematologic disease is commonly performed with single-marker purified CD34^+^ cells. However, it is well-established that CD34^+^ cells are a heterogeneous mix of predominantly lineage-committed progenitor cells containing only very few true HSCs with long-term multilineage engraftment potential (1, 2). Moreover, the proportion of true HSCs is variable across individuals, requiring at least 2 x 10^6^ viable CD34^+^ cells per kg of body weight to meet safety and feasibility standards. This results in excessive use of costly reagents (e.g. lentiviral vectors or nucleases) for gene transfer to reliably target HSCs (3–9). Moreover, the efficiency of gene-modification seen *ex vivo* does not correlate with the *in vivo* engraftment of modified cells commonly stabilizing at a significantly lower frequency (4, 10). These major hurdles significantly limit the broad availability of this very promising technology as a potential cure for many patients.

HSC gene therapy and transplantation would benefit from the ability to isolate, target, and modify a more HSC-enriched subset that provides short-term reconstitution as well as long-term multilineage engraftment. Availability of a refined target would overcome all currently existing limitations simultaneously: 1) greatly reduce the amount of modifying reagents needed for manufacturing, 2) result in more reliable genetic-modification of HSCs, and 3) increase the predictability of transplant success *in vivo*.

Within the last four decades, a plethora of cell surface markers have been described for the identification and purification of human HSCs (1, 11–15). Similarly, a great variety of HSC-enrichment strategies has been proposed for gene therapy approaches. Some of the most promising candidate HSC-enrichment strategies include the purification of the CD38^low/–^ (16, 17), CD133^+^ (18), or CD90^+^ (19) CD34 subsets. While each strategy claims to improve individual aspects of gene therapy and editing, a side by side analysis of these phenotypes has not been reported.

Here, we applied up-to-date read-outs (*in vitro*, *in silico,* and *in vivo*) for a systematic and objective side-by-side comparison of candidate HSC-enrichment strategies with the goal to identify the most refined HSC-enriched target for gene therapy/editing and transplantation.

## RESULTS

### CD34^+^CD90^+^ cells are phenotypically the most refined target for HSC gene therapy

Multilineage long-term engraftment and bone marrow (BM) reconstitution is driven by CD34^+^CD90^+^ hematopoietic stem and progenitor cells (HSPCs) in the pre-clinical NHP stem cell transplantation and gene therapy model (19–21). To evaluate the translational potential of the CD34^+^CD90^+^ phenotype, we compared this subset to alternative HSC-enrichment strategies including CD34^+^CD133^+^ (18) and CD34^+^CD38^low/–^ (16, 17) HSPCs.

To compare the fold-reduction of target cells for each strategy, we initially analyzed the frequency of CD133^+^, CD38^low/-^ and CD90^+^ subsets in steady-state bone marrow (ssBM)-derived CD34^+^ cells across nine different healthy donors (**Figure 1A, B**). On average 42% (±2.8%, SD) of CD34^+^ cells co-expressed CD133 on the cell surface reducing the target cell count for HSC-gene therapy approximately 2.4-fold. Lack of CD38 expression (CD38^low/-^) was observed on 16.5% (±5.2%, SD) of CD34^+^ cells or a 5.8-fold enrichment of more primitive HSCs. 12.5-fold reduction of target cells was reached for CD90^+^ cells comprising on average 7.5% (±4.7%, SD) of bulk CD34^+^ HSPCs.

**Figure 1.**
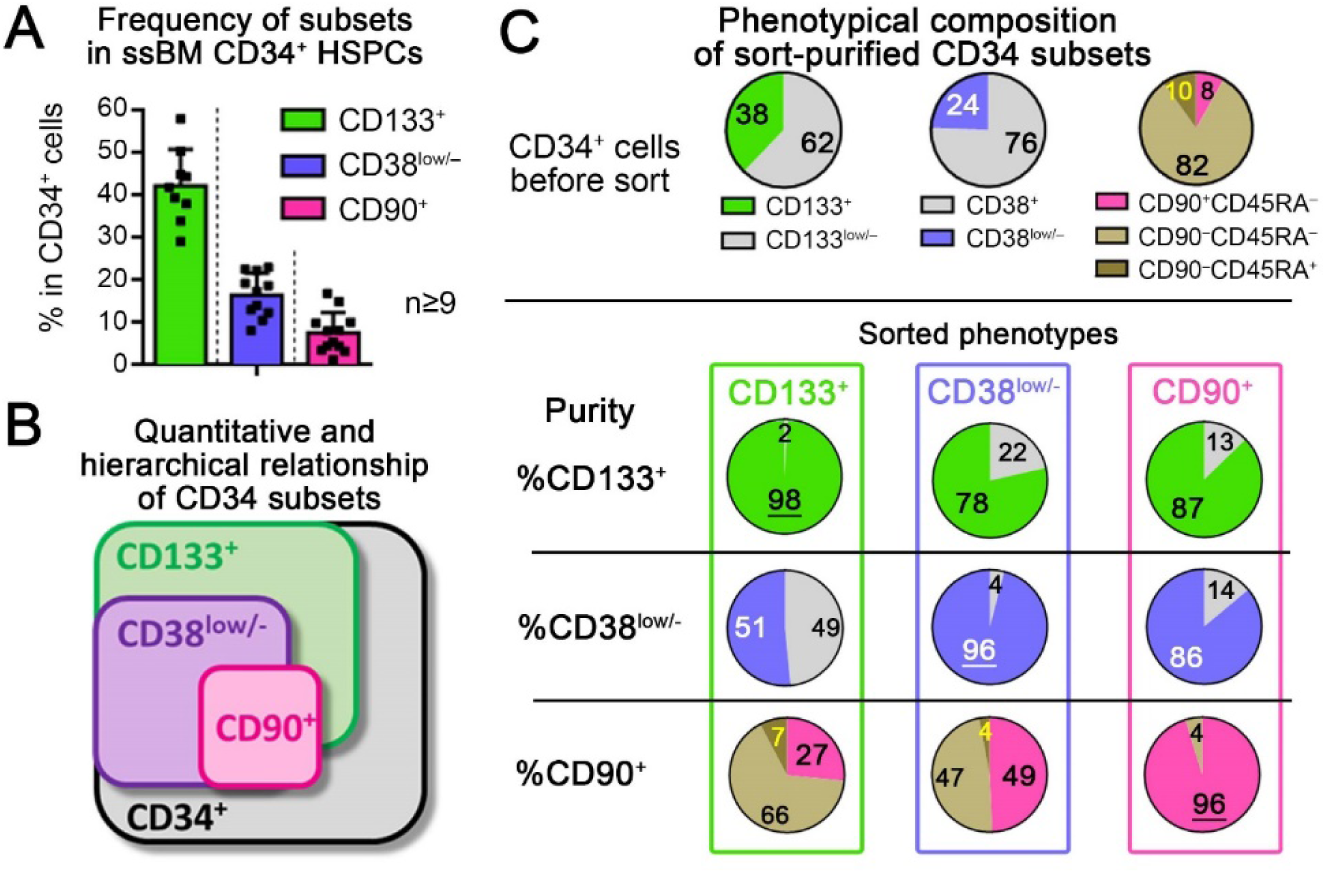
Quantitative comparison of phenotypically defined target CD34 subsets. (**A**) Average frequency of CD133^+^, CD38^low/–^ and CD90^+^ HSPCs within bulk ssBM-derived CD34^+^ cells (mean ± SD, n≥9 independent healthy donors). (**B**) Schematic of the quantitative and hierarchical relationship of phenotypical CD34 subsets. (**C**) Frequency of phenotypically defined subsets in bulk CD34^+^ cells before sort (top row) and cross-contamination of phenotypical subsets within sort-purified CD133^+^ (2^nd^ row), CD38^low/–^ (3^rd^ row) and CD90^+^ (4^th^ row) HSPCs.

Since the purification of CD90^+^ HSPCs is currently limited to fluorescence assisted cell sorting (FACS), we next evaluate the feasibility of flow-sorting for the enrichment of human HSC-enriched CD34 subsets. All three phenotypes were sort-purified and HSC-enriched subpopulations analyzed for contaminating cell types (**Figure 1C** and **Supplemental Figure 1**). The CD133^+^ subpopulation showed approximately 50% enrichment of CD38^low/–^ but only 27% of the CD90^+^ subset. Purified CD38^low/–^ cells were markedly enriched for CD133^+^ (78%) and CD90^+^ (49%) cells but still contained a significant proportion of CD90^−^ HSPCs (51%). CD90^+^ cells demonstrated more than 86% enrichment of CD133^+^ as well as CD38^low/–^ HSPCs.

Based on this phenotypic assessment, CD90^+^ HSPCs demonstrate the most refined HSC-enriched CD34 subset. Purification of this phenotype yields the greatest reduction of target cells for HSC gene therapy in comparison to CD133^+^ or CD38^low/– CD34^ subsets.

### CD90^+^ HSPCs are depleted for transcriptionally lineage-committed progenitors

Single cell transcriptome analysis has proven useful in determining the heterogeneity of complex cell populations (22). Here, we used single-cell RNA sequencing (scRNAseq) on the 10X Genomics chromium platform to assess the transcriptional heterogeneity and depletion of lineage-committed progenitor cells within sort-purified CD133^+^, CD38^low/-^, and CD90^+^ CD34 subsets.

To identify transcriptionally distinct CD34 subsets of primitive HSPCs and lineage-committed progenitor cells, we initially generated a scRNAseq reference map. CD34^+^ cells from ssBM of two healthy human individuals were sequenced (**Supplemental Table 1**) and the data analyzed using t-distributed stochastic neighbor embedding (tSNE) and principle component analysis (PCA).

Seven transcriptionally distinct groups of single cells were identified within CD34^+^ cell populations using the Seurat un-supervised graph-based clustering tool (23) (**Supplemental Table 2**). Clusters were mapped onto both the tSNE (**Figure 2A**, **Supplemental Figure S2A**) and PCA (**Figure 2B**, **Supplemental Figure S2B**) visualizations. Comparison of clusters demonstrated expression of genes associate with primitive HSPCs predominantly localized in cluster 1 (**Supplemental Figures 3** and **4**). Increased expression of genes for lymphoid, myeloid, and erythroid differentiation were observed in clusters 2 and 3, clusters 4 and 5, and clusters 6 and 7, respectively. CD34 expression was detectable in all clusters, with a gradual increase towards more primitive HSPCs (**Figure 2C** and **Supplemental Figure 5A**). In good agreement to previous publications (1, 24–27), CD133 mRNA expression was restricted to transcriptionally lympho-myeloid committed progenitors as well as immature HSPCs (clusters 1-5). CD38 expression was scattered throughout the reference map and showed no obvious association or absence in distinct clusters. Surprisingly, CD90 expression was undetectable in both donors.

**Figure 2.**
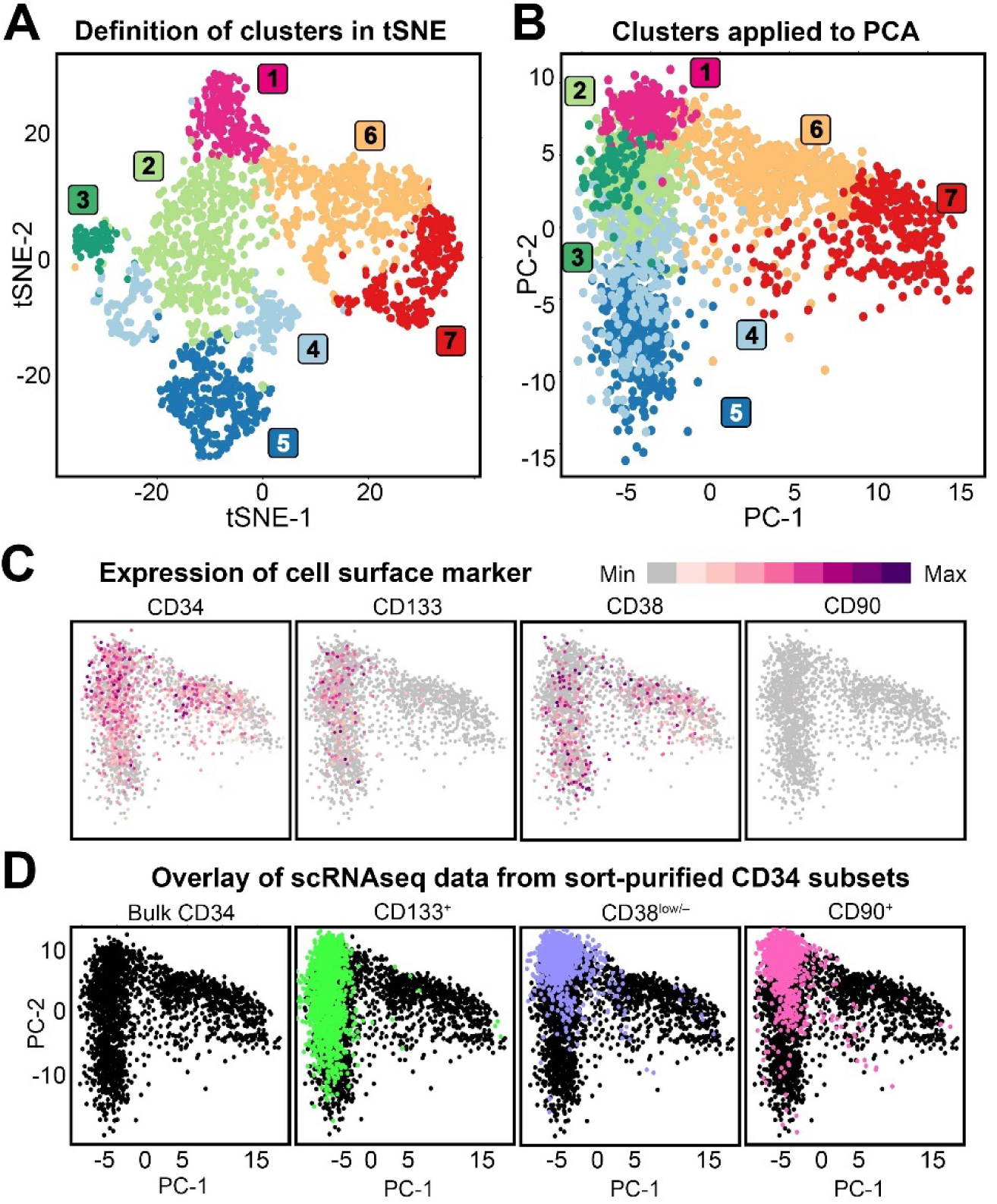
ScRNAseq of ssBM-derived CD34^+^ HSPCs and sort-purified CD34 subsets. (**A**) Dimensional reduction (tSNE) and graph-based clustering (PCA) of scRNAseq data from ssBM-derived CD34^+^ cells. (**B**) PCA based transformation with clusters defined in panel A projected onto the PCA analysis. (**C**) Expression of CD34, CD133, CD38, and CD90 in ssBM-derived CD34^+^ cells. Level of expression is color coded as shown in the legend. (**D**) Overlay of scRNAseq data from CD34^+^ cells (black, 1^st^ plot) with sort-purified CD133^+^ (green, 2^nd^ plot), CD38^low/–^ (purple, 3^rd^ plot), and CD90^+^ (pink, 4^th^ plot) HSPCs.

To now compare phenotypical CD34 subsets, we sort-purified CD133^+^, CD38^low/–^, and CD90^+^ cell fractions, performed scRNAseq, and overlaid the resulting data onto the CD34 reference map (**Figure 2D**, **Supplemental Table 1 and Supplemental Figure 5B**). Similar to the observed CD133 mRNA expression, sort-purified CD133^+^ HSPCs exclusively colocalized with lympho-myeloid and more primitive HSPCs. Sort-purified CD38^low/–^ cells predominantly overlaid with cluster 1, with individual cells spreading into both the lympho-myeloid as well as the erythroid branch. Surprisingly, mapping of sort-purified CD90^+^ cells was almost indistinguishable from CD38^low/–^ cells demonstrating a significant enrichment of immature HSPCs (cluster 1), as well as depletion of lympho-myeloid as well as erythroid primed progenitors.

Since CD90^+^ and CD38^low/-^ cells show significant phenotypical differences but are undistinguishable by scRNAseq, we hypothesized that the mRNA capture efficiency of scRNAseq may be insufficient (28–31) to detect subtle transcriptional difference in between both phenotypes. Consequently, we performed bulk-RNAseq on CD90^+^ (Population *a*) and CD38^low/-^ (Population *e*) cells and mapped the data onto our scRNAseq reference map (**Figure 3A**). To validate this “bulk onto scRNAseq” mapping strategy, additional CD34 subsets enriched for lympho-myeloid and erythro-myeloid progenitor cells were added. Briefly, HSPCs were sorted into CD34^+^CD90^+^ (Population *a*), CD34^+^CD90^−^CD45RA^−^CD133^+^ (Population *b*), CD34^+^CD90^−^CD45RA^+^CD133^+^ (Population *c*), and CD34^+^CD90^−^CD45RA^−^CD133^low/–^ (Population *d*) subsets (26, 32, 33) as well as CD34^+^CD38^low/–^ (Population *e*), CD34^+^CD38^low/–^CD90^+^ (Population *f*), and CD34^+^CD38^low/–^ CD90^−^ (Population *g*) populations (16, 17, 27) (**Figure 3A** and **Supplemental Figure 6**).

**Figure 3.**
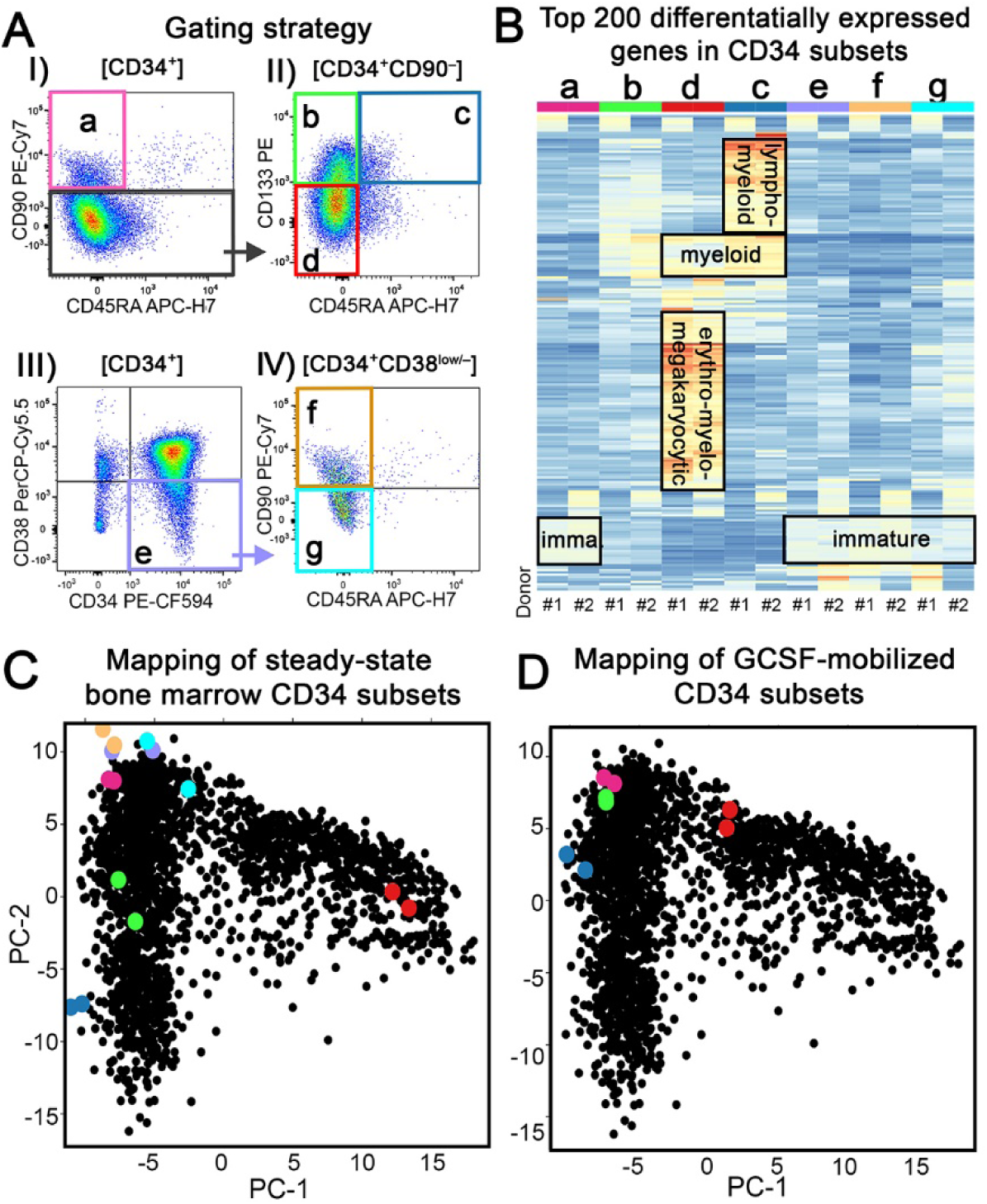
Bulk RNAseq of ssBM-derived CD34 subsets. (**A**) Gating strategy for phenotypically-defined human CD34 subpopulations. (**B**) Heat map of the Top 200 differentially expressed genes in phenotypically-defined CD34 subsets listed in panel A from two independent human donors. A detailed list of the Top 200 genes can be found in **Supplemental Table 3**. (**C**) Overlay of bulk RNAseq data from sort-purified ssBM subsets (Populations *a*–*g* color coded as defined in panel A) on the ssBM scRNAseq PCA reference map (black dots) (**D**) Overlay of bulk RNAseq data from sort-purified GCSF-mobilized CD34 subsets (Population *a*–*d* color coded as defined in panel A) on the ssBM scRNAseq PCA reference map (black dots).

Consistent with the reported lineage potentials of the sort-purified CD34 subsets (16, 17, 19, 26, 27, 32, 33), we observed upregulation of lymphoid genes (*JCHAIN*, *DNTT*, *CD2*, *IGLL1*) in Population *c*, erythro-myelo-megakaryocytic genes (*HBB*, *HDC*, *GATA1*, *CD36*) enriched in Population *d*, and expression of more immature marker genes (*AVP*, *HES1*, *DLK1*) in Populations *a*, *e*, *f* and *g* (**Figure 3B**, **Supplemental Table 3**). Individual myeloid genes (*ELANE*, *MPO*, *CEBPD*) were simultaneously upregulated in Populations *c* and *d*. Interestingly, Population *b* did not show a unique cluster of differentially expressed genes but shared features of genes associated with lympho-myeloid and myeloid-primed as well as immature HSPCs (**Figure 3B**, **Supplemental Table 3**).

To confirm this manual assessment, we mapped the bulk-RNAseq data onto the CD34 scRNAseq reference map (**Figure 3C**). As expected, lympho-myeloid primed cells (Population *c*) mapped within clusters 4 and 5 and erythro-myelo-megakaryocytic primed HSPCs (Population *d*) plotted within cluster 7. Population *b* showed greater heterogeneity (distance in between dots) and localized within clusters 2 and 3 of the lympho-myeloid arm. Populations *a*, *e*, *f*, and *g* were closely co-localized within cluster 1 at the top of the reference map. More detailed comparison of Populations *a*, *e*, *f*, and *g* revealed that CD90^+^ HSPCs (Population *a*) demonstrated the lowest donor to donor variability. In contrast, CD38^low/–^ (Population *e*) HSPCs showed the greatest heterogeneity despite nearly identical proportions of CD38^low/–^CD90^+^ (36/33%) and CD38^low/–^ CD90^−^ (64/67%) subsets in both donors. CD38^low/–^CD90^−^ (Population *g*) subsets were similarly heterogeneous and shifted towards the erythroid-primed clusters matching the reported enrichment of erythro-myeloid progenitors in CD133^low/–^CD38^−^CD90^−^ HSPCs (33, 34). Most importantly, sorting of the CD38^low/–^CD90^+^ subset (Population *f*) significantly reduced the donor to donor variability and led to closer co-localization with CD90^+^ HSPCs (Population *a*).

Finally, we sort-purified Population *a*-*d* HSPCs from two GCSF-mobilized CD34^+^ donors, the most frequently used stem cell source for HSC gene therapy and editing approaches, and bulk RNAseq data was mapped onto the reference map (**Figure 3D**). No meaningful differences in the transcriptional signature were observed in between CD90^+^ (Population *a*) HSPCs from both sources (**Supplemental Table 3 and 4**). However, CD90^−^CD45RA^−^CD133^+^ (Population *b*) HSPCs reported to contain multipotent progenitors (27, 35) co-localized much closer with CD90^+^ HSPCs (Population *a*). Pairwise comparison of CD90^+^ (Population *a*) with CD90^−^CD45RA^−^ CD133^+^ (Population *b*) HSPCs confirmed that GCSF mobilization significantly reduced the transcriptional heterogeneity in between both CD34 subsets (ssBM: 615 genes; GCSF: 131 genes; **Supplemental Figure 7**, **Supplemental Tables 5–8**).

In conclusion, combining single cell and bulk RNAseq we here show that CD90^+^ HSPCs are almost entirely depleted for transcriptionally lineage committed progenitor cells and significantly enriched for primitive HSPCs. Most importantly for the clinical application, the transcriptional signatures within the CD90^+^ subset is not impacted by GCSF-mediated mobilization.

### BM reconstitution potential is restricted to CD34^+^CD90^+^ cell fractions

Pre-clinical experiments in the NHP and mouse showed that GCSF-primed CD34^+^CD90^+^ HSPCs were solely responsible for the reconstitution of the BM stem cell compartment (19, 21).To ensure that the depletion of CD90^-^ HSPCs in humans does not impact the *in vivo* multilineage engraftment potential, Population *a*-*d* cells from human GCSF-mobilized CD34^+^ HSPCs were sort-purified and transplanted into sub-lethally irradiated adult NSG mice.

The highest engraftment in the peripheral blood (PB), BM, and thymus was observed in cohorts transplanted with CD90^+^ (Population *b*) cells (**Figure 4A,B**, **Supplemental Figure 8A-D, and Supplemental Figure 9A,B**). Lower level and less consistent multilineage engraftment of human cells was observed upon transplantation of Population *b*. Mice receiving Population *c* showed locally restricted human chimerism in the thymus, whereas Population *d* did not show any human engraftment in the analyzed tissues. Engraftment and reconstitution of the entire BM stem cell compartment including the recovery of phenotypically primitive human CD34^+^CD90^+^ HSPCs was exclusively observed after transplantation of CD90^+^ cells (**Figure 4C**, **Supplemental Figure 8E,F, and Supplemental Figure 9C**). Similarly, erythroid, myeloid, and erythro-myeloid colony-forming cell (CFC) potentials were only detected in mice transplanted with CD90^+^ cells (**Figure 4D**, **Supplemental Figure 9D**).

**Figure 4.**
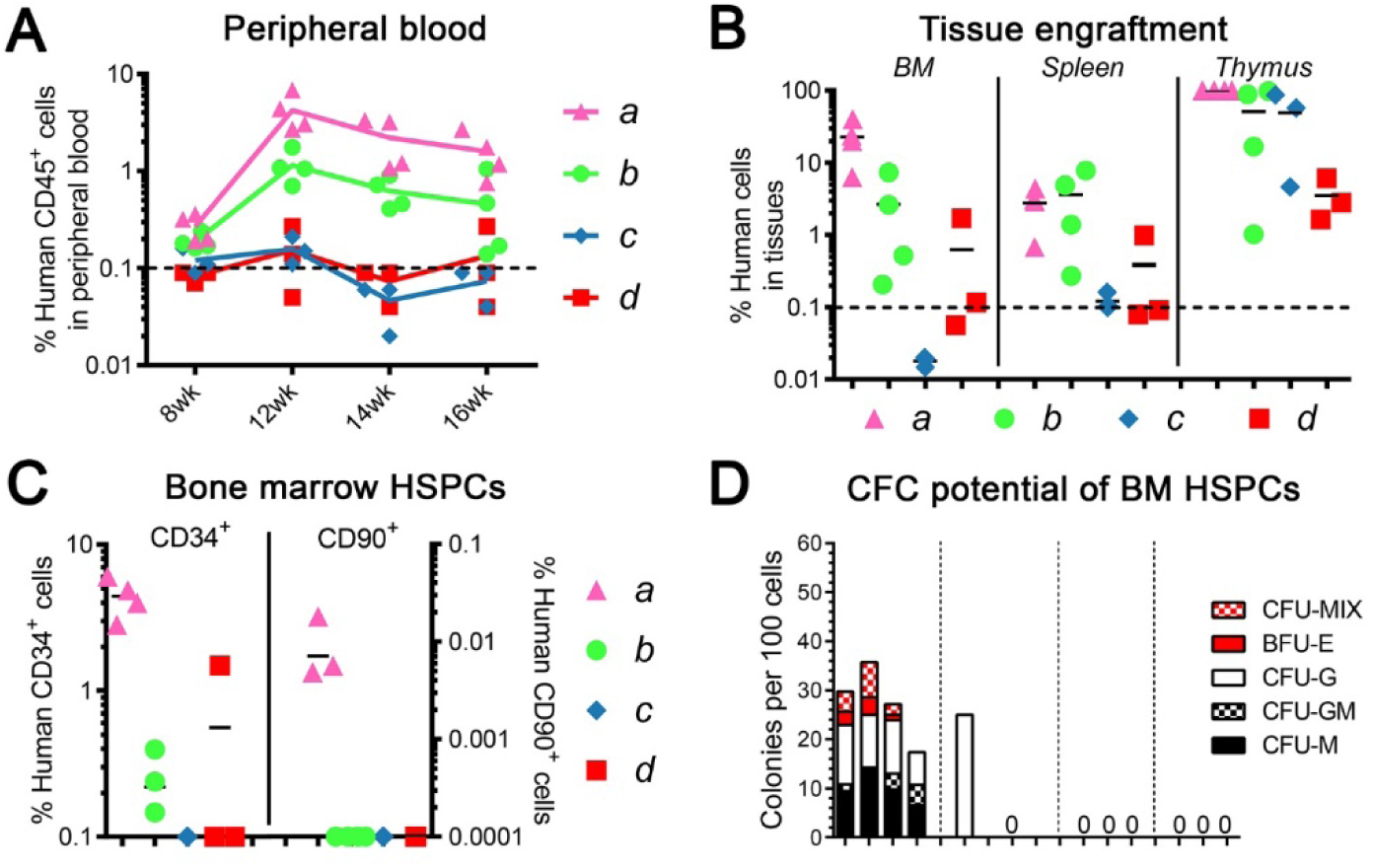
Multilineage engraftment potential of human CD34 subpopulations. Frequency of human chimerism in the (**A**) PB and (**B**) BM, spleen, and thymus after transplantation of sort-purified CD34 subpopulations (1×10^5^ cells per mouse). (**C**) Frequency of engrafted human CD34^+^ and CD90^+^ HSPCs. CD34^+^ frequency use left y-axis, CD90^+^ frequency right y-axis. (**D**) Erythroid, myeloid and erythro-myeloid colony-forming cell (CFC) potential of engrafted human CD34^+^ cells. CFU, colony-forming unit; CFU-M, macrophages; CFU-G, granulocytes; CFU-GM, granulocyte/macrophage; and BFU-E, erythroid; CFU-MIX, erythro-myeloid colonies.

To confirm that human CD90^−^CD45RA^−^CD133^+^ HSPCs (Population *b*) do not contain primitive HSCs with multilineage long-term engraftment potential, we performed limiting dilution experiments (**Supplemental Figure 10**). Gradually increasing the number of transplanted cells from Population *b* led to greater multilineage engraftment of human cells in all tissues including CD34^+^ cells in the BM stem cell compartment (**Supplemental Figure 10A-E**). However, none of the mice demonstrated human CD34^+^CD90^+^ HSPCs *in vivo* after transplant with this population, and engrafted CD34^+^ cells were restricted to erythroid and myeloid colony types lacking mixed CFU potentials (**Supplemental Figure 10F**). The number of SRCs (SCID-repopulating cells) in Population *b* was calculated to be 1 in 4.6×10^5^ transplanted cells (**Supplemental Figure 10G**).

In summary, mouse xenograft experiments confirm enrichment of primitive human HSPCs with multilineage engraftment and BM reconstitution potential in the CD34^+^CD90^+^ phenotype (Population *a*). Furthermore, CD34^+^CD90^−^CD133^+^CD45RA^−^ HSPCs (Population *b*) contain low-level multilineage engraftment potential but lack the ability to recover the entire stem cell compartment.

### Sort-purification increases the transduction efficiency in HSC-enriched CD90^+^ cells

Our data indicates that CD90^+^ HSPCs are phenotypically, transcriptionally, and functionally the most refined target for HSC gene therapy. Consequently, we next evaluated the feasibility of a flow-cytometry based, good manufacturing practice (GMP)-compatible, large-scale sort-purification strategy for the enrichment and direct targeting of CD90^+^ cells in a clinical setting.

CD34^+^ cells from six healthy GCSF-mobilized donors were enriched on the Miltenyi CliniMACS Prodigy according to our previously established protocol (**Figure 5A**, **Supplemental Table 9**) (36). Leukapheresis products yielded on average 2.56×10^8^ CD34^+^ cells with a purity greater than 93.5±1.9%. Donor #4 insufficiently mobilized CD34^+^ cells and was excluded from the study.

**Figure 5:**
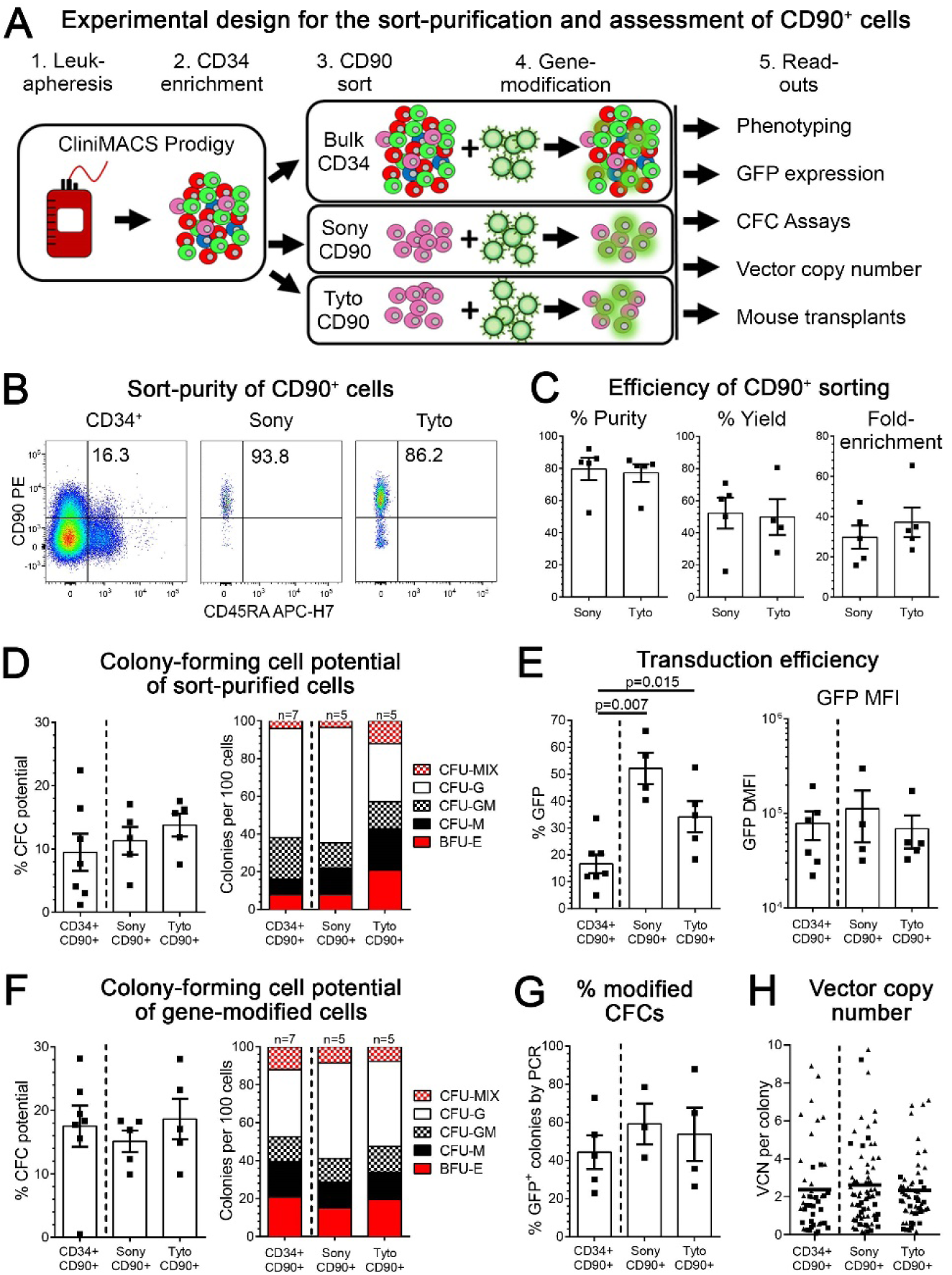
Sort-purification and quality control of CD90^+^ HSPCs. **(A)** Schematic of the experimental design. (**B**) Flow-cytometric assessment of cells before sort (CD34^+^, 1^st^ plot) and purified CD90^+^ cells after sorting on the Sony (2^nd^ plot) and Tyto (3^rd^ plot). (**C**) Comparison of the purity, yield and fold-enrichment of CD90^+^ HSPCs on the Sony and Tyto sorter. (**D**) CFC potential of CD90^+^ cells within bulk CD34^+^ HSPCs (CD34^+^CD90^+^) and sort-purified CD90^+^ subsets. (**E**) Flow-cytometric quantification of GFP-expressing CD90^+^ cells within bulk CD34^+^ cell and sort-purified CD90^+^ subsets (left) and the delta-MFI of GFP expression in gene-modified cells (right). (**F**) Erythroid, myeloid and erythro-myeloid CFC potential of gene-modified CD34^+^CD90^+^ and sort-purified CD90^+^ subset. Individual colonies from all three conditions in (F) were picked and (**G**) the gene-modification efficiency in CFCs determine by PCR as well as (**H**) the VCN in modified CFCs quantified by qPCR. (statistics: means ± SEM; significance values: two-tailed paired t test)

Sort-purification of CD90^+^ HSPCs was performed comparing the jet-in-air sorter FX500 from Sony with the cartridge-based closed-system sorter MACSQuant Tyto from Miltenyi Biotech (**Figure 5A**). Sorted CD90^+^ cell fractions on both machines showed no significant phenotypical or quantitative differences. They both reached an average of 79.5±6.9% (Sony) and 77.1±5.5% (Tyto) purity with a yield of 52.3±9.6% and 49.9±11.1%, respectively (**Figure 5B**,C). Of special interest for HSC gene therapy, the average fold-enrichment of target cells was greater than 29.7±5.7-fold on the Sony sorter and up to 37.2±7.3-fold on the Tyto (**Figure 5C**). To determine whether the sort-purification impacted the differentiation potential of HSC-enriched CD34 subsets, cells were introduced into CFC assays. Sort-purified cell fractions did not demonstrate any differences in the total nor compositional CFC potential compared to unprocessed cells (**Figure 5D**).

Next, sorted CD90^+^ as well as bulk CD34^+^ cells were transduced with a lentivirus encoding for green fluorescent protein (GFP) according to our clinically approved CD34-mediated gene therapy protocol (37, 38). Five days post-transduction, the gene-modification efficiency in the CD90^+^ subset of bulk-transduced CD34 cells reached 16.5±3.4% (SEM), whereas 2.3-fold (Tyto: 38.1±5.5%, SEM, p=0.012) and 3.1-fold (Sony: 51.9±5.8%, SEM, p=0.006) higher frequencies of GFP were seen in the sort-purified CD90^+^ cell fractions (**Figure 5E**). Despite significant differences in the gene-modification efficiency, no significant increase in the overall mean fluorescence intensity (MFI) of GFP was observed transducing purified CD90^+^ cells (**Figure 5E**). Gene-modified cells were introduced into CFC assay showing no significant quantitative and qualitative differences (**Figure 5F**). Finally, individual colonies were extracted to determine the gene-modification efficiency within erythro-myeloid colonies as well as to precisely quantify the vector copy number (VCN). No obvious differences were found in the expression of GFP within CFCs from all three conditions (**Figure 5G**). Similar to the MFI of GFP, almost identical VCNs were observed in all colonies (**Figure 5H**).

In summary, the sort-purification of CD90^+^ cells is technically feasible and does not impact the cells phenotypical and functional properties. Most importantly, purification of CD90^+^ cells reduces the number of target cells and simultaneously improves the efficiency of lentivirus-mediated gene-transfer without increasing the MFI as well as VCN without the need for additional transduction enhancers.

### Improved in vivo engraftment of sort-purified and gene-modified CD90^+^ HSPCs

Regardless of high gene-modification efficiency *ex vivo*, the frequency of long-term *in vivo* engraftment of modified cells is often both unpredictable and, in many studies, significantly lower compared to values determined during quality control of the infusion product (10, 20, 39). To compare the multilineage long-term engraftment potential of gene-modified bulk CD34^+^ with sort-purified CD90^+^ HSPCs, cells were transplanted into sublethally irradiated adult NSG mice. Of note, in order to mimic the average percentage of CD90^+^ cells within bulk CD34^+^ cell fractions mice transplanted with sort-purified CD90^+^ cells received 1/10^th^ the cell number compared to mice transplanted with bulk CD34^+^ cells.

Multilineage engraftment of human cells in the PB of transplanted mice was followed longitudinally by flow-cytometry (**Figure 6A**). Mice transplanted with bulk CD34^+^ as well as Tyto-sorted CD90^+^ cells showed persisting levels of human engraftment, whereas human chimerism in the vast majority of mice receiving Sony-sorted CD90^+^ HSPCs gradually declined towards the end of study. Similar trends in the overall frequency of human multilineage engraftment were seen at the day of necropsy in the BM, spleen, and thymus (**Figure 6B**, **Supplemental Figure 11**). Overall, the greatest frequency of human chimerism was found in mice transplanted with bulk CD34^+^ and Tyto-sorted CD90^+^ HSPCs followed by Sony-sorted CD90^+^ cells.

**Figure 6:**
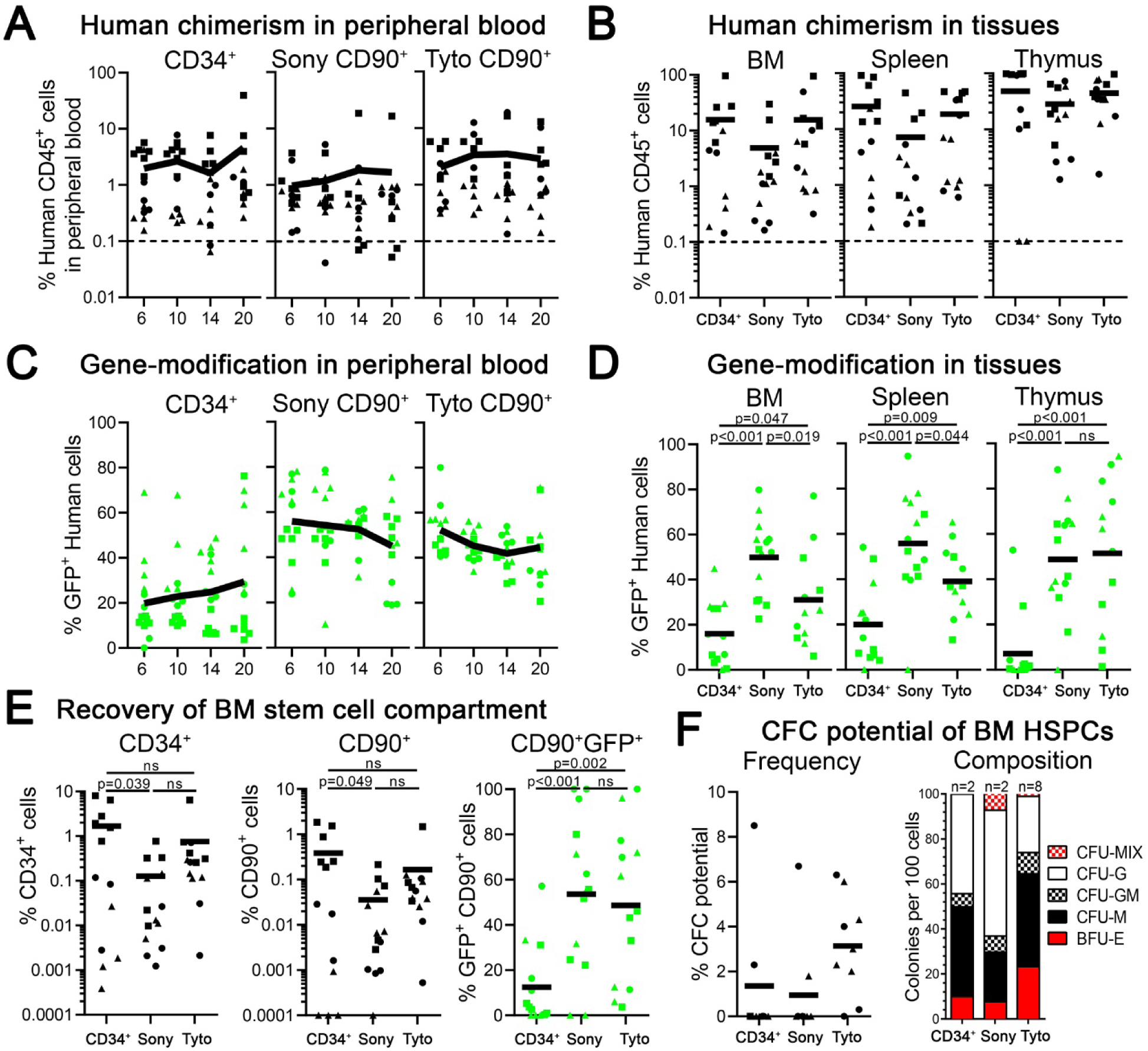
Transplantation of sort-purified and gene-modified CD90^+^ cells into NSG mice. (**A**) Frequency of human chimerism in the PB over time. (**B**) Human chimerism in the BM, spleen and thymus at 20 weeks post-transplant. Frequency of GFP^+^ human cells in (**C**) the PB over time and (**D**) tissues at 20 weeks post-transplant. (**E**) Engraftment of human CD34^+^, CD90^+^ and CD90^+^GFP^+^ HSPCs in the BM at necropsy. (**F**) Erythroid, myeloid and erythro-myeloid colony-forming potential of engrafted human CD34^+^ cells.

Lentiviral-mediated delivery of GFP enabled us to flow-cytometrically follow the frequency of gene-modification longitudinally in the PB as well as at the day of necropsy in tissues. Consistent with the efficiency of gene-modification we saw *ex vivo* (**Figure 5E**), a 2- to 3-fold higher frequency of GFP^+^ human cells was seen in the PB and tissues of animals receiving sort-purified CD90^+^ cells compared to bulk CD34^+^ cells (**Figure 6C,D**). The frequency of gene-modification in the PB of animals transplanted with sort-purified CD90^+^ cells remained consistent throughout the follow-up, whereas in animals receiving gene-modified bulk CD34^+^ cells two patterns were observed; human cells 1) either gradually lost the gene-modification leveling out at about 10-15% or 2) started to show a rapid increase in GFP^+^ human cells up to almost 80% towards the end of study.

Next, we comprehensively analyzed the BM stem cell compartment of engrafted mice. Mice transplanted with bulk CD34^+^ and Tyto-sorted CD90^+^ cells demonstrated significantly greater engraftment of human CD34^+^ and CD34^+^CD90^+^ cells compared to Sony-sorted CD90^+^ HSPCs (**Figure 6E**). Most importantly, the frequency of gene-modified CD90^+^ cells was in average 2- to 3-fold higher in mice engrafted with sort-purified and gene-modified CD90^+^ compared to bulk CD34^+^ HSPCs. Finally, we introduced BM-resident human CD34^+^ into CFC assays to test the maintenance of erythro-myeloid differentiation potentials (**Figure 6F**). Engrafted human HSPCs in only two mice transplanted with either bulk CD34^+^ or Sony-sorted CD90^+^ gave rise to colonies, whereas in 8 out of 9 cases erythro-myeloid colonies were seen in mice receiving Tyto-sorted CD90^+^ cells.

In summary, gene-modification of sort-purified CD34^+^CD90^+^ cells on the Tyto does not alter the multilineage long-term engraftment potential of human HSPCs in the mouse xenograft model. Most importantly, gene-editing efficiencies observed *ex vivo* successfully translated into the *in vivo* transplant and remained stable throughout the long-term follow-up.

## DISCUSSION

Here, we show that the enrichment of the human CD34^+^CD90^+^ population has the potential to overcome current limitations of HSC gene therapy. Sort-purification of this HSC-enriched CD34 subset resulted in up to 40-fold reduction of target cells and consequently significant savings of costly reagents (here lentiviral vectors). Most importantly, sole purification of this phenotype improved the gene-modification efficiency of HSCs with long-term *in vivo* engraftment potential up to 3-fold perfectly reflecting the *ex vivo* assessment, enhancing the predictability of HSC gene therapy, and improving the long-term stability of modified cells at high levels. Thus, CD34^+^CD90^+^ HSCs demonstrate the most refined target population for HSC gene therapy and transplantation.

In order to systematic and objectively compare human candidate HSC-enriched target cell populations for HSC gene therapy, we here combined up-to-date phenotypical, transcriptional, and functional read-outs. We initially analyzed the phenotypical heterogeneity, the relationship in between subsets, as well as the quantitative reduction of target cells for each CD34 subset using flow-cytometry and cell sorting. Next, we purified previously reported HSC-enriched cell populations as well as our recently defined CD34^+^CD90^+^ subset and performed high-throughput scRNAseq analysis in combination with bulk RNAseq. In addition, comprehensive mouse xenograft assays were performed to evaluate the multilineage engraftment and BM reconstitution potential of human CD34^+^ subsets. In this comprehensive side-by-side comparison, we observed the greatest reduction of target cells in the CD34^+^CD90^+^ cell fraction without compromising the multilineage engraftment and BM reconstitution potential in mouse xenograft transplants. Furthermore, scRNAseq data showed that CD34^+^CD90^+^ cells are almost entirely depleted for lineage-committed progenitor cells, while phenotypically, transcriptionally, and functionally primitive HSPCs are markedly enriched. Most importantly for the clinical translation, we compared two GMP-compatible FACS platforms for the sort-purification and gene-modification of HSC-enriched CD34 subsets. Purity and yield of sorting, gene-modification efficiency, as well as maintenance of stem cell features was confirmed by flow-cytometry, with *in vitro* assays, and mouse xenograft transplants. We demonstrate that direct transduction of sort-purified CD34^+^CD90^+^ cells significantly reduced gene-modifying reagents and, at the same time, enhanced the gene-modification efficiency without compromising the long-term multilineage engraftment potential.

These findings are all in line with our recently performed competitive reconstitution studies in the NHP to identify a refined target cell population for HSC gene therapy and transplantation (19). In our previous study, we reported the CD34^+^CD90^+^ phenotype to be exclusively responsible for both rapid short-term as well as robust long-term hematopoietic reconstitution. Here we show that human multilineage engraftment potential with full recovery of the BM stem cell compartment is similarly restricted to CD34^+^CD90^+^ HSPCs, whereas CD34^+^ cells that lost CD90 expression lack full multilineage xenograft reconstitution potential. This finding is also consistent with previous publications describing the enrichment of human HSCs within CD90-expressing CD34 subsets. The first CD90-mediated isolation of human HSCs from fetal BM, adult BM, and cytokine-mobilized stem cells was reported in the 1990s (11, 40). Thereafter, lin^−^CD34^+^CD90^+^ subsets were shown to be enriched for long-term culture-initiating cells (LTC-ICs) with thymus engraftment potential in the SCID mouse model (11, 40). However, alternative CD90/Thy-1 antibody clones (F15 421-5 and GM201) were used at this time that recognized a significantly higher expressed CD90 variant on up to 60% of cytokine-mobilized CD34^+^ cells (11). Refining the phenotype with additional cell surface markers, switching to the currently used CD90 clone 5E10, and using an improved mouse model, Majeti *et al*. later confirmed the enrichment of human HSCs in CD34^+^CD38^−^CD90^+^CD45RA^−^ cell fractions when using UCB- and BM-derived CD34 subsets (27). With the highest level of purification, Notta *et al*. demonstrated engraftment and reconstitution potential of single human CD34^+^CD38^−^CD90^+/–^CD45RA^−^CD49f^+^ cells from UCB after intrafemoral injection into NSG mice (1). Despite significant differences in the phenotype, cell source, level of HSC purification, mouse model (NOD/SCID or NSG), and mode of transplantation (intravenous, intrafemoral), all groups including our studies associate CD90 expression with human HSCs, whereas HSPCs lacking CD90 demonstrate limited engraftment potential and do not re-express or recover the HSC-enriched CD90^+^ subset. Adding to this, we re-validated findings from the 1990s using the CD90 antibody clone 5E10 showing that CD34^+^CD90^+^ HSPCs from GCSF-mobilized leukapheresis products, the most commonly used stem cell source for HSC gene therapy and transplantation, exclusively contain primitive HSCs with multilineage engraftment and BM reconstitution potential.

Recent studies aiming to refine the target for HSC gene therapy and transplantation combined flow-cytometry with functional *in vitro* and/or *in vivo* read-outs (16, 17). Here, we additionally applied single-cell as well as bulk RNA sequencing to comprehensively compare the CD34^+^CD90^+^ phenotype with alternative HSC-enrichment strategies. In contrast to most approaches performing transcriptomics (41–43), we initially performed scRNAseq on CD34^+^ cells to build a detailed and comprehensive reference map rather than using historically defined human CD34 subsets for comparison. This strategy allowed us to define transcriptionally distinct clusters and perform an unbiased assessment of the expression profile in these clusters. Furthermore, we were able to determine the reproducibility of scRNAseq for CD34^+^ HSPCs and ultimately establish a baseline for the comparison of phenotypically defined HSC-enriched cell fractions. Having this baseline further enabled us to objectively compare scRNAseq data from different donors as well as reliably map sort-purified CD34 subsets. Most importantly, this strategy provided the foundation to overlay scRNAseq with bulk RNAseq data from multiple donors without any computational correction or manipulation of the dataset for donor-, batch-, source- or preparation-dependent variability.

Interestingly, we were not able to reliably determine transcriptional differences between CD34^+^CD90^+^ and CD34^+^CD38^low/–^ HSPCs by scRNAseq. Using the 10x scRNAseq V2 platform, we were unable to determine transcriptionally-distinct and meaningful clusters or define more detailed hierarchical structures within either subset. Similar to previous reports on Lin^−^ CD34^+^CD38^low/–^ cells describing a Continuum of LOw-primed UnDifferentiated (CLOUD) cells without hierarchical structures (41–44), CD34^+^CD90^+^ and CD34^+^CD38^low/–^ HSPCs were indistinguishable even though CD38^low/–^ cells were displaying clear differences in their phenotypical composition. Platform-dependent limitations in the mRNA capturing efficiency (31), technical noise (45), and low-level expression of key genes currently limit the ability to reliably distinguish differences in RNA expression (42) required to identify cell subsets with short- and long-term engraftment potential within the CD34^+^CD90^+^ phenotype.

Despite the current limitation in scRNAseq, our approach clearly shows that purified CD34^+^CD90^+^ as well as CD34^+^CD38^low/–^ HSPCs are similarly depleted for transcriptionally lineage-committed progenitor cells. However, combining phenotypical, quantitative, and transcriptional data, the CD34^+^CD90^+^ subset was determined to have the greatest overall reduction of target cells. With an average 12.5-fold reduction in total cells compared to a 5.8-fold for the CD34^+^CD38^low/–^ subset, isolation and targeting of CD34^+^CD90^+^ HSPCs significantly increases the feasibility of currently existing HSC gene therapy approaches. Most importantly, transplantation of purified CD34^+^CD90^+^ HSPCs did not compromise the multilineage engraftment potential and BM reconstitution capacity of CD34 subsets from GCSF-mobilized leukapheresis products.

Particularly important for the application of HSC gene therapy as a routine treatment option, we have successfully demonstrated that the large-scale sort-purification of the HSC-enriched CD90^+^ subset for human stem cell sources is technically feasible and significantly reduces the target cell number as well as costs for HSC gene therapy in comparison to reported small molecule mediated approaches where bulk CD34^+^ cells are targeted and no reduction of modifying agents or improvement of feasibility would be achieved. CD34^+^CD90^+^ cells can be efficiently purified using either the Sony droplet-cell sorter which is clean-room dependent or the fully-closed, portable MACSQuant Tyto from Miltenyi without any significant differences in the purity and yield of cells. However, more comprehensive follow-up studies will be required to further investigate the underlying biological mechanism impacted by both cell-sorting strategies causing the shown differences in multilineage engraftment potential as well as the efficiency of gene-modification. Specific analysis of stress-mediated responses as well as activation of cell cycle pathways may help to explain the observed variances in the frequency of gene-modification and long-term engraftment seen comparing both systems.

Most surprisingly, we found that the sort-purification CD34^+^CD90^+^ cells significantly increased the transduction efficiency of HSCs with multilineage long-term engraftment potential compared to gold standard CD34^+^ cells. In particular, the observation that the gene-modification efficiencies determined in culture directly translate into the frequency of gene-modification *in vivo* will help to increase consistency of quality in stem cell products and ultimately make HSC gene therapy applications more predictable. These findings should have important implications for currently available as well as future HSC gene therapy and gene editing protocols hence only a single change in procedures is require for the purification of CD34^+^CD90^+^ cells. Isolation of this HSC-enriched phenotype will allow more targeted gene modification, allow a more controlled vector copy number without the application of small molecule transduction enhancer, and thus likely reduce unwanted off target effects.

Application of this HSC-enriched subset will require the implementation of additional processing steps into existing clinical protocols. While the sort-purification will generate further expenses, costs performing these steps will be easily compensated by the 30- to 40-fold savings in expensive modifying reagents. Gene therapy vectors are currently estimated to be the biggest hurdle in terms of large-scale production and by far the dominating cost factor limiting the routine application of HSC gene therapy (7–9). Our HSC-targeted gene therapy strategy will help to overcome both bottlenecks at the same time: a single large-scale virus production can be used for up to 30 patients preventing the shortage of GMP-grade vectors; at the same time, modifying reagents will no longer be the major factor for the price determination in HSC gene therapy. Availability of new sorting technology for GMP-grade closed-system cell sorting of human HSCs (46) in combination with our HSC-enriched target population will bring HSC gene therapy a significant step closer towards a feasible routing application.

Of special interest for the clinical routine, we have seen a remarkable consistency in gene-marking of CD90^+^ cells before transplantation, in *ex vivo* culture, and long-term in the mouse xenograft model. Commonly observed donor-to-donor variability in the CD34 population leading to unpredictable levels of transduction *in vivo* were not seen for sort-purified and gene-modified CD90^+^ cells. Especially the drop of *ex vivo* gene-marking onto a significantly lower lever *in vivo* previously observed in human and NHPs (4, 10) was not evident in our experiments. These findings not only provide additional evidence that CD90^+^ HSPCs are uniquely required for robust long-term engraftment of gene-modified cells at high levels but further provides a rapid quality control before transplantation to predict the long-term success for gene modification strategies.

In summary, this study describes a human HSC-enriched cell population with unique phenotypical, transcriptional, and functional features. Isolation of these HSC-enriched CD34^+^CD90^+^ HSPCs has the potential to improve the targeting efficiency of current clinical HSC gene therapy and editing applications, and also reduce toxicity from potential off-target modifications. Conservation of this HSC-enriched phenotype among different human stem cell sources and in the pre-clinical NHP model (19) further highlights its biological relevance. Species-independent similarities of engraftment patterns in mouse xenograft (human) and autologous NHP transplants (19, 27), as well as robust long-term multilineage engraftment with more than two years of follow-up in the NHP (19) are promising indicators for a successful clinical translation of this HSC-enrichment strategy.

## MATERIALS AND METHODS

### Cell sources and CD34^+^ enrichment

Fresh, whole BM in sodium heparin for single cell RNA sequencing was purchased from StemExpress (Folsom, CA). GCSF-mobilized leukapheresis collections were purchased from the Co-operative Center for Excellence in Hematology (CCEH) at the Fred Hutchinson Cancer Research Center. All human samples were obtained after informed consent according to the Declaration of Helsinki stating that the protocols used in this study have been approved by a local ethics committee/institutional review board of the Fred Hutchinson Cancer Research Center. Human CD34^+^ cells from steady-state BM and GCSF-mobilized leukapheresis collections were harvested and enriched as previously described (19, 36). Enrichment of CD34^+^ cell fractions was performed according to the manufacturer’s instructions (Miltenyi Biotech, Bergisch Gladbach, Germany).

### Flow cytometry analysis and FACS

Fluorochrome-conjugated antibodies used for flow cytometric analysis and FACS of human cells are listed in **Supplemental Table 10**. Dead cells and debris were excluded via FSC/SSC gating. Flow cytometric analysis was performed on an LSR IIu (BD, Franklin Lakes, NJ), Fortessa X50 (BD), and FACSAria IIu (BD). Cells for scRNAseq, bulk RNAseq, and *in vitro* assays were sorted using a FACSAria IIu cell sorter (BD). Large-scale clinical-grade purification of CD34^+^CD90^+^ cells was performed on the FX500 (Sony Biotechnology, San Jose, CA) and the MACSQuant Tyto (Miltenyi Biotech). Post-sort purity was assessed on the FACSAria IIu reanalyzing at least 500 cells for each sample. Data was acquired using FACSDiva^TM^ Version 6.1.3 and newer (BD). Data analysis was performed using FlowJo Version 8 and higher (BD).

### RNA isolation for bulk RNAseq

Total RNA from sort-purified CD34 subsets of GCSF-mobilized leukapheresis products was extracted with the Arcturus PicoPure RNA Isolation Kit (Thermo Fisher Scientific, Waltham, MA) according to the manufacturer’s protocol. Total RNA from sort-purified CD34 subsets of steady-state BM was extracted with the RNeasy Micro Kit (Qiagen, Hilden, Germany) according to the manufacturer’s protocol. Detailed methods on the data analysis can be found in the Supplemental Methods.

### RNA quality control for bulk RNAseq

Total RNA integrity was analyzed using an Agilent 2200 TapeStation (Agilent Technologies, Inc., Santa Clara, CA) and quantified using a Trinean DropSense96 spectrophotometer (Caliper Life Sciences, Hopkinton, MA).

### Single cell RNA sequencing

Steady-state BM-derived CD34^+^ cells and CD34-subsets for single-cell RNA sequencing were sort-purified and processed using the Chromium Single Cell 3ʹ (v2) platform from 10X Genomics (Pleasanton, CA). Separation of single cells, RNA extraction, and library preparation were performed in accordance with the 10X Chromium Single Cell Gene Expression Solution protocol. Detailed methods on the data analysis can be found in the Supplemental Methods.

### Next-generation sequencing

Sequencing of GCSF-mobilized and steady-state BM bulk RNAseq samples was performed using Illumina (San Diego, CA) HiSeq 2500 in rapid mode employing a paired-end, 50-base read length (PE50) sequencing strategy. Sequencing of single cell RNAseq samples was performed using an Illumina HiSeq 2500 in rapid mode employing 26 base read length for read 1 (10x barcode and 10bp Unique Molecular Index (UMI)) and 98 base read length for read 2 (cDNA sequence). Image analysis and base calling was performed using Illumina’s Real Time Analysis (v1.18) software, followed by ’demultiplexing’ of indexed reads and generation of fastq files, using Illumina’s bcl2fastq Conversion Software (v1.8.4).

### Lentiviral vectors

The vector used in this study (pRSCSFFV.P140K.PGK.eGFP-sW) is a SIN LV vector produced with a third-generation split packaging system and pseudo-typed by the vesicular stomatitis virus G protein (VSV.G). The vector for these studies was produced by our institutional Vector Production Core (PI H.-P.K.). Infectious titer was determined by flow cytometry evaluating eGFP protein expression following titrated transduction of HT1080 human fibrosarcoma-derived cells with research-grade LV vector preparations.

### Colony-forming cell (CFC) assay

For CFC assays, 1,000-1,200 sort-purified CD34-subpopulations were seeded into MethoCult H4435 (Stemcell Technologies) or H4230 supplemented with hIL-3, IL-5, G-CSF, SCF, TPO, and GM-SCF [all PeproTech], each at 100ug/mL as well as EPO [Amgen, Thousand Oaks, CA, USA] at 4U/mL for the large-scale clinical-grade experiments according to our established clinical protocols (37, 47). Colonies were scored after 12 to 14 days, discriminating colony forming unit-(CFU-) granulocyte (CFU-G), macrophage (CFU-M), granulocyte-macrophage (CFU-GM) and burst forming unit-erythrocyte (BFU-E). Colonies consisting of erythroid and myeloid cells were scored as CFU-MIX.

### Measurement of transduction efficiency and vector copy number (VCN)

To measure the transduction efficiency and vector copy number, at least 80 colonies were picked for each condition. Genomic DNA was isolated by incubating tubes at 95°C for 2 h on a thermal cycler. Crude DNA preparations were then subjected to PCR using LV-specific primers (Fwd: 5’ AGAGATGGGTGCGAGAGCGTCA and Rev: 5’-TGCCTTGGTGGGTGCTACTCCTAA [Integrated DNA Technologies; IDT, Coralville, IA, USA]) and, in a separate reaction, actin-specific primers were used (human Fwd: 5’ TCCTGTGGCACTCACGAAACT and Rev: 5’-GAAGCATTTGCGGTGGACGAT [IDT]). Colonies containing expected bands for both lentivirus and actin were scored as transduced. Reactions which did not yield actin products were considered non-evaluable.

Vector copy number (VCN) per genome equivalent was determined as previously described (38). Briefly, VCNs were assessed by a multiplex TaqMan 5’ nuclease quantitative real-time PCR assay in triplicate reactions. Colony gDNA samples were subjected to a lentivirus-specific primer/probe combination (Fwd, 5’-TGAAAGCGAAAGGGAAACCA; Rev, 5’-CCGTGCGCGCTTCAG; probe, 5’-AGCTCTCTCGACGCAGGACTCGGC [IDT]) as well as an endogenous control (TaqMan Copy Number Reference assay RNaseP, Thermo Fisher Scientific, Pittsburgh, PA, USA) using TaqMan GTXpress Master Mix (Applied Biosystems Foster City, CA). Samples with an average VCN ≥0.5 were considered transduced.

### Mouse xenograft transplantation

Adult (8 to 12 weeks) NSG mice (NOD.Cg-*Prkdc^scid^Il2rg^tm1Wjl^*/SzJ) received a radiation dose of 275 cGy followed 4 hours later by a 200 µL intravenous injection of sort-purified human CD34^+^ cells or CD34-subpopulations. Beginning at 8 weeks post-injection, blood samples were collected every 2 to 4 weeks and analysed by flow cytometry. After 16 to 20 weeks, animals were sacrificed, and tissues were harvested and analysed. All animal studies were carried out at Fred Hutchinson Cancer Research Center in compliance with the approved Institutional Animal Care and Use Committee (IACUC) protocol #1483.

### Statistics

Data analysis of limiting dilution experiments was performed as previously described (48). Statistical analysis of data was performed using GraphPad Prism Version 5. Significance analyses were performed with the unpaired, two-sided Student’s t-test (*: p < 0.05; **: p < 0.01; ***: p <0.001).

## Supporting information

Supplemental Data incl. suppl. methods, tables, and figures

## AUTHOR CONTRIBUTIONS

SR and HPK designed the study. SR, YYC and AMP performed RNAseq. DP and ME analyzed RNAseq data. SR, YYC, AMP, and MC performed longitudinal mouse follow-up, necropsy and data analysis. SR, AMP, YYC, MC, SS, AB and TE performed clinical-grade sorting experiments, follow-up experiments, and data analysis. SR, DP and ME generated the figures. HPK and JEA funded the study. SR and HPK wrote the manuscript. All authors reviewed and edited the final manuscript.

## ACKNOWLEDGEMENTS

Human PB stem cells and steady-state BM were kindly provided by Shelly Heimfeld. We thank Helen Crawford for help in preparing this manuscript and figures.

## DATA AND MATERIALS AVAILABILITY

All original RNAseq data were uploaded to the NCBI database. The BioProject Accession code is available upon request.

