## Supplemental Data incl. suppl. methods, tables, and figures for "Sort-purification of human CD34^+^CD90^+^ cells reduces target cell population and improves lentiviral transduction for gene therapy"

**LIST OF SUPPLEMENTAL ITEMS:**

1. Supplemental Methods
2. Supplemental Table 1. Experimental parameters for single cell RNA sequencing
3. Supplemental Table 2. Differentially expressed genes in Cluster 1, XHS25, CD34$^+$ cells
4. Supplemental Table 3. Top 200 differentially expressed genes in ssBM bulk RNAseq samples.
5. Supplemental Table 4. Top 200 differential genes in GCSF-mobilized bulk RNAseq samples.
6. Supplemental Table 5. Genes upregulated in ssBM cells from Population *a*
7. Supplemental Table 6. Genes upregulated in ssBM cells from Population *b*
8. Supplemental Table 7. Genes upregulated in GCSF-mobilized cells from Population *a*
9. Supplemental Table 8. Genes upregulated in GCSF-mobilized cells from Population *b*
10. Supplemental Table 9. Summary of mobilization, leukapheresis and CD34 enrichment parameters
11. Supplemental Table 10. Antibodies
12. Supplemental Figure 1. Quality control of sort-purified CD34-subpopulations for scRNAseq
13. Supplemental Figure 2. Transcriptionally distinct ssBM CD34 clusters in a second donor
14. Supplemental Figure 3. A scRNAseq reference map of ssBM-derived CD34$^+$ HSPCs
15. Supplemental Figure 4. Donor-independent reproducibility of the scRNAseq ssBM reference map
16. Supplemental Figure 5. Transcriptional mapping of sort-purified CD34 subsets from a second donor
17. Supplemental Figure 6. Quality control of sort-purified CD34-subpopulations for bulk RNAseq
18. Supplemental Figure 7. Differentially expressed genes in GCSF-mobilized bulk CD34 subsets

19. Supplemental Figure 8. Multilineage engraftment potential of human CD34 subpopulations
20. Supplemental Figure 9. Engraftment potential of human CD34 subsets
21. Supplemental Figure 10. Engraftment potential of human HSPCs from Population *b*
22. Supplemental Figure 11. Engraftment potential of gene-modified human bulk $CD34^+$ and sort-purified $CD34^+CD90^+$ cells.

### SUPPLEMENTAL METHODS:

***Expression analysis for bulk RNAseq.*** RNAseq expression analysis was performed in shared resources at the Fred Hutchinson Cancer Research Center. RNAseq libraries of GCSF-mobilized CD34 subsets were prepared using the NuGEN Ovation SoLo RNAseq System (Tecan Genomics, Redwood City, CA, USA). RNAseq libraries of steady-state BM CD34 subsets were prepared using the SMART-Seq v4 Ultra Low Input RNA Kit (Takara Bio Inc., Kusatsu, Japan) and Nextera XT Index Kit v2 (Illumina, Inc., San Diego, CA, USA). Work was performed on a Sciclone NGSx Workstation (PerkinElmer, Waltham, MA, USA). Library size distribution was validated using an Agilent 2200 TapeStation (Agilent Technologies, Santa Clara, CA, USA). Additional library QC, blending of pooled indexed libraries, and cluster optimization was performed using Life Technologies Invitrogen Qubit® 2.0 Fluorometer (Life Technologies-Invitrogen, Carlsbad, CA, USA). RNAseq libraries were pooled and clustered onto a flow cell lane.

***Quantification of transcripts.*** The quantification was performed using kallisto (v0.43.1) (49). Human genome assembly (GRCh38) from National Center for Biotechnology Information (NCBI) was used as the reference. The compressed fastq files (.fastq.gz) were input to kallisto. The human reference transcriptome was processed to create a transcriptome index using "kallisto index " option with the default k-mer length. The abundances of the transcripts were quantified by aligning the raw reads to the reference with bootstrapping, using the option "kallisto quant -b 100". The bootstrapping was performed to obtain confidence intervals on transcript quantification. Kallisto generated two output files with the alignment information. The abundances.tsv reported the abundances as estimated counts (est_counts) and transcripts per million (tpm), while the abundances.h5 file had the abundance estimates, bootstrap estimates, transcript length information, and the run information.

***Data analysis.*** The counts (abundances.tsv) from kallisto were imported into R in the form of a matrix with the tximport package (v.1.10.1). The Human RefSeq Reference Genome Annotation file (v.38_p12), was downloaded from the Human Genome Resources at NCBI to obtain the gene IDs. Each transcript ID and its count was then associated with the corresponding gene ID for summarization of gene-level counts. The count matrix was analyzed for differential gene expression using the DESeq2 package from Bioconductor in R (v.1.22.2).(50) The count matrix was pre-filtered by keeping the rows that have a minimum of one transcript before analysis with DESeq2. The result obtained was a list of differentially expressed genes with significant p-values and log-fold changes. Clustering and principal component analysis (PCA) was performed on the normalized data, which identified the genes that were contributing to the variance in the samples.

***Alignment and counting.*** The 10X Genomics Cell Ranger software suite (v2.0.0) was used to covert the raw sequence reads into single-cell gene expression counts. The "cellranger count" command with default option was run for alignment, filtering, cell barcode counting, and UMI counting. Cell barcode is a known nucleotide sequence that acts as a unique identifier for a single GEM (Gelbead-in-Emulsion) droplet. Each barcode contains reads from a single cell. UMIs are random 10bp nucleotide sequences that help determine which reads came from the same transcript. The cDNA was aligned to human reference genome (hg38) using the STAR aligner (v.2.6.1). UMIs were also filtered for a minimum of Qual = 10. Reads were marked as PCR duplicates if two or more read pairs shared the same cell barcode, UMI, and gene ID. Valid cell barcodes were determined based on the final UMI distributions. Valid cell barcodes with a valid UMI mapped to exons (Ensembl GTF GRCh38) were used to generate the final cell barcode matrix (.mtx).

***Dimensional reduction and clustering.*** The single cell data analysis was performed using Seurat (v2.3.4) (23), an R toolkit for single cell genomic data. The 10X runs for the $CD34^{+}$ cells and the CD34-subsets were merged by combining the cell barcode matrices into a single Seurat object. The gene expression data for each cell was log normalized. The genes were regressed based on the number of UMIs (nUMI), then scaled and centered to improve downstream analysis. PCA was run on the highly variable genes to compute linear dimensional reduction. The cells were clustered based on similar gene expression patterns using the first 10 principal components (PC) with a resolution of 0.4. t-distributed Stochastic Neighbor Embedding (tSNE) was used to visualize the gene clusters and the $CD34^{+}$ cells and CD34-subsets. The positively differentially expressed genes were found for all the clusters based on the Wilcoxon rank sum test with a log-fold change threshold of 0.25. The gene expression patterns of marker genes were visualized on a tSNE dimensional reduction plot and a PCA dimensional reduction plot.

***Single cell and bulk RNAseq combined analysis.*** DESeq2 (v.1.22.2) was run on the bulk RNA data as described above to create an un-normalized count matrix. The raw counts were transformed into a Single Cell Experiment (SCE) object along with the corresponding donor and gene information. The SCE is an R package that includes methods to store single cell data information. The raw counts were used to compute the normalized counts and log counts, which are necessary to convert the SCE data object in to a Seurat data object. Using the Seurat package, the bulk RNAseq data was converted from an SCE object to a Seurat object. UMI counts were generated for the bulk RNA data and added as metadata to the object. Next, the bulk RNAseq data was merged with the single cell RNAseq data to create a combined Seurat dataset. The combined dataset was then log normalized and scaled as described above. This maintained uniformity in the scaling and normalization of both the single cell RNA and bulk RNA data together.

***Transforming the data with significant principal components.*** The 10X run for the $CD34^+$ cells was also analyzed by Seurat (v.2.3.4). The data was normalized and scaled. Variable genes were identified for the data and PCA was run on the variable genes. The genes were clustered using the first 10 PCs with a resolution of 0.4. The genes that defined PC1 and PC2 were extracted from the Seurat object.

A matrix was created by sub-setting the scaled count data matrix of the combined dataset using the PC1 and PC2 genes from the $CD34^+$ data. This matrix was then multiplied with PC1 and PC2 values. The combined single cell and bulk RNA data was thus linearly transformed with the $CD34^+$ cells as the reference and was used for further downstream analysis.

***Overlaying the cell populations on the reference $CD34^+$ cell population.*** Points specific for each of the different cell types from the bulk RNA data, the $CD34^+$ cell population and the CD34 subset cell population were extracted from the combined dataset. PCA was used as the linear dimensional transform. The $CD34^+$ population was plotted as the reference, and the cell types from the bulk RNA data were overlaid on the reference to see where they map. Similarly, the CD34 subsets were visualized against the $CD34^+$ reference map.

***Software and packages.***
FlowJo v.10.2 and higher https://www.flowjo.com
Kallisto v.0.43.1 - https://pachterlab.github.io/kallisto
DESeq2 v.1.22.2 - http://www.bioconductor.org/packages/release/bioc/html/DESeq2.html
Tximport v.1.10.1 - http://bioconductor.org/packages/release/bioc/html/tximport.html
10X Genomics Chromium- https://www.10xgenomics.com/product-list/#single-cell
10X Genomics Cell Ranger v.2.0.0 - https://support.10xgenomics.com/single-cell-gene-expression/software/overview/welcome
STAR aligner v.2.6.1 - https://github.com/alexdobin/STAR
Seurat v.2.3.4 - https://satijalab.org/seurat/

SCE v.1.4.1 -
https://www.bioconductor.org/packages/release/bioc/html/SingleCellExperiment.html

**Supplemental Table 1.** Experimental parameters for scRNAseq

| Donor | 1 | | | | 2 | |
|---|---|---|---|---|---|---|
| **Population** | **CD34$^{+}$** | **CD133$^{+}$** | **CD38$^{low/-}$** | **CD90$^{+}$** | **CD34$^{+}$** | **CD90$^{+}$** |
| # of cells | 2,162 | 2,019 | 2,472 | 1,523 | 1,449 | 1,189 |
| Mean reads/cell | 75,692 | 71,796 | 64,918 | 75,971 | 62,234 | 64,282 |
| Sequencing saturation | 76.6% | 76.9% | 81.2% | 81.7% | 75.2% | 79.1% |
| Fraction reads in cells | 94.7% | 94.6% | 96.7% | 96.4% | 90.5% | 94.5% |
| Valid barcodes | 97.7% | 98.0% | 97.6% | 97.9% | 98.4% | 98.4% |
| Total genes detected | 18,132 | 18,036 | 18,150 | 17,430 | 17,183 | 16,321 |
| Q30 bases in barcodes | 97.4% | 96.8% | 97.4% | 97.0% | 96.2% | 96.2% |
| Q30 bases in RNA reads | 91.3% | 87.2% | 87.7% | 86.0% | 73.5% | 73.6% |
| Q30 bases in sample index | 96.3% | 96.3% | 96.5% | 96.2% | 96.2% | 95.0% |
| Q30 bases in UMI | 97.5% | 96.8% | 97.4% | 97.0% | 96.3% | 96.3% |

**Supplemental Table 2.** Differentially expressed genes in CD34+ cells in Clusters 1 through 7

| Gene | p_val | avg_logFC | pct.1 | pct.2 | p_val_adj |
|---|---|---|---|---|---|
| **Cluster1** | | | | | |
| AVP | 1.03E-115 | 2.353924651 | 0.981 | 0.502 | 3.49E-111 |
| FTH1 | 1.71E-83 | 1.085854554 | 1 | 0.999 | 5.75E-79 |
| HLA-DQB1 | 2.49E-55 | 0.975850333 | 0.711 | 0.299 | 8.40E-51 |
| IDS | 4.68E-57 | 0.891481775 | 0.829 | 0.473 | 1.58E-52 |
| HLA-E | 4.26E-67 | 0.809263775 | 0.995 | 0.875 | 1.44E-62 |
| DUSP1 | 3.91E-28 | 0.76632821 | 0.654 | 0.386 | 1.32E-23 |
| BST2 | 8.26E-52 | 0.752152954 | 0.891 | 0.648 | 2.78E-47 |
| RP11-386I14.4 | 4.26E-22 | 0.751011429 | 0.768 | 0.646 | 1.43E-17 |
| VIM | 9.70E-31 | 0.741547331 | 0.967 | 0.927 | 3.27E-26 |
| MAFF | 6.49E-49 | 0.739097989 | 0.559 | 0.185 | 2.19E-44 |
| FOS | 8.89E-25 | 0.735488763 | 0.744 | 0.526 | 2.99E-20 |
| BEX1 | 2.47E-28 | 0.713080752 | 0.588 | 0.292 | 8.32E-24 |
| TCOF1 | 5.53E-23 | 0.69756564 | 0.526 | 0.299 | 1.86E-18 |
| CD37 | 1.30E-47 | 0.645982327 | 0.943 | 0.791 | 4.39E-43 |
| NAMPT | 1.40E-26 | 0.643196574 | 0.526 | 0.264 | 4.72E-22 |
| HLA-DRB1 | 1.88E-20 | 0.635065174 | 0.782 | 0.695 | 6.33E-16 |
| LST1 | 3.56E-28 | 0.622670938 | 0.801 | 0.596 | 1.20E-23 |
| EIF3E | 7.65E-64 | 0.613628537 | 1 | 0.976 | 2.58E-59 |
| HOPX | 1.76E-43 | 0.613337243 | 0.692 | 0.282 | 5.95E-39 |
| HES1 | 1.30E-32 | 0.588868581 | 0.275 | 0.054 | 4.37E-28 |
| JUN | 5.06E-24 | 0.577589983 | 0.706 | 0.46 | 1.70E-19 |
| LMNA | 1.76E-13 | 0.573224391 | 0.63 | 0.459 | 5.93E-09 |
| MLLT3 | 3.92E-25 | 0.558557455 | 0.668 | 0.439 | 1.32E-20 |
| CEBPB | 1.32E-25 | 0.551536095 | 0.431 | 0.178 | 4.46E-21 |
| SOCS2 | 2.60E-30 | 0.546971417 | 0.592 | 0.308 | 8.75E-26 |
| ICAM3 | 1.01E-26 | 0.538519031 | 0.848 | 0.717 | 3.41E-22 |
| GNA15 | 1.81E-27 | 0.537884801 | 0.834 | 0.676 | 6.11E-23 |
| HLA-DPA1 | 8.59E-29 | 0.534735506 | 0.929 | 0.727 | 2.89E-24 |
| MEG3 | 2.02E-101 | 0.531563048 | 0.322 | 0.012 | 6.81E-97 |
| CD52 | 6.99E-28 | 0.52799503 | 0.791 | 0.512 | 2.36E-23 |
| EIF4A2 | 5.96E-26 | 0.518842472 | 0.886 | 0.779 | 2.01E-21 |
| FOSB | 7.52E-18 | 0.513311832 | 0.488 | 0.273 | 2.53E-13 |
| HLA-DPB1 | 3.47E-25 | 0.508814929 | 0.915 | 0.734 | 1.17E-20 |
| ALDH1A1 | 6.12E-18 | 0.507369148 | 0.611 | 0.429 | 2.06E-13 |
| PPP1CB | 2.10E-15 | 0.499101171 | 0.706 | 0.591 | 7.08E-11 |
| LRRFIP1 | 5.44E-27 | 0.496793361 | 0.882 | 0.72 | 1.83E-22 |
| CD74 | 5.48E-31 | 0.486985875 | 0.986 | 0.971 | 1.85E-26 |
| ID2 | 1.97E-10 | 0.486850753 | 0.502 | 0.355 | 6.63E-06 |
| NEAT1 | 4.55E-19 | 0.479820832 | 0.929 | 0.836 | 1.53E-14 |
| CRHBP | 5.30E-44 | 0.465456032 | 0.384 | 0.08 | 1.79E-39 |
| HLA-DRA | 9.08E-29 | 0.46407168 | 0.976 | 0.897 | 3.06E-24 |
| COMMD6 | 6.01E-36 | 0.461057784 | 0.972 | 0.964 | 2.03E-31 |
| ID1 | 4.67E-26 | 0.460080186 | 0.313 | 0.088 | 1.57E-21 |
| GADD45A | 1.60E-10 | 0.454692376 | 0.479 | 0.356 | 5.40E-06 |
| TSC22D3 | 7.07E-14 | 0.454170716 | 0.716 | 0.582 | 2.38E-09 |
| ZFAS1 | 3.84E-43 | 0.452325707 | 1 | 0.989 | 1.29E-38 |
| PIM1 | 4.34E-19 | 0.4503663 | 0.616 | 0.381 | 1.46E-14 |
| PABPC1 | 1.82E-43 | 0.449636534 | 1 | 0.995 | 6.12E-39 |
| CTD-3252C9.4 | 1.06E-19 | 0.447752438 | 0.251 | 0.074 | 3.57E-15 |
| ANKRD28 | 6.22E-23 | 0.442273671 | 0.953 | 0.921 | 2.09E-18 |
| HSD17B11 | 3.97E-17 | 0.436022043 | 0.806 | 0.732 | 1.34E-12 |

**Supplemental Table 2.** Differentially expressed genes in $CD34^{+}$ cells in Clusters 1 through 7

| Gene | p_val | avg_logFC | pct.1 | pct.2 | p_val_adj |
|---|---|---|---|---|---|
| ZFP36L2 | 2.13E-12 | 0.413234851 | 0.72 | 0.6 | 7.17E-08 |
| MT-ND2 | 8.99E-41 | 0.409917186 | 1 | 0.999 | 3.03E-36 |
| MALAT1 | 3.75E-54 | 0.405582722 | 1 | 1 | 1.26E-49 |
| DDX5 | 1.28E-34 | 0.402617525 | 1 | 0.985 | 4.30E-30 |
| ZNF331 | 9.37E-19 | 0.402477611 | 0.284 | 0.098 | 3.16E-14 |
| CIRBP | 6.08E-38 | 0.400230174 | 1 | 0.991 | 2.05E-33 |
| RP1-313I6.12 | 6.52E-16 | 0.397016139 | 0.408 | 0.207 | 2.20E-11 |
| MDK | 3.57E-15 | 0.393164594 | 0.512 | 0.303 | 1.20E-10 |
| RNF125 | 4.09E-16 | 0.384635603 | 0.45 | 0.25 | 1.38E-11 |
| PTPRC | 4.14E-15 | 0.383886265 | 0.654 | 0.507 | 1.40E-10 |
| SQSTM1 | 2.83E-09 | 0.381792328 | 0.607 | 0.52 | 9.55E-05 |
| PNRC1 | 1.34E-18 | 0.381150979 | 0.953 | 0.891 | 4.52E-14 |
| PCDH9 | 5.34E-28 | 0.38082192 | 0.332 | 0.092 | 1.80E-23 |
| FXYD5 | 9.14E-23 | 0.37906437 | 0.967 | 0.946 | 3.08E-18 |
| YBX3 | 4.48E-13 | 0.377647661 | 0.853 | 0.803 | 1.51E-08 |
| TCF4 | 3.66E-12 | 0.377538437 | 0.526 | 0.37 | 1.23E-07 |
| RBM23 | 1.90E-13 | 0.374783295 | 0.488 | 0.324 | 6.39E-09 |
| EIF1 | 1.09E-45 | 0.374707464 | 1 | 0.999 | 3.67E-41 |
| FOSL2 | 4.44E-15 | 0.372805176 | 0.365 | 0.175 | 1.49E-10 |
| H1F0 | 3.32E-10 | 0.369789578 | 0.607 | 0.476 | 1.12E-05 |
| N4BP2L2 | 3.78E-11 | 0.369105821 | 0.592 | 0.495 | 1.27E-06 |
| EIF2S3 | 1.54E-12 | 0.36691505 | 0.777 | 0.729 | 5.19E-08 |
| HMGA2 | 2.15E-13 | 0.363950558 | 0.403 | 0.231 | 7.25E-09 |
| SNHG7 | 1.41E-15 | 0.363676934 | 0.773 | 0.706 | 4.74E-11 |
| SYPL1 | 3.81E-13 | 0.363520942 | 0.659 | 0.522 | 1.29E-08 |
| HLA-DMA | 5.12E-11 | 0.359864819 | 0.711 | 0.637 | 1.73E-06 |
| PPP1R15A | 2.02E-11 | 0.35161848 | 0.815 | 0.765 | 6.82E-07 |
| CLU | 2.00E-10 | 0.351204638 | 0.327 | 0.178 | 6.75E-06 |
| RSL1D1 | 9.52E-19 | 0.349092663 | 0.905 | 0.849 | 3.21E-14 |
| AJ006998.2 | 5.52E-37 | 0.347009663 | 0.232 | 0.032 | 1.86E-32 |
| ARPC5L | 5.45E-13 | 0.346977233 | 0.73 | 0.635 | 1.84E-08 |
| GLTSCR2 | 9.72E-29 | 0.345468299 | 0.986 | 0.986 | 3.28E-24 |
| TFPI | 1.36E-10 | 0.344888346 | 0.559 | 0.446 | 4.59E-06 |
| BEX2 | 3.12E-11 | 0.344463118 | 0.626 | 0.515 | 1.05E-06 |
| DNAJB6 | 8.36E-16 | 0.341681698 | 0.877 | 0.872 | 2.82E-11 |
| HLA-DQA1 | 1.16E-11 | 0.338184958 | 0.393 | 0.224 | 3.91E-07 |
| TCEAL2 | 2.02E-62 | 0.336363608 | 0.194 | 0.007 | 6.81E-58 |
| SELM | 4.15E-48 | 0.336112804 | 0.209 | 0.016 | 1.40E-43 |
| RELB | 1.28E-08 | 0.334808699 | 0.436 | 0.317 | 0.00043288 |
| LAPTM4A | 6.25E-14 | 0.33185925 | 0.905 | 0.868 | 2.11E-09 |
| EEF2 | 1.28E-33 | 0.33120199 | 1 | 0.995 | 4.32E-29 |
| EIF1B | 8.22E-12 | 0.330575053 | 0.81 | 0.789 | 2.77E-07 |
| NBL1 | 2.31E-10 | 0.32990587 | 0.251 | 0.119 | 7.79E-06 |
| APP | 3.67E-11 | 0.329056417 | 0.422 | 0.261 | 1.24E-06 |
| LAPTM5 | 4.04E-19 | 0.326375524 | 0.943 | 0.907 | 1.36E-14 |
| BEX5 | 1.36E-16 | 0.324942338 | 0.303 | 0.118 | 4.59E-12 |
| EIF5 | 9.47E-10 | 0.323760585 | 0.687 | 0.629 | 3.19E-05 |
| CCNI | 3.13E-17 | 0.322709514 | 0.957 | 0.955 | 1.06E-12 |
| EVI2B | 1.10E-10 | 0.320522256 | 0.445 | 0.283 | 3.71E-06 |
| KIAA0125 | 3.49E-09 | 0.31280148 | 0.682 | 0.604 | 0.00011764 |
| SOCS3 | 1.32E-14 | 0.309188673 | 0.19 | 0.055 | 4.45E-10 |
| FNIP1 | 8.77E-10 | 0.307659527 | 0.678 | 0.637 | 2.96E-05 |
| TSPYL2 | 3.97E-10 | 0.306051007 | 0.332 | 0.186 | 1.34E-05 |
| INSIG1 | 8.73E-09 | 0.296219245 | 0.284 | 0.155 | 0.00029407 |
| VAMP2 | 1.44E-08 | 0.294509969 | 0.791 | 0.789 | 0.00048642 |

**Supplemental Table 2.** Differentially expressed genes in CD34+ cells in Clusters 1 through 7

| Gene | p_val | avg_logFC | pct.1 | pct.2 | p_val_adj |
|---|---|---|---|---|---|
| IL18 | 5.81E-10 | 0.287963231 | 0.346 | 0.202 | 1.96E-05 |
| TAF1D | 4.71E-10 | 0.287072175 | 0.701 | 0.649 | 1.59E-05 |
| EIF3D | 3.91E-12 | 0.286881519 | 0.938 | 0.879 | 1.32E-07 |
| C6orf48 | 7.44E-16 | 0.285435402 | 0.972 | 0.952 | 2.51E-11 |
| AIF1 | 4.93E-13 | 0.2792119 | 0.957 | 0.89 | 1.66E-08 |
| TPT1 | 7.86E-44 | 0.270713984 | 1 | 1 | 2.65E-39 |
| TNFRSF14 | 5.05E-09 | 0.265251532 | 0.289 | 0.157 | 0.00017006 |
| CISH | 4.51E-12 | 0.264061235 | 0.237 | 0.094 | 1.52E-07 |
| ST13 | 1.32E-11 | 0.258695643 | 0.967 | 0.921 | 4.44E-07 |
| TPTEP1 | 1.54E-08 | 0.258242645 | 0.299 | 0.17 | 0.00051802 |
| UBXN1 | 1.45E-08 | 0.253853365 | 0.73 | 0.703 | 0.00048762 |
| ARMCX1 | 2.97E-09 | 0.250233829 | 0.265 | 0.134 | 0.00010015 |
| **Cluster2** | | | | | |
| ELANE | 2.69E-161 | 2.744038 | 0.948 | 0.457 | 9.07E-157 |
| PRTN3 | 3.37E-180 | 2.657692 | 0.893 | 0.258 | 1.13E-175 |
| LYZ | 1.25E-147 | 2.581613 | 0.921 | 0.39 | 4.21E-143 |
| AZU1 | 3.75E-196 | 2.53456 | 0.966 | 0.329 | 1.26E-191 |
| MPO | 1.83E-176 | 2.39908 | 0.997 | 0.764 | 6.17E-172 |
| CFD | 8.97E-170 | 1.615848 | 0.875 | 0.25 | 3.02E-165 |
| SRGN | 4.84E-136 | 1.600164 | 0.988 | 0.763 | 1.63E-131 |
| CTSG | 2.04E-165 | 1.450522 | 0.768 | 0.139 | 6.87E-161 |
| AREG | 9.25E-75 | 1.238173 | 0.814 | 0.431 | 3.12E-70 |
| CALR | 4.41E-83 | 1.136824 | 0.933 | 0.73 | 1.49E-78 |
| RNASE2 | 3.74E-176 | 1.060139 | 0.686 | 0.079 | 1.26E-171 |
| LGALS1 | 2.97E-64 | 1.057621 | 0.857 | 0.535 | 1.00E-59 |
| CSTA | 2.25E-189 | 1.015599 | 0.567 | 0.028 | 7.59E-185 |
| CLEC11A | 4.05E-89 | 0.953917 | 0.942 | 0.781 | 1.36E-84 |
| TMSB4X | 6.42E-80 | 0.837832 | 1 | 0.989 | 2.16E-75 |
| HSP90B1 | 1.32E-64 | 0.801296 | 0.927 | 0.778 | 4.46E-60 |
| MS4A3 | 1.93E-161 | 0.798535 | 0.607 | 0.053 | 6.49E-157 |
| HCST | 1.42E-62 | 0.771956 | 0.78 | 0.404 | 4.79E-58 |
| CST7 | 2.22E-104 | 0.766323 | 0.497 | 0.074 | 7.47E-100 |
| PRSS57 | 3.71E-99 | 0.760102 | 0.988 | 0.905 | 1.25E-94 |
| MGST1 | 4.77E-60 | 0.741063 | 0.729 | 0.374 | 1.61E-55 |
| ANXA1 | 1.05E-61 | 0.729686 | 0.939 | 0.743 | 3.53E-57 |
| CYBA | 8.49E-83 | 0.728603 | 0.979 | 0.847 | 2.86E-78 |
| C1QTNF4 | 3.50E-49 | 0.719108 | 0.802 | 0.506 | 1.18E-44 |
| IGLL1 | 3.18E-38 | 0.67795 | 0.835 | 0.625 | 1.07E-33 |
| SEC61G | 4.00E-59 | 0.665769 | 0.973 | 0.908 | 1.35E-54 |
| RAB32 | 2.07E-58 | 0.625277 | 0.649 | 0.281 | 6.99E-54 |
| MCL1 | 4.62E-40 | 0.600544 | 0.787 | 0.597 | 1.56E-35 |
| MT-ND4L | 1.14E-46 | 0.600183 | 0.957 | 0.818 | 3.83E-42 |
| HSPB1 | 8.02E-54 | 0.599185 | 0.927 | 0.776 | 2.70E-49 |
| PLEK | 1.15E-54 | 0.586194 | 0.607 | 0.241 | 3.87E-50 |
| CPA3 | 1.57E-55 | 0.563443 | 0.518 | 0.169 | 5.29E-51 |
| CTC-425F1.4 | 1.30E-21 | 0.563306 | 0.207 | 0.057 | 4.38E-17 |
| APLP2 | 9.24E-35 | 0.537109 | 0.784 | 0.587 | 3.11E-30 |
| TNFSF13B | 5.06E-48 | 0.536009 | 0.546 | 0.219 | 1.70E-43 |
| SSR4 | 1.71E-45 | 0.527248 | 0.939 | 0.863 | 5.76E-41 |
| HSPA5 | 2.86E-39 | 0.523157 | 0.918 | 0.77 | 9.65E-35 |
| SNHG25 | 1.11E-42 | 0.513366 | 0.948 | 0.895 | 3.73E-38 |
| RNASE3 | 7.73E-104 | 0.509533 | 0.329 | 0.016 | 2.61E-99 |
| MT-CO2 | 2.03E-78 | 0.500118 | 1 | 1 | 6.84E-74 |
| P4HB | 7.23E-39 | 0.491479 | 0.869 | 0.704 | 2.43E-34 |

**Supplemental Table 2.** Differentially expressed genes in CD34+ cells in Clusters 1 through 7

| Gene | p_val | avg_logFC | pct.1 | pct.2 | p_val_adj |
|---|---|---|---|---|---|
| DBI | 1.39E-26 | 0.489876 | 0.692 | 0.497 | 4.67E-22 |
| JUND | 1.36E-25 | 0.488885 | 0.896 | 0.782 | 4.57E-21 |
| TMEM258 | 2.40E-43 | 0.475689 | 0.945 | 0.87 | 8.10E-39 |
| LCP1 | 1.57E-29 | 0.467256 | 0.683 | 0.459 | 5.29E-25 |
| C4orf48 | 6.60E-31 | 0.464646 | 0.595 | 0.333 | 2.22E-26 |
| NPW | 1.35E-27 | 0.463511 | 0.655 | 0.405 | 4.56E-23 |
| TYROBP | 7.96E-33 | 0.452162 | 0.451 | 0.185 | 2.68E-28 |
| SAT1 | 3.55E-30 | 0.451953 | 0.884 | 0.769 | 1.20E-25 |
| EREG | 6.47E-19 | 0.44688 | 0.561 | 0.372 | 2.18E-14 |
| SDF2L1 | 9.68E-29 | 0.444351 | 0.454 | 0.213 | 3.26E-24 |
| PPIB | 1.54E-29 | 0.439397 | 0.857 | 0.768 | 5.20E-25 |
| ATP5I | 1.74E-38 | 0.43772 | 0.963 | 0.919 | 5.85E-34 |
| MT-ATP6 | 4.48E-49 | 0.43431 | 1 | 0.999 | 1.51E-44 |
| PLAC8 | 4.82E-23 | 0.426721 | 0.86 | 0.738 | 1.62E-18 |
| GSTP1 | 1.80E-39 | 0.41591 | 0.988 | 0.958 | 6.06E-35 |
| ROMO1 | 6.46E-26 | 0.41124 | 0.823 | 0.684 | 2.18E-21 |
| TMSB10 | 7.56E-49 | 0.409791 | 1 | 0.998 | 2.55E-44 |
| FAM101B | 1.06E-29 | 0.399849 | 0.494 | 0.228 | 3.57E-25 |
| ATP6V0B | 3.26E-23 | 0.396569 | 0.771 | 0.64 | 1.10E-18 |
| SERF2 | 9.22E-51 | 0.39246 | 1 | 0.993 | 3.11E-46 |
| MT-ND4 | 6.58E-49 | 0.391914 | 1 | 1 | 2.22E-44 |
| MT-ND5 | 1.33E-35 | 0.389783 | 0.991 | 0.991 | 4.48E-31 |
| COX7B | 1.72E-25 | 0.389771 | 0.927 | 0.867 | 5.80E-21 |
| NDUFA3 | 1.32E-22 | 0.385684 | 0.774 | 0.637 | 4.46E-18 |
| MAP3K8 | 5.03E-20 | 0.382961 | 0.78 | 0.616 | 1.69E-15 |
| C14orf2 | 7.99E-41 | 0.382948 | 0.997 | 0.978 | 2.69E-36 |
| GRN | 2.10E-22 | 0.380456 | 0.555 | 0.354 | 7.07E-18 |
| MT-ND3 | 6.72E-51 | 0.377773 | 1 | 0.999 | 2.26E-46 |
| GPX1 | 7.86E-30 | 0.374621 | 0.939 | 0.923 | 2.65E-25 |
| FTL | 1.20E-41 | 0.373425 | 0.997 | 1 | 4.04E-37 |
| NKG7 | 1.27E-34 | 0.371147 | 0.393 | 0.134 | 4.29E-30 |
| PDIA6 | 1.22E-17 | 0.367405 | 0.738 | 0.641 | 4.10E-13 |
| VAMP8 | 4.51E-22 | 0.363554 | 0.86 | 0.809 | 1.52E-17 |
| POLR2L | 2.23E-20 | 0.362418 | 0.89 | 0.773 | 7.50E-16 |
| UQCR11 | 3.72E-25 | 0.357988 | 0.924 | 0.853 | 1.25E-20 |
| NUCB2 | 1.22E-22 | 0.357183 | 0.915 | 0.875 | 4.12E-18 |
| MT-ND1 | 3.18E-41 | 0.353454 | 1 | 0.999 | 1.07E-36 |
| VAMP5 | 3.96E-14 | 0.35012 | 0.524 | 0.362 | 1.33E-09 |
| SERPINB10 | 3.56E-67 | 0.348605 | 0.22 | 0.011 | 1.20E-62 |
| MYDGF | 1.37E-16 | 0.348426 | 0.665 | 0.544 | 4.63E-12 |
| FKBP2 | 8.98E-22 | 0.348035 | 0.43 | 0.223 | 3.03E-17 |
| RAB31 | 2.12E-30 | 0.346673 | 0.293 | 0.082 | 7.13E-26 |
| PDIA4 | 6.27E-20 | 0.340459 | 0.409 | 0.217 | 2.11E-15 |
| FOSL2 | 1.68E-25 | 0.338962 | 0.39 | 0.158 | 5.64E-21 |
| PHPT1 | 1.21E-14 | 0.3376 | 0.668 | 0.553 | 4.09E-10 |
| SPI1 | 2.44E-24 | 0.332759 | 0.476 | 0.244 | 8.23E-20 |
| ATP5E | 9.11E-33 | 0.330138 | 0.994 | 0.987 | 3.07E-28 |
| CSF3R | 8.33E-20 | 0.329386 | 0.637 | 0.397 | 2.81E-15 |
| UQCRQ | 1.42E-20 | 0.329019 | 0.939 | 0.88 | 4.77E-16 |
| CD302 | 1.73E-18 | 0.326487 | 0.421 | 0.227 | 5.82E-14 |
| SNHG9 | 8.05E-18 | 0.323892 | 0.902 | 0.897 | 2.71E-13 |
| XBP1 | 1.65E-14 | 0.322896 | 0.823 | 0.755 | 5.56E-10 |
| CANX | 5.26E-15 | 0.322416 | 0.787 | 0.699 | 1.77E-10 |
| MT-CO3 | 3.36E-42 | 0.320441 | 1 | 1 | 1.13E-37 |
| NDUFC2 | 7.01E-14 | 0.319152 | 0.811 | 0.754 | 2.36E-09 |

**Supplemental Table 2.** Differentially expressed genes in CD34+ cells in Clusters 1 through 7

| Gene | p_val | avg_logFC | pct.1 | pct.2 | p_val_adj |
|---|---|---|---|---|---|
| FABP5 | 2.48E-13 | 0.316524 | 0.631 | 0.519 | 8.36E-09 |
| TMED10 | 6.51E-16 | 0.315693 | 0.716 | 0.607 | 2.20E-11 |
| RPN1 | 4.28E-12 | 0.312348 | 0.485 | 0.363 | 1.44E-07 |
| SPARC | 1.66E-18 | 0.311369 | 0.415 | 0.214 | 5.58E-14 |
| LAMTOR4 | 1.86E-13 | 0.309527 | 0.805 | 0.761 | 6.28E-09 |
| MIF | 1.41E-24 | 0.308745 | 0.966 | 0.938 | 4.73E-20 |
| TMEM205 | 2.44E-15 | 0.307507 | 0.439 | 0.278 | 8.23E-11 |
| FLNA | 8.76E-15 | 0.305929 | 0.482 | 0.318 | 2.95E-10 |
| NDUFA13 | 6.77E-17 | 0.301454 | 0.823 | 0.715 | 2.28E-12 |
| AIF1 | 2.62E-20 | 0.297127 | 0.976 | 0.882 | 8.84E-16 |
| PLD1 | 5.88E-28 | 0.2958 | 0.28 | 0.084 | 1.98E-23 |
| SEC61B | 2.30E-22 | 0.295512 | 0.93 | 0.889 | 7.75E-18 |
| USMG5 | 4.29E-15 | 0.291731 | 0.89 | 0.847 | 1.45E-10 |
| NDUFB2 | 3.28E-12 | 0.290223 | 0.735 | 0.645 | 1.11E-07 |
| TPP1 | 2.03E-15 | 0.289601 | 0.384 | 0.209 | 6.83E-11 |
| ATOX1 | 3.16E-10 | 0.28749 | 0.604 | 0.515 | 1.06E-05 |
| MYL6 | 1.80E-23 | 0.287302 | 0.982 | 0.969 | 6.08E-19 |
| IQGAP1 | 1.56E-10 | 0.28394 | 0.677 | 0.572 | 5.24E-06 |
| SPCS3 | 6.44E-12 | 0.282456 | 0.555 | 0.43 | 2.17E-07 |
| PPIA | 1.74E-24 | 0.28117 | 0.997 | 0.989 | 5.85E-20 |
| RASGRP2 | 5.00E-11 | 0.279351 | 0.524 | 0.395 | 1.68E-06 |
| NGFRAP1 | 1.19E-11 | 0.278936 | 0.762 | 0.697 | 4.01E-07 |
| MT-CO1 | 1.52E-26 | 0.27842 | 1 | 1 | 5.11E-22 |
| FNDC3B | 4.78E-26 | 0.277402 | 0.28 | 0.088 | 1.61E-21 |
| PTPRE | 4.35E-16 | 0.277239 | 0.372 | 0.189 | 1.47E-11 |
| NDUFS6 | 2.43E-10 | 0.275211 | 0.695 | 0.599 | 8.18E-06 |
| KBTBD11 | 7.28E-27 | 0.274548 | 0.253 | 0.07 | 2.45E-22 |
| CNIH4 | 5.87E-11 | 0.273355 | 0.345 | 0.214 | 1.98E-06 |
| SMIM24 | 9.30E-12 | 0.272184 | 0.765 | 0.611 | 3.14E-07 |
| HMGN2 | 2.56E-27 | 0.270688 | 0.997 | 0.991 | 8.63E-23 |
| IFI27L2 | 1.07E-13 | 0.269049 | 0.442 | 0.273 | 3.61E-09 |
| RNASEH2C | 1.66E-09 | 0.268946 | 0.665 | 0.567 | 5.58E-05 |
| CKLF | 8.24E-12 | 0.267731 | 0.451 | 0.303 | 2.78E-07 |
| FAM45A | 1.06E-09 | 0.266033 | 0.445 | 0.324 | 3.57E-05 |
| NEAT1 | 3.34E-15 | 0.265367 | 0.89 | 0.838 | 1.12E-10 |
| IGFBP7 | 7.39E-14 | 0.263755 | 0.759 | 0.642 | 2.49E-09 |
| PET100 | 1.32E-12 | 0.263593 | 0.851 | 0.815 | 4.45E-08 |
| HLA-DRB1 | 3.51E-13 | 0.257118 | 0.78 | 0.69 | 1.18E-08 |
| NUFIP2 | 6.37E-09 | 0.256966 | 0.439 | 0.331 | 0.000215 |
| HGF | 1.33E-34 | 0.255717 | 0.256 | 0.055 | 4.47E-30 |
| PDIA3 | 1.10E-11 | 0.254455 | 0.814 | 0.761 | 3.69E-07 |
| RETN | 2.88E-46 | 0.253997 | 0.149 | 0.007 | 9.72E-42 |
| TXN | 1.46E-14 | 0.252063 | 0.966 | 0.948 | 4.92E-10 |
| ANAPC11 | 9.99E-11 | 0.252011 | 0.701 | 0.614 | 3.37E-06 |
| RUNX1 | 4.44E-09 | 0.250016 | 0.61 | 0.516 | 0.00015 |
| **Cluster3** | | | | | |
| HBB | 2.96E-83 | 2.787861 | 0.986 | 0.755 | 9.97E-79 |
| CA1 | 1.01E-161 | 1.619034 | 0.835 | 0.165 | 3.41E-157 |
| AHSP | 6.45E-169 | 1.43328 | 0.786 | 0.119 | 2.17E-164 |
| APOC1 | 2.31E-156 | 1.419228 | 0.961 | 0.299 | 7.78E-152 |
| BLVRB | 1.24E-147 | 1.349222 | 0.958 | 0.36 | 4.18E-143 |
| S100A6 | 1.00E-122 | 1.292952 | 0.996 | 0.755 | 3.38E-118 |
| FAM178B | 2.66E-144 | 1.287929 | 0.832 | 0.183 | 8.95E-140 |
| HBD | 9.29E-152 | 1.149007 | 0.779 | 0.127 | 3.13E-147 |

**Supplemental Table 2.** Differentially expressed genes in $CD34^+$ cells in Clusters 1 through 7

| Gene | p_val | avg_logFC | pct.1 | pct.2 | p_val_adj |
|---|---|---|---|---|---|
| ATPIF1 | 1.32E-113 | 1.131086 | 0.993 | 0.755 | 4.44E-109 |
| PRDX2 | 2.62E-95 | 1.046908 | 0.979 | 0.691 | 8.83E-91 |
| APOE | 2.11E-151 | 0.85443 | 0.804 | 0.135 | 7.09E-147 |
| TMEM14C | 2.15E-105 | 0.850809 | 0.982 | 0.54 | 7.25E-101 |
| UROD | 7.02E-111 | 0.827546 | 0.954 | 0.412 | 2.37E-106 |
| S100A4 | 3.79E-88 | 0.825483 | 0.996 | 0.898 | 1.28E-83 |
| TUBB2A | 2.89E-169 | 0.761819 | 0.716 | 0.077 | 9.75E-165 |
| MPC2 | 4.85E-92 | 0.747358 | 0.989 | 0.679 | 1.64E-87 |
| TFR2 | 1.75E-147 | 0.729605 | 0.856 | 0.164 | 5.91E-143 |
| LMNA | 8.20E-69 | 0.719628 | 0.909 | 0.41 | 2.76E-64 |
| PLIN2 | 4.57E-79 | 0.704074 | 0.898 | 0.389 | 1.54E-74 |
| HIST1H4C | 1.24E-30 | 0.693404 | 0.968 | 0.749 | 4.17E-26 |
| CALM2 | 3.46E-78 | 0.667808 | 0.989 | 0.744 | 1.16E-73 |
| KCNH2 | 1.99E-176 | 0.663931 | 0.702 | 0.064 | 6.71E-172 |
| HMGB2 | 1.54E-58 | 0.663117 | 0.996 | 0.866 | 5.20E-54 |
| REXO2 | 8.65E-99 | 0.66154 | 0.849 | 0.257 | 2.91E-94 |
| EPCAM | 3.63E-168 | 0.653653 | 0.691 | 0.068 | 1.22E-163 |
| CKS1B | 3.35E-66 | 0.650566 | 0.779 | 0.299 | 1.13E-61 |
| LINC00152 | 6.35E-73 | 0.634383 | 0.979 | 0.602 | 2.14E-68 |
| PVT1 | 3.48E-134 | 0.628236 | 0.782 | 0.137 | 1.17E-129 |
| FBXO7 | 1.08E-75 | 0.620988 | 0.979 | 0.604 | 3.65E-71 |
| HNRNPAB | 8.81E-73 | 0.594187 | 0.926 | 0.459 | 2.97E-68 |
| TMEM14B | 5.21E-65 | 0.594151 | 0.954 | 0.64 | 1.75E-60 |
| NFKBIA | 5.63E-41 | 0.589784 | 0.919 | 0.631 | 1.90E-36 |
| DUT | 4.76E-47 | 0.588257 | 0.975 | 0.731 | 1.60E-42 |
| CA2 | 3.24E-144 | 0.577725 | 0.618 | 0.061 | 1.09E-139 |
| EMP3 | 3.84E-65 | 0.570029 | 0.982 | 0.677 | 1.30E-60 |
| CNRIP1 | 1.55E-128 | 0.564396 | 0.793 | 0.139 | 5.22E-124 |
| SMIM1 | 1.33E-124 | 0.559092 | 0.765 | 0.135 | 4.47E-120 |
| KIAA0101 | 3.13E-43 | 0.547589 | 0.958 | 0.608 | 1.06E-38 |
| CDK4 | 4.41E-66 | 0.540224 | 0.912 | 0.427 | 1.49E-61 |
| H2AFZ | 1.89E-53 | 0.539341 | 0.996 | 0.872 | 6.37E-49 |
| H1FX | 8.11E-35 | 0.538999 | 0.986 | 0.918 | 2.73E-30 |
| CENPU | 2.53E-63 | 0.537207 | 0.835 | 0.348 | 8.52E-59 |
| SLIRP | 6.22E-60 | 0.535948 | 0.937 | 0.532 | 2.10E-55 |
| ISOC2 | 1.26E-86 | 0.530929 | 0.804 | 0.239 | 4.25E-82 |
| HNRNPA2B1 | 5.09E-56 | 0.524532 | 0.993 | 0.832 | 1.72E-51 |
| MINOS1 | 3.40E-62 | 0.524265 | 0.912 | 0.426 | 1.15E-57 |
| ALDH1A1 | 1.22E-61 | 0.520157 | 0.898 | 0.378 | 4.12E-57 |
| TYMS | 1.47E-39 | 0.51875 | 0.909 | 0.519 | 4.96E-35 |
| SERBP1 | 6.15E-59 | 0.516772 | 0.975 | 0.825 | 2.07E-54 |
| ECH1 | 8.87E-58 | 0.51606 | 0.909 | 0.481 | 2.99E-53 |
| MIR4435-2HG | 9.68E-73 | 0.514029 | 0.863 | 0.313 | 3.26E-68 |
| PA2G4 | 1.66E-61 | 0.513073 | 0.996 | 0.789 | 5.59E-57 |
| CD59 | 2.11E-59 | 0.50653 | 0.832 | 0.344 | 7.12E-55 |
| HSP90AA1 | 8.17E-64 | 0.500186 | 1 | 0.979 | 2.75E-59 |
| MCM7 | 3.48E-48 | 0.498765 | 0.888 | 0.476 | 1.17E-43 |
| NOP58 | 5.56E-46 | 0.496705 | 0.912 | 0.526 | 1.87E-41 |
| CXADR | 5.84E-106 | 0.494734 | 0.67 | 0.115 | 1.97E-101 |
| HMBS | 3.07E-98 | 0.490346 | 0.719 | 0.158 | 1.03E-93 |
| SYNGR1 | 1.15E-41 | 0.489235 | 0.919 | 0.606 | 3.88E-37 |
| TPGS2 | 2.40E-65 | 0.487827 | 0.842 | 0.342 | 8.09E-61 |
| TXNIP | 5.16E-45 | 0.487733 | 0.951 | 0.601 | 1.74E-40 |
| ATP5G1 | 5.07E-49 | 0.486279 | 0.947 | 0.612 | 1.71E-44 |
| CAT | 3.67E-47 | 0.483576 | 0.958 | 0.658 | 1.24E-42 |

**Supplemental Table 2.** Differentially expressed genes in CD34$^{+}$ cells in Clusters 1 through 7

| Gene | p_val | avg_logFC | pct.1 | pct.2 | p_val_adj |
|---|---|---|---|---|---|
| HMGB1 | 2.90E-66 | 0.482383 | 1 | 0.988 | 9.78E-62 |
| PCNA | 5.84E-53 | 0.480507 | 0.751 | 0.274 | 1.97E-48 |
| HMGN5 | 1.67E-61 | 0.480234 | 0.811 | 0.311 | 5.61E-57 |
| MPST | 2.06E-50 | 0.478001 | 0.919 | 0.508 | 6.95E-46 |
| NAA38 | 1.04E-51 | 0.476104 | 0.982 | 0.74 | 3.49E-47 |
| NUDC | 1.35E-53 | 0.475885 | 0.933 | 0.547 | 4.55E-49 |
| SOD1 | 1.38E-52 | 0.472889 | 0.989 | 0.816 | 4.66E-48 |
| MARCKSL1 | 7.06E-48 | 0.472153 | 0.951 | 0.542 | 2.38E-43 |
| CKS2 | 7.04E-39 | 0.471816 | 0.825 | 0.406 | 2.37E-34 |
| PDLIM1 | 6.02E-60 | 0.470259 | 0.898 | 0.365 | 2.03E-55 |
| FKBP4 | 5.01E-63 | 0.466522 | 0.744 | 0.254 | 1.69E-58 |
| HSPD1 | 1.34E-35 | 0.465203 | 0.972 | 0.677 | 4.52E-31 |
| FHL2 | 4.22E-126 | 0.462497 | 0.681 | 0.096 | 1.42E-121 |
| PRKAR2B | 3.04E-86 | 0.460816 | 0.775 | 0.198 | 1.02E-81 |
| MRPL52 | 3.89E-54 | 0.457338 | 0.891 | 0.419 | 1.31E-49 |
| C1QBP | 5.94E-50 | 0.456002 | 0.979 | 0.783 | 2.00E-45 |
| BSG | 1.80E-45 | 0.455517 | 0.979 | 0.646 | 6.05E-41 |
| CCT5 | 2.05E-62 | 0.455347 | 0.832 | 0.318 | 6.91E-58 |
| RANBP1 | 4.23E-46 | 0.455252 | 0.979 | 0.727 | 1.42E-41 |
| HIST2H2AC | 1.30E-09 | 0.454835 | 0.607 | 0.399 | 4.39E-05 |
| GTF2A2 | 2.26E-53 | 0.453877 | 0.916 | 0.457 | 7.61E-49 |
| NDFIP1 | 1.07E-67 | 0.450882 | 0.86 | 0.306 | 3.59E-63 |
| DNAJC9 | 2.61E-56 | 0.450384 | 0.832 | 0.327 | 8.78E-52 |
| TUBB4B | 6.59E-47 | 0.446851 | 0.695 | 0.256 | 2.22E-42 |
| CACYBP | 1.97E-50 | 0.444564 | 0.895 | 0.45 | 6.64E-46 |
| POLR2L | 1.38E-41 | 0.444494 | 0.982 | 0.762 | 4.64E-37 |
| ZFP36L1 | 3.14E-78 | 0.442039 | 0.649 | 0.144 | 1.06E-73 |
| PHB | 7.58E-45 | 0.441981 | 0.912 | 0.539 | 2.55E-40 |
| ITGA2B | 8.46E-70 | 0.441782 | 0.839 | 0.262 | 2.85E-65 |
| ACTB | 9.75E-62 | 0.441321 | 1 | 0.995 | 3.28E-57 |
| PTBP1 | 3.57E-49 | 0.441017 | 0.93 | 0.525 | 1.20E-44 |
| TOMM40 | 1.69E-57 | 0.440772 | 0.789 | 0.305 | 5.69E-53 |
| TRIB2 | 1.83E-101 | 0.439053 | 0.67 | 0.117 | 6.16E-97 |
| NCL | 6.31E-46 | 0.437974 | 0.996 | 0.909 | 2.13E-41 |
| HSPE1 | 1.62E-29 | 0.434049 | 0.888 | 0.541 | 5.45E-25 |
| PPP1R14A | 3.74E-111 | 0.431792 | 0.653 | 0.099 | 1.26E-106 |
| HBA1 | 1.42E-59 | 0.430859 | 0.333 | 0.046 | 4.79E-55 |
| MCM5 | 1.65E-45 | 0.429281 | 0.828 | 0.377 | 5.54E-41 |
| POLR2F | 4.04E-57 | 0.42924 | 0.849 | 0.336 | 1.36E-52 |
| HMGA1 | 2.06E-53 | 0.425192 | 0.996 | 0.931 | 6.93E-49 |
| NME4 | 2.96E-44 | 0.423559 | 0.951 | 0.655 | 9.96E-40 |
| TRMT112 | 6.36E-42 | 0.422347 | 0.916 | 0.551 | 2.14E-37 |
| MAP2K2 | 6.16E-60 | 0.41902 | 0.821 | 0.303 | 2.08E-55 |
| NET1 | 1.40E-58 | 0.418855 | 0.796 | 0.284 | 4.73E-54 |
| MRPS34 | 5.80E-65 | 0.418008 | 0.744 | 0.234 | 1.95E-60 |
| BLVRA | 8.73E-67 | 0.41699 | 0.821 | 0.282 | 2.94E-62 |
| DECR1 | 1.14E-57 | 0.416942 | 0.793 | 0.293 | 3.84E-53 |
| HES6 | 3.41E-121 | 0.416589 | 0.604 | 0.072 | 1.15E-116 |
| CCDC112 | 5.44E-91 | 0.416332 | 0.649 | 0.128 | 1.83E-86 |
| TPM1 | 1.25E-83 | 0.415811 | 0.786 | 0.194 | 4.23E-79 |
| ATP5J2 | 2.27E-41 | 0.415645 | 0.989 | 0.739 | 7.66E-37 |
| GLRX5 | 2.29E-52 | 0.414755 | 0.782 | 0.32 | 7.71E-48 |
| RAN | 2.14E-40 | 0.414183 | 0.986 | 0.755 | 7.21E-36 |
| ATP5G3 | 9.70E-47 | 0.413719 | 0.986 | 0.885 | 3.27E-42 |
| TUBB | 3.49E-19 | 0.413025 | 0.979 | 0.861 | 1.18E-14 |

**Supplemental Table 2.** Differentially expressed genes in $CD34^{+}$ cells in Clusters 1 through 7

| Gene | p_val | avg_logFC | pct.1 | pct.2 | p_val_adj |
|---|---|---|---|---|---|
| PCCB | 3.69E-79 | 0.412448 | 0.716 | 0.18 | 1.24E-74 |
| DNPH1 | 7.81E-47 | 0.409908 | 0.919 | 0.478 | 2.63E-42 |
| CYC1 | 3.51E-42 | 0.40988 | 0.93 | 0.585 | 1.18E-37 |
| STK25 | 1.96E-72 | 0.409299 | 0.775 | 0.229 | 6.59E-68 |
| ERH | 1.68E-43 | 0.409123 | 0.986 | 0.723 | 5.65E-39 |
| TIMM8B | 8.59E-57 | 0.406925 | 0.793 | 0.294 | 2.89E-52 |
| NASP | 9.11E-37 | 0.406255 | 0.947 | 0.625 | 3.07E-32 |
| ANK1 | 2.16E-127 | 0.406025 | 0.568 | 0.057 | 7.27E-123 |
| FXN | 1.88E-72 | 0.403459 | 0.772 | 0.224 | 6.34E-68 |
| EIF4G2 | 8.41E-39 | 0.403018 | 0.986 | 0.771 | 2.83E-34 |
| EIF5A | 1.52E-41 | 0.400981 | 0.996 | 0.871 | 5.12E-37 |
| DAD1 | 1.09E-40 | 0.400129 | 0.916 | 0.498 | 3.67E-36 |
| OAZ1 | 1.11E-49 | 0.399691 | 0.996 | 0.888 | 3.74E-45 |
| SUPT16H | 1.22E-56 | 0.398733 | 0.811 | 0.305 | 4.11E-52 |
| NOP10 | 1.11E-36 | 0.393817 | 0.961 | 0.664 | 3.75E-32 |
| NME1 | 1.71E-37 | 0.393708 | 0.895 | 0.554 | 5.76E-33 |
| MCM3 | 3.17E-71 | 0.39323 | 0.684 | 0.179 | 1.07E-66 |
| FCGRT | 1.22E-39 | 0.393159 | 0.814 | 0.388 | 4.10E-35 |
| CST3 | 6.61E-41 | 0.391757 | 0.958 | 0.681 | 2.23E-36 |
| GAR1 | 1.99E-51 | 0.3907 | 0.765 | 0.288 | 6.71E-47 |
| DYNLRB1 | 1.74E-46 | 0.385982 | 0.912 | 0.438 | 5.85E-42 |
| YBX1 | 4.51E-64 | 0.385742 | 1 | 0.992 | 1.52E-59 |
| SLC39A3 | 7.78E-61 | 0.38552 | 0.775 | 0.253 | 2.62E-56 |
| SNRPD3 | 1.15E-39 | 0.385423 | 0.874 | 0.48 | 3.89E-35 |
| UBAC1 | 8.81E-66 | 0.384608 | 0.632 | 0.165 | 2.97E-61 |
| MRPL12 | 7.51E-58 | 0.38417 | 0.751 | 0.255 | 2.53E-53 |
| HIST1H1D | 2.28E-26 | 0.383972 | 0.698 | 0.346 | 7.67E-22 |
| TIMM13 | 3.24E-35 | 0.383959 | 0.968 | 0.733 | 1.09E-30 |
| CENPF | 3.22E-59 | 0.3818 | 0.604 | 0.161 | 1.08E-54 |
| FUS | 4.09E-39 | 0.38173 | 0.993 | 0.864 | 1.38E-34 |
| HEBP1 | 4.57E-48 | 0.380707 | 0.768 | 0.303 | 1.54E-43 |
| ADCK3 | 2.17E-55 | 0.380449 | 0.8 | 0.298 | 7.31E-51 |
| CD82 | 3.65E-43 | 0.378442 | 0.835 | 0.397 | 1.23E-38 |
| SMIM10 | 1.56E-111 | 0.375975 | 0.611 | 0.084 | 5.26E-107 |
| ITGB1 | 4.68E-44 | 0.373877 | 0.768 | 0.314 | 1.58E-39 |
| SNRPG | 2.55E-35 | 0.373613 | 0.951 | 0.623 | 8.60E-31 |
| CTNNBL1 | 1.03E-52 | 0.372534 | 0.846 | 0.326 | 3.46E-48 |
| POP7 | 1.45E-61 | 0.372211 | 0.726 | 0.223 | 4.87E-57 |
| BOLA3 | 1.13E-63 | 0.372133 | 0.716 | 0.216 | 3.80E-59 |
| RUVBL2 | 7.42E-46 | 0.371848 | 0.853 | 0.383 | 2.50E-41 |
| MYL12A | 9.74E-38 | 0.371259 | 0.958 | 0.604 | 3.28E-33 |
| MPP1 | 6.45E-49 | 0.371002 | 0.825 | 0.35 | 2.17E-44 |
| ATP5B | 3.09E-44 | 0.370725 | 0.989 | 0.873 | 1.04E-39 |
| TRAP1 | 8.05E-67 | 0.370402 | 0.695 | 0.195 | 2.71E-62 |
| PSMC3 | 3.12E-34 | 0.370216 | 0.912 | 0.527 | 1.05E-29 |
| STRA13 | 1.34E-45 | 0.369671 | 0.712 | 0.269 | 4.51E-41 |
| TUBA1C | 1.17E-34 | 0.369294 | 0.853 | 0.446 | 3.94E-30 |
| TUBA1B | 3.85E-24 | 0.369234 | 0.996 | 0.952 | 1.30E-19 |
| FECH | 1.18E-69 | 0.368418 | 0.618 | 0.148 | 3.99E-65 |
| TAF9 | 7.10E-33 | 0.368033 | 0.937 | 0.622 | 2.39E-28 |
| SDCBP | 3.78E-34 | 0.36717 | 0.891 | 0.49 | 1.27E-29 |
| CENPV | 1.66E-50 | 0.365373 | 0.821 | 0.324 | 5.59E-46 |
| MTCH2 | 2.13E-48 | 0.361005 | 0.747 | 0.282 | 7.18E-44 |
| SRM | 8.27E-36 | 0.360497 | 0.849 | 0.442 | 2.79E-31 |
| STMN1 | 1.86E-33 | 0.359375 | 0.996 | 0.853 | 6.26E-29 |

**Supplemental Table 2.** Differentially expressed genes in $CD34^{+}$ cells in Clusters 1 through 7

| Gene | p_val | avg_logFC | pct.1 | pct.2 | p_val_adj |
|---|---|---|---|---|---|
| TPI1 | 3.93E-32 | 0.359139 | 0.982 | 0.858 | 1.32E-27 |
| PPA2 | 3.62E-43 | 0.358923 | 0.688 | 0.259 | 1.22E-38 |
| C11orf31 | 3.31E-35 | 0.358059 | 0.968 | 0.684 | 1.11E-30 |
| PSMA7 | 3.66E-33 | 0.357065 | 0.982 | 0.799 | 1.23E-28 |
| TUFM | 1.61E-35 | 0.356712 | 0.968 | 0.733 | 5.43E-31 |
| EIF4EBP1 | 7.19E-45 | 0.354532 | 0.853 | 0.377 | 2.42E-40 |
| SMS | 1.94E-35 | 0.353201 | 0.888 | 0.503 | 6.55E-31 |
| HSPA8 | 7.71E-20 | 0.353101 | 0.888 | 0.628 | 2.60E-15 |
| GNG5 | 5.43E-35 | 0.351877 | 0.94 | 0.646 | 1.83E-30 |
| NDUFAF3 | 1.47E-54 | 0.349846 | 0.782 | 0.27 | 4.95E-50 |
| PRMT1 | 4.21E-34 | 0.348933 | 0.916 | 0.574 | 1.42E-29 |
| PSMB2 | 1.24E-35 | 0.348297 | 0.877 | 0.471 | 4.16E-31 |
| RBX1 | 2.14E-31 | 0.346806 | 0.916 | 0.575 | 7.20E-27 |
| PSMA4 | 8.97E-32 | 0.346037 | 0.937 | 0.638 | 3.02E-27 |
| PCBP1 | 1.87E-28 | 0.343591 | 0.968 | 0.723 | 6.30E-24 |
| GATA1 | 6.12E-91 | 0.342818 | 0.611 | 0.102 | 2.06E-86 |
| UQCRQ | 3.45E-37 | 0.34256 | 0.996 | 0.873 | 1.16E-32 |
| YWHAB | 4.92E-32 | 0.342363 | 0.937 | 0.596 | 1.66E-27 |
| TSPO | 7.33E-31 | 0.342275 | 0.965 | 0.749 | 2.47E-26 |
| EAPP | 1.45E-31 | 0.341543 | 0.916 | 0.561 | 4.89E-27 |
| PAFAH1B3 | 8.80E-36 | 0.340916 | 0.835 | 0.425 | 2.97E-31 |
| YWHAQ | 7.35E-30 | 0.340623 | 0.958 | 0.743 | 2.48E-25 |
| NFIA | 2.90E-86 | 0.33973 | 0.604 | 0.107 | 9.78E-82 |
| LSM3 | 1.14E-31 | 0.339667 | 0.965 | 0.655 | 3.84E-27 |
| SNRPB | 3.34E-35 | 0.338771 | 0.989 | 0.846 | 1.13E-30 |
| H2AFV | 2.02E-30 | 0.338225 | 0.965 | 0.7 | 6.82E-26 |
| CSF1 | 2.90E-77 | 0.337314 | 0.446 | 0.063 | 9.77E-73 |
| IER5 | 4.23E-55 | 0.337258 | 0.656 | 0.192 | 1.43E-50 |
| POMP | 2.74E-32 | 0.337163 | 0.982 | 0.804 | 9.23E-28 |
| CCNB2 | 4.91E-56 | 0.33661 | 0.47 | 0.104 | 1.65E-51 |
| PKIG | 3.57E-46 | 0.33637 | 0.87 | 0.367 | 1.20E-41 |
| RBBP7 | 2.83E-34 | 0.336285 | 0.881 | 0.475 | 9.54E-30 |
| HNRNPK | 1.42E-36 | 0.333656 | 0.993 | 0.897 | 4.78E-32 |
| CAST | 1.31E-40 | 0.33359 | 0.702 | 0.279 | 4.40E-36 |
| ATP5A1 | 7.27E-37 | 0.333347 | 0.993 | 0.872 | 2.45E-32 |
| XRCC5 | 7.86E-34 | 0.332868 | 0.877 | 0.469 | 2.65E-29 |
| NDUFC2 | 2.55E-34 | 0.33233 | 0.975 | 0.73 | 8.60E-30 |
| SRSF2 | 7.98E-27 | 0.332221 | 0.965 | 0.768 | 2.69E-22 |
| PFN1 | 4.52E-38 | 0.331905 | 1 | 0.96 | 1.52E-33 |
| POLD2 | 5.73E-33 | 0.33182 | 0.825 | 0.423 | 1.93E-28 |
| TMEM141 | 1.65E-43 | 0.331728 | 0.737 | 0.286 | 5.57E-39 |
| DDX39A | 3.39E-33 | 0.331694 | 0.856 | 0.44 | 1.14E-28 |
| APOBEC3C | 5.07E-47 | 0.330981 | 0.663 | 0.217 | 1.71E-42 |
| NMI | 4.03E-46 | 0.330959 | 0.8 | 0.308 | 1.36E-41 |
| ETFA | 2.65E-50 | 0.330852 | 0.73 | 0.248 | 8.94E-46 |
| HSD17B10 | 1.40E-36 | 0.330214 | 0.867 | 0.427 | 4.70E-32 |
| UBE2C | 8.63E-20 | 0.330114 | 0.365 | 0.14 | 2.91E-15 |
| UQCR10 | 1.06E-30 | 0.329954 | 0.937 | 0.636 | 3.57E-26 |
| KPNA2 | 2.95E-29 | 0.327564 | 0.705 | 0.331 | 9.96E-25 |
| UBB | 4.03E-40 | 0.327455 | 0.996 | 0.986 | 1.36E-35 |
| AKR1C3 | 1.25E-48 | 0.326692 | 0.695 | 0.23 | 4.21E-44 |
| HDAC2 | 5.71E-31 | 0.326659 | 0.895 | 0.498 | 1.92E-26 |
| PARVB | 7.10E-41 | 0.326352 | 0.796 | 0.346 | 2.39E-36 |
| MRPL1 | 1.11E-45 | 0.325997 | 0.73 | 0.269 | 3.74E-41 |
| PITHD1 | 1.62E-36 | 0.32568 | 0.86 | 0.449 | 5.47E-32 |

**Supplemental Table 2.** Differentially expressed genes in CD34$^{+}$ cells in Clusters 1 through 7

| Gene | p_val | avg_logFC | pct.1 | pct.2 | p_val_adj |
|---|---|---|---|---|---|
| BCLAF1 | 6.32E-32 | 0.325393 | 0.996 | 0.784 | 2.13E-27 |
| UBL5 | 1.69E-29 | 0.324385 | 0.979 | 0.792 | 5.70E-25 |
| MRPL20 | 1.32E-46 | 0.323634 | 0.751 | 0.284 | 4.44E-42 |
| CD36 | 6.56E-143 | 0.323326 | 0.463 | 0.021 | 2.21E-138 |
| TIMP1 | 9.56E-32 | 0.323128 | 0.853 | 0.453 | 3.22E-27 |
| HNRNPR | 1.29E-30 | 0.323027 | 0.958 | 0.609 | 4.33E-26 |
| SLC25A39 | 1.46E-35 | 0.322812 | 0.842 | 0.398 | 4.92E-31 |
| SSB | 5.67E-28 | 0.322757 | 0.947 | 0.643 | 1.91E-23 |
| CD40LG | 6.26E-98 | 0.32273 | 0.449 | 0.045 | 2.11E-93 |
| GPX4 | 2.24E-33 | 0.32036 | 0.989 | 0.84 | 7.54E-29 |
| HBS1L | 6.11E-46 | 0.320183 | 0.758 | 0.281 | 2.06E-41 |
| UBE2S | 7.72E-54 | 0.319005 | 0.66 | 0.196 | 2.60E-49 |
| CCT6A | 3.84E-29 | 0.318886 | 0.93 | 0.592 | 1.29E-24 |
| MAF1 | 1.53E-27 | 0.318498 | 0.912 | 0.598 | 5.16E-23 |
| ATF7IP2 | 8.41E-11 | 0.318457 | 0.818 | 0.622 | 2.83E-06 |
| CENPW | 3.09E-40 | 0.317993 | 0.614 | 0.219 | 1.04E-35 |
| DTYMK | 2.91E-58 | 0.317726 | 0.639 | 0.178 | 9.82E-54 |
| DSTN | 9.42E-25 | 0.31707 | 0.93 | 0.649 | 3.17E-20 |
| MT2A | 8.32E-30 | 0.316216 | 0.751 | 0.338 | 2.80E-25 |
| DNAJC15 | 2.98E-30 | 0.316213 | 0.832 | 0.43 | 1.00E-25 |
| STOML2 | 3.46E-30 | 0.316072 | 0.954 | 0.622 | 1.17E-25 |
| IMP3 | 6.93E-46 | 0.315808 | 0.688 | 0.242 | 2.34E-41 |
| MAP7 | 1.83E-55 | 0.31526 | 0.604 | 0.161 | 6.16E-51 |
| PARP1 | 5.48E-34 | 0.31518 | 0.821 | 0.411 | 1.85E-29 |
| RP11-354E11.2 | 3.59E-38 | 0.314827 | 0.684 | 0.254 | 1.21E-33 |
| POLR2E | 2.87E-31 | 0.314613 | 0.94 | 0.559 | 9.68E-27 |
| LYAR | 3.17E-89 | 0.314485 | 0.498 | 0.07 | 1.07E-84 |
| TK1 | 3.16E-34 | 0.314384 | 0.502 | 0.17 | 1.06E-29 |
| TFDP1 | 9.59E-39 | 0.314321 | 0.835 | 0.378 | 3.23E-34 |
| CNBP | 2.83E-30 | 0.313579 | 0.979 | 0.777 | 9.53E-26 |
| BIRC5 | 3.53E-35 | 0.313424 | 0.523 | 0.18 | 1.19E-30 |
| NDUFB6 | 7.85E-34 | 0.313171 | 0.793 | 0.373 | 2.64E-29 |
| SELENBP1 | 3.55E-103 | 0.312666 | 0.365 | 0.021 | 1.19E-98 |
| NDUFB9 | 1.97E-33 | 0.311972 | 0.982 | 0.839 | 6.65E-29 |
| TESC | 6.77E-64 | 0.311112 | 0.625 | 0.155 | 2.28E-59 |
| SLC39A8 | 2.40E-39 | 0.310847 | 0.807 | 0.342 | 8.09E-35 |
| MRPL4 | 4.02E-39 | 0.310576 | 0.793 | 0.344 | 1.35E-34 |
| PRPF19 | 1.22E-40 | 0.31016 | 0.779 | 0.315 | 4.12E-36 |
| ODC1 | 4.45E-28 | 0.309869 | 0.909 | 0.559 | 1.50E-23 |
| MRPL51 | 1.02E-33 | 0.309116 | 0.818 | 0.384 | 3.43E-29 |
| FSCN1 | 1.04E-40 | 0.309079 | 0.765 | 0.305 | 3.52E-36 |
| CCT2 | 1.12E-26 | 0.309062 | 0.965 | 0.708 | 3.79E-22 |
| ENY2 | 1.71E-29 | 0.308736 | 0.954 | 0.684 | 5.77E-25 |
| NOLC1 | 9.58E-49 | 0.307961 | 0.635 | 0.198 | 3.23E-44 |
| GTF3C6 | 4.02E-34 | 0.307346 | 0.782 | 0.371 | 1.36E-29 |
| COX5A | 1.56E-30 | 0.307234 | 0.993 | 0.835 | 5.27E-26 |
| SLC39A4 | 5.70E-49 | 0.306696 | 0.632 | 0.2 | 1.92E-44 |
| CCT7 | 1.81E-28 | 0.306459 | 0.898 | 0.547 | 6.10E-24 |
| PRELID1 | 7.52E-25 | 0.30595 | 0.923 | 0.638 | 2.53E-20 |
| CDCA4 | 2.54E-44 | 0.304771 | 0.537 | 0.159 | 8.56E-40 |
| TFRC | 1.00E-31 | 0.304558 | 0.73 | 0.331 | 3.37E-27 |
| PRKDC | 8.06E-33 | 0.304429 | 0.751 | 0.342 | 2.72E-28 |
| MTX1 | 1.59E-41 | 0.304023 | 0.744 | 0.289 | 5.34E-37 |
| SDHA | 6.48E-36 | 0.303479 | 0.782 | 0.342 | 2.18E-31 |

**Supplemental Table 2.** Differentially expressed genes in CD34+ cells in Clusters 1 through 7

| Gene | p_val | avg_logFC | pct.1 | pct.2 | p_val_adj |
|---|---|---|---|---|---|
| GADD45GIP1 | 1.06E-25 | 0.303013 | 0.909 | 0.595 | 3.58E-21 |
| TXN2 | 1.06E-26 | 0.302338 | 0.846 | 0.491 | 3.58E-22 |
| HN1 | 2.43E-29 | 0.302302 | 0.804 | 0.421 | 8.20E-25 |
| PXMP2 | 1.40E-39 | 0.302275 | 0.754 | 0.307 | 4.73E-35 |
| AP2M1 | 1.10E-26 | 0.302206 | 0.958 | 0.732 | 3.70E-22 |
| RALBP1 | 3.39E-38 | 0.302175 | 0.796 | 0.334 | 1.14E-33 |
| CYCS | 2.25E-25 | 0.302047 | 0.895 | 0.551 | 7.58E-21 |
| EBNA1BP2 | 3.20E-44 | 0.301589 | 0.775 | 0.295 | 1.08E-39 |
| SUMO2 | 1.31E-33 | 0.301373 | 0.993 | 0.923 | 4.41E-29 |
| HNRNPD | 6.71E-23 | 0.301244 | 0.937 | 0.608 | 2.26E-18 |
| DYNLL1 | 1.07E-42 | 0.301185 | 0.561 | 0.168 | 3.60E-38 |
| MYL4 | 9.93E-124 | 0.301064 | 0.474 | 0.034 | 3.34E-119 |
| HADH | 1.26E-42 | 0.300451 | 0.744 | 0.285 | 4.25E-38 |
| CALM3 | 5.68E-26 | 0.300095 | 0.916 | 0.593 | 1.91E-21 |
| TMEM109 | 2.30E-42 | 0.29974 | 0.702 | 0.258 | 7.75E-38 |
| GHITM | 5.49E-28 | 0.299393 | 0.863 | 0.489 | 1.85E-23 |
| PDCD4 | 4.07E-26 | 0.299088 | 0.877 | 0.496 | 1.37E-21 |
| ZNF451 | 1.59E-34 | 0.299043 | 0.807 | 0.349 | 5.35E-30 |
| DKC1 | 7.85E-35 | 0.298722 | 0.775 | 0.333 | 2.65E-30 |
| LPCAT3 | 1.40E-70 | 0.29801 | 0.554 | 0.111 | 4.71E-66 |
| PCMT1 | 6.59E-49 | 0.297107 | 0.628 | 0.192 | 2.22E-44 |
| MT-CO3 | 8.31E-50 | 0.297042 | 1 | 1 | 2.80E-45 |
| MCM2 | 4.79E-58 | 0.296994 | 0.558 | 0.137 | 1.61E-53 |
| MPDU1 | 1.99E-50 | 0.296902 | 0.646 | 0.2 | 6.70E-46 |
| VBP1 | 4.66E-43 | 0.296884 | 0.779 | 0.308 | 1.57E-38 |
| SNX5 | 1.05E-35 | 0.296768 | 0.775 | 0.32 | 3.53E-31 |
| MAD2L1 | 2.66E-47 | 0.295969 | 0.596 | 0.183 | 8.95E-43 |
| ACADVL | 4.22E-28 | 0.295812 | 0.863 | 0.509 | 1.42E-23 |
| NOSIP | 4.37E-43 | 0.295796 | 0.751 | 0.281 | 1.47E-38 |
| RAC1 | 4.72E-31 | 0.295758 | 0.982 | 0.824 | 1.59E-26 |
| MRPL37 | 6.39E-34 | 0.29562 | 0.818 | 0.383 | 2.15E-29 |
| SRRM2 | 7.70E-30 | 0.295262 | 0.975 | 0.727 | 2.60E-25 |
| 7-Sep | 6.18E-26 | 0.294928 | 0.968 | 0.72 | 2.08E-21 |
| CISD2 | 4.10E-36 | 0.293931 | 0.74 | 0.308 | 1.38E-31 |
| MRPL41 | 4.49E-33 | 0.293841 | 0.835 | 0.409 | 1.51E-28 |
| C17orf89 | 2.53E-29 | 0.293835 | 0.881 | 0.475 | 8.52E-25 |
| TMBIM6 | 2.27E-26 | 0.29371 | 0.965 | 0.671 | 7.64E-22 |
| FDPS | 1.80E-35 | 0.293495 | 0.789 | 0.342 | 6.08E-31 |
| SRI | 6.39E-24 | 0.292894 | 0.874 | 0.537 | 2.15E-19 |
| CMSS1 | 2.48E-48 | 0.292822 | 0.691 | 0.226 | 8.36E-44 |
| EIF3B | 1.23E-34 | 0.291207 | 0.737 | 0.311 | 4.13E-30 |
| METAP2 | 6.52E-21 | 0.291203 | 0.947 | 0.689 | 2.20E-16 |
| PSMD8 | 1.44E-26 | 0.290456 | 0.923 | 0.598 | 4.86E-22 |
| HDAC7 | 1.43E-46 | 0.290392 | 0.66 | 0.214 | 4.81E-42 |
| TMOD1 | 5.83E-97 | 0.289713 | 0.519 | 0.064 | 1.96E-92 |
| PPIA | 2.27E-35 | 0.289501 | 1 | 0.988 | 7.65E-31 |
| RNF187 | 5.52E-38 | 0.289497 | 0.832 | 0.355 | 1.86E-33 |
| RAB11A | 4.09E-28 | 0.289292 | 0.891 | 0.526 | 1.38E-23 |
| EIF6 | 4.52E-28 | 0.289194 | 0.898 | 0.527 | 1.52E-23 |
| BANF1 | 3.95E-32 | 0.289132 | 0.856 | 0.425 | 1.33E-27 |
| UQCRC1 | 2.39E-27 | 0.288592 | 0.874 | 0.486 | 8.04E-23 |
| PSMC4 | 4.88E-42 | 0.288378 | 0.691 | 0.243 | 1.65E-37 |
| PPM1G | 8.81E-28 | 0.288333 | 0.884 | 0.485 | 2.97E-23 |
| PTGES3 | 8.63E-24 | 0.287759 | 0.961 | 0.76 | 2.91E-19 |
| HBQ1 | 5.15E-99 | 0.287358 | 0.425 | 0.038 | 1.74E-94 |

**Supplemental Table 2.** Differentially expressed genes in $CD34^+$ cells in Clusters 1 through 7

| Gene | p_val | avg_logFC | pct.1 | pct.2 | p_val_adj |
|---|---|---|---|---|---|
| STRADB | 1.49E-52 | 0.286468 | 0.572 | 0.154 | 5.01E-48 |
| PRDX3 | 1.40E-42 | 0.286313 | 0.674 | 0.239 | 4.71E-38 |
| PSMB1 | 3.66E-27 | 0.28621 | 0.982 | 0.868 | 1.23E-22 |
| PSMB6 | 9.73E-26 | 0.285669 | 0.93 | 0.563 | 3.28E-21 |
| CENPM | 1.01E-54 | 0.285644 | 0.512 | 0.124 | 3.41E-50 |
| YIF1B | 1.71E-41 | 0.285061 | 0.761 | 0.283 | 5.75E-37 |
| EXOSC8 | 4.99E-42 | 0.285053 | 0.674 | 0.238 | 1.68E-37 |
| ACSS1 | 2.30E-47 | 0.284898 | 0.604 | 0.185 | 7.75E-43 |
| THOC7 | 1.17E-24 | 0.284 | 0.912 | 0.579 | 3.93E-20 |
| PSMC1 | 2.65E-25 | 0.283963 | 0.919 | 0.546 | 8.94E-21 |
| IP6K2 | 2.02E-24 | 0.283803 | 0.87 | 0.498 | 6.79E-20 |
| HIGD1A | 6.95E-42 | 0.283118 | 0.586 | 0.192 | 2.34E-37 |
| PUF60 | 5.41E-35 | 0.282969 | 0.839 | 0.391 | 1.82E-30 |
| NDUFA6 | 2.48E-26 | 0.282413 | 0.853 | 0.485 | 8.36E-22 |
| SAMM50 | 3.14E-42 | 0.282283 | 0.716 | 0.261 | 1.06E-37 |
| UBE2I | 6.04E-24 | 0.282011 | 0.888 | 0.581 | 2.03E-19 |
| AK2 | 9.97E-31 | 0.281613 | 0.73 | 0.337 | 3.36E-26 |
| CHCHD3 | 2.11E-36 | 0.281374 | 0.768 | 0.315 | 7.11E-32 |
| HSPA9 | 9.78E-28 | 0.281038 | 0.937 | 0.612 | 3.29E-23 |
| SSBP1 | 4.98E-22 | 0.280998 | 0.933 | 0.661 | 1.68E-17 |
| HP1BP3 | 7.33E-26 | 0.280717 | 0.849 | 0.464 | 2.47E-21 |
| DNMT1 | 9.05E-33 | 0.280314 | 0.737 | 0.316 | 3.05E-28 |
| PYCR1 | 6.34E-56 | 0.279733 | 0.554 | 0.139 | 2.14E-51 |
| PDHA1 | 1.15E-52 | 0.27959 | 0.596 | 0.165 | 3.87E-48 |
| GLRX3 | 6.47E-36 | 0.278852 | 0.758 | 0.313 | 2.18E-31 |
| PPP1CC | 5.21E-24 | 0.278402 | 0.874 | 0.542 | 1.76E-19 |
| ATAD3A | 2.59E-46 | 0.278109 | 0.625 | 0.197 | 8.71E-42 |
| VDAC3 | 5.77E-27 | 0.277411 | 0.811 | 0.423 | 1.94E-22 |
| ECHS1 | 1.80E-31 | 0.277272 | 0.768 | 0.362 | 6.08E-27 |
| ZWINT | 3.11E-48 | 0.276754 | 0.505 | 0.135 | 1.05E-43 |
| SLC38A5 | 2.40E-59 | 0.276745 | 0.575 | 0.139 | 8.09E-55 |
| JTB | 5.41E-23 | 0.276597 | 0.982 | 0.739 | 1.82E-18 |
| ATG3 | 2.41E-26 | 0.275706 | 0.775 | 0.38 | 8.13E-22 |
| TTLL12 | 1.44E-83 | 0.275522 | 0.495 | 0.072 | 4.85E-79 |
| PDZD8 | 5.72E-28 | 0.275476 | 0.835 | 0.398 | 1.93E-23 |
| DDX1 | 4.80E-27 | 0.275138 | 0.8 | 0.403 | 1.62E-22 |
| PSMD7 | 1.16E-24 | 0.273427 | 0.919 | 0.628 | 3.90E-20 |
| YARS | 7.02E-46 | 0.272749 | 0.67 | 0.222 | 2.37E-41 |
| UQCC2 | 2.60E-49 | 0.272579 | 0.649 | 0.197 | 8.76E-45 |
| CHCHD2 | 4.43E-30 | 0.272547 | 1 | 0.966 | 1.49E-25 |
| NUSAP1 | 2.96E-33 | 0.271736 | 0.512 | 0.179 | 9.97E-29 |
| GMNN | 1.52E-35 | 0.271484 | 0.632 | 0.231 | 5.12E-31 |
| FUNDC2 | 4.74E-35 | 0.270563 | 0.786 | 0.349 | 1.60E-30 |
| TSPAN4 | 1.18E-50 | 0.270278 | 0.604 | 0.17 | 3.99E-46 |
| PSMD14 | 7.34E-41 | 0.270254 | 0.723 | 0.265 | 2.47E-36 |
| PSMD6 | 4.09E-26 | 0.269661 | 0.825 | 0.427 | 1.38E-21 |
| SSRP1 | 1.38E-25 | 0.269632 | 0.874 | 0.476 | 4.67E-21 |
| VPS37B | 1.48E-36 | 0.269574 | 0.709 | 0.273 | 4.98E-32 |
| LAMTOR1 | 1.02E-23 | 0.269501 | 0.888 | 0.527 | 3.45E-19 |
| IFRD2 | 1.15E-46 | 0.26933 | 0.635 | 0.191 | 3.88E-42 |
| HNRNPU | 6.84E-21 | 0.269277 | 0.993 | 0.775 | 2.31E-16 |
| CTA-392E5.1 | 1.04E-112 | 0.269033 | 0.354 | 0.014 | 3.49E-108 |
| NDUFS6 | 1.67E-21 | 0.268482 | 0.884 | 0.572 | 5.63E-17 |
| UPF3A | 9.60E-27 | 0.268172 | 0.853 | 0.43 | 3.23E-22 |
| GCSH | 3.54E-33 | 0.268165 | 0.751 | 0.328 | 1.19E-28 |

**Supplemental Table 2.** Differentially expressed genes in CD34+ cells in Clusters 1 through 7

| Gene | p_val | avg_logFC | pct.1 | pct.2 | p_val_adj |
|---|---|---|---|---|---|
| ETFB | 1.77E-25 | 0.267917 | 0.891 | 0.524 | 5.97E-21 |
| SLC40A1 | 6.89E-52 | 0.267671 | 0.698 | 0.209 | 2.32E-47 |
| NDUFB11 | 4.90E-26 | 0.267607 | 0.986 | 0.843 | 1.65E-21 |
| MTHFD1 | 1.75E-47 | 0.266891 | 0.621 | 0.189 | 5.89E-43 |
| ANP32E | 5.52E-31 | 0.266287 | 0.698 | 0.301 | 1.86E-26 |
| KIF22 | 3.50E-32 | 0.2658 | 0.667 | 0.274 | 1.18E-27 |
| RFXANK | 1.48E-41 | 0.265764 | 0.737 | 0.272 | 4.99E-37 |
| ARL2 | 6.83E-32 | 0.265027 | 0.761 | 0.334 | 2.30E-27 |
| ARL4A | 1.36E-34 | 0.264925 | 0.667 | 0.253 | 4.57E-30 |
| FAM210B | 4.58E-60 | 0.264836 | 0.561 | 0.132 | 1.54E-55 |
| NDUFAF2 | 1.48E-38 | 0.263234 | 0.684 | 0.251 | 4.99E-34 |
| RPA3 | 1.62E-32 | 0.26274 | 0.663 | 0.27 | 5.45E-28 |
| DCTPP1 | 4.04E-43 | 0.262062 | 0.688 | 0.237 | 1.36E-38 |
| HNRNPH3 | 1.90E-21 | 0.261914 | 0.947 | 0.655 | 6.41E-17 |
| PAICS | 1.74E-26 | 0.261431 | 0.789 | 0.408 | 5.85E-22 |
| SNRPD1 | 1.48E-23 | 0.261392 | 0.982 | 0.809 | 5.00E-19 |
| DHRS11 | 1.42E-59 | 0.26137 | 0.537 | 0.12 | 4.80E-55 |
| SERPINE2 | 2.61E-28 | 0.261331 | 0.782 | 0.371 | 8.80E-24 |
| ACSM3 | 1.48E-37 | 0.260911 | 0.635 | 0.221 | 4.97E-33 |
| PDAP1 | 5.14E-26 | 0.260747 | 0.849 | 0.445 | 1.73E-21 |
| SSX2IP | 2.85E-48 | 0.259624 | 0.575 | 0.16 | 9.61E-44 |
| PSMG1 | 3.76E-43 | 0.25942 | 0.625 | 0.199 | 1.27E-38 |
| ANP32A | 3.12E-25 | 0.25929 | 0.807 | 0.421 | 1.05E-20 |
| NDUFAB1 | 9.22E-26 | 0.259202 | 0.825 | 0.454 | 3.11E-21 |
| CYTL1 | 7.74E-25 | 0.2592 | 0.754 | 0.37 | 2.61E-20 |
| PGAM1 | 2.16E-20 | 0.259086 | 0.793 | 0.461 | 7.28E-16 |
| MRPS15 | 8.62E-25 | 0.258792 | 0.835 | 0.451 | 2.90E-20 |
| FAM133B | 1.77E-18 | 0.258628 | 0.958 | 0.713 | 5.98E-14 |
| TST | 9.86E-83 | 0.258282 | 0.456 | 0.06 | 3.32E-78 |
| PIN4 | 2.47E-42 | 0.258274 | 0.628 | 0.206 | 8.31E-38 |
| LSM4 | 5.78E-20 | 0.258181 | 0.912 | 0.623 | 1.95E-15 |
| TALDO1 | 1.81E-20 | 0.25788 | 0.975 | 0.825 | 6.11E-16 |
| NEDD4L | 7.51E-48 | 0.257765 | 0.54 | 0.142 | 2.53E-43 |
| MDH2 | 2.64E-20 | 0.257748 | 0.958 | 0.729 | 8.91E-16 |
| VCP | 5.14E-30 | 0.257588 | 0.779 | 0.36 | 1.73E-25 |
| NDUFB2 | 5.70E-22 | 0.257561 | 0.94 | 0.616 | 1.92E-17 |
| ATIC | 1.15E-31 | 0.257435 | 0.705 | 0.3 | 3.86E-27 |
| ARL6IP1 | 9.06E-15 | 0.256886 | 0.842 | 0.517 | 3.05E-10 |
| HIST1H1E | 2.49E-20 | 0.256847 | 0.681 | 0.363 | 8.39E-16 |
| SNRPC | 1.15E-19 | 0.256118 | 0.94 | 0.687 | 3.88E-15 |
| MAGOH | 8.83E-26 | 0.255878 | 0.923 | 0.547 | 2.98E-21 |
| SAE1 | 1.69E-39 | 0.255752 | 0.653 | 0.231 | 5.71E-35 |
| LRRC59 | 7.47E-34 | 0.254813 | 0.733 | 0.303 | 2.52E-29 |
| PRPF31 | 7.60E-37 | 0.254502 | 0.684 | 0.258 | 2.56E-32 |
| RUVBL1 | 4.59E-59 | 0.254396 | 0.456 | 0.091 | 1.55E-54 |
| MRPL27 | 6.03E-49 | 0.254207 | 0.596 | 0.17 | 2.03E-44 |
| BBC3 | 3.18E-63 | 0.254088 | 0.411 | 0.064 | 1.07E-58 |
| MGAT4B | 6.04E-65 | 0.253437 | 0.456 | 0.079 | 2.03E-60 |
| SSNA1 | 1.52E-25 | 0.253365 | 0.772 | 0.387 | 5.14E-21 |
| CD320 | 1.56E-39 | 0.25334 | 0.589 | 0.198 | 5.27E-35 |
| SMC2 | 4.96E-41 | 0.253233 | 0.509 | 0.151 | 1.67E-36 |
| PABPC4 | 2.99E-21 | 0.252995 | 0.891 | 0.548 | 1.01E-16 |
| FADS2 | 1.95E-49 | 0.252639 | 0.544 | 0.142 | 6.56E-45 |
| UROS | 3.82E-25 | 0.252515 | 0.796 | 0.414 | 1.29E-20 |
| PSMD4 | 5.63E-24 | 0.25229 | 0.856 | 0.471 | 1.90E-19 |

**Supplemental Table 2.** Differentially expressed genes in CD34+ cells in Clusters 1 through 7

| Gene | p_val | avg_logFC | pct.1 | pct.2 | p_val_adj |
|---|---|---|---|---|---|
| RILP | 5.04E-54 | 0.252068 | 0.572 | 0.145 | 1.70E-49 |
| DHFR | 2.42E-39 | 0.25198 | 0.484 | 0.14 | 8.17E-35 |
| BOP1 | 3.10E-40 | 0.251792 | 0.632 | 0.215 | 1.04E-35 |
| CUTA | 3.18E-20 | 0.251786 | 0.993 | 0.79 | 1.07E-15 |
| HSPBP1 | 6.66E-43 | 0.251677 | 0.596 | 0.189 | 2.24E-38 |
| ORC4 | 4.36E-34 | 0.251181 | 0.681 | 0.267 | 1.47E-29 |
| **Cluster4** | | | | | |
| DNTT | 5.97E-118 | 1.972602 | 0.929 | 0.109 | 2.01E-113 |
| CYGB | 4.16E-190 | 1.588961 | 0.776 | 0.029 | 1.40E-185 |
| JCHAIN | 1.42E-111 | 1.411026 | 0.741 | 0.059 | 4.78E-107 |
| LTB | 3.33E-102 | 1.320222 | 0.659 | 0.052 | 1.12E-97 |
| CD99 | 6.99E-46 | 1.282879 | 1 | 0.857 | 2.36E-41 |
| IGHM | 9.21E-32 | 1.249772 | 0.871 | 0.404 | 3.10E-27 |
| CD79A | 5.81E-120 | 1.145807 | 0.635 | 0.037 | 1.96E-115 |
| ACTG1 | 4.11E-41 | 1.087308 | 1 | 0.996 | 1.39E-36 |
| SOD2 | 1.65E-16 | 0.882305 | 0.788 | 0.634 | 5.57E-12 |
| GLRX | 9.15E-27 | 0.876932 | 0.788 | 0.44 | 3.08E-22 |
| TSC22D3 | 2.83E-23 | 0.821382 | 0.882 | 0.584 | 9.55E-19 |
| RBM38 | 2.97E-23 | 0.703977 | 0.518 | 0.161 | 1.00E-18 |
| PAG1 | 5.26E-32 | 0.696414 | 0.424 | 0.076 | 1.77E-27 |
| DSTN | 4.62E-16 | 0.673597 | 0.847 | 0.679 | 1.56E-11 |
| CD74 | 1.00E-23 | 0.658762 | 1 | 0.972 | 3.38E-19 |
| CORO1A | 2.02E-15 | 0.654411 | 0.788 | 0.521 | 6.82E-11 |
| MSI2 | 4.51E-18 | 0.639279 | 0.835 | 0.615 | 1.52E-13 |
| BAALC | 2.47E-16 | 0.637514 | 0.612 | 0.295 | 8.31E-12 |
| ADA | 4.64E-13 | 0.61167 | 0.565 | 0.307 | 1.56E-08 |
| KLF6 | 1.11E-16 | 0.606846 | 0.741 | 0.43 | 3.74E-12 |
| GYPC | 1.25E-15 | 0.605047 | 0.976 | 0.914 | 4.23E-11 |
| MZB1 | 3.36E-11 | 0.597898 | 0.494 | 0.24 | 1.13E-06 |
| ZFP36L2 | 1.05E-11 | 0.596329 | 0.8 | 0.604 | 3.53E-07 |
| LRRFIP1 | 2.63E-12 | 0.58568 | 0.871 | 0.73 | 8.87E-08 |
| UBE2J1 | 2.19E-14 | 0.584148 | 0.753 | 0.582 | 7.39E-10 |
| SLC43A2 | 5.54E-17 | 0.577836 | 0.4 | 0.117 | 1.87E-12 |
| DDAH2 | 9.74E-13 | 0.572112 | 0.835 | 0.725 | 3.28E-08 |
| HLA-DRA | 1.25E-15 | 0.562159 | 0.953 | 0.903 | 4.22E-11 |
| SASH3 | 2.34E-18 | 0.556039 | 0.435 | 0.13 | 7.87E-14 |
| RABAC1 | 9.44E-12 | 0.555847 | 0.788 | 0.626 | 3.18E-07 |
| CALM1 | 5.53E-12 | 0.544873 | 0.812 | 0.693 | 1.86E-07 |
| CD52 | 1.36E-08 | 0.529952 | 0.729 | 0.532 | 0.000459 |
| MED13L | 8.73E-11 | 0.527819 | 0.529 | 0.297 | 2.94E-06 |
| SH3KBP1 | 2.95E-10 | 0.527572 | 0.565 | 0.348 | 9.95E-06 |
| SORL1 | 1.47E-11 | 0.526219 | 0.518 | 0.257 | 4.95E-07 |
| NEIL1 | 9.63E-60 | 0.515813 | 0.318 | 0.017 | 3.24E-55 |
| LST1 | 2.93E-11 | 0.515719 | 0.8 | 0.608 | 9.87E-07 |
| HMGB1 | 1.45E-18 | 0.50326 | 1 | 0.989 | 4.88E-14 |
| UBC | 9.09E-20 | 0.496596 | 1 | 0.984 | 3.06E-15 |
| CYFIP2 | 3.93E-09 | 0.494818 | 0.412 | 0.198 | 0.000133 |
| VPREB1 | 6.24E-47 | 0.488919 | 0.341 | 0.029 | 2.10E-42 |
| 9-Sep | 2.50E-10 | 0.479332 | 0.529 | 0.288 | 8.41E-06 |
| LAT2 | 3.53E-10 | 0.478548 | 0.518 | 0.282 | 1.19E-05 |
| KMT2E | 1.28E-09 | 0.46599 | 0.882 | 0.818 | 4.30E-05 |
| ARID4B | 2.51E-08 | 0.451769 | 0.812 | 0.758 | 0.000845 |
| SCAI | 1.41E-10 | 0.451668 | 0.4 | 0.174 | 4.75E-06 |
| DDX5 | 1.76E-18 | 0.444069 | 1 | 0.986 | 5.94E-14 |

**Supplemental Table 2.** Differentially expressed genes in CD34+ cells in Clusters 1 through 7

| Gene | p_val | avg_logFC | pct.1 | pct.2 | p_val_adj |
|---|---|---|---|---|---|
| NPY | 1.73E-59 | 0.441628 | 0.247 | 0.009 | 5.83E-55 |
| PTMA | 1.55E-15 | 0.434748 | 1 | 1 | 5.23E-11 |
| ITM2C | 2.87E-09 | 0.42314 | 0.871 | 0.706 | 9.66E-05 |
| HLA-DPB1 | 3.50E-09 | 0.415811 | 0.859 | 0.747 | 0.000118 |
| BAHCC1 | 1.19E-16 | 0.377263 | 0.271 | 0.056 | 4.00E-12 |
| EEF1D | 3.15E-16 | 0.373909 | 1 | 0.997 | 1.06E-11 |
| ARHGAP27 | 3.31E-16 | 0.372062 | 0.247 | 0.048 | 1.12E-11 |
| NEGR1 | 3.26E-12 | 0.360537 | 0.259 | 0.067 | 1.10E-07 |
| TMSB10 | 5.06E-17 | 0.36005 | 1 | 0.999 | 1.70E-12 |
| MALAT1 | 5.85E-18 | 0.353599 | 1 | 1 | 1.97E-13 |
| FAM129C | 4.90E-33 | 0.351868 | 0.247 | 0.022 | 1.65E-28 |
| TRBC2 | 5.23E-09 | 0.346737 | 0.353 | 0.144 | 0.000176 |
| IL7R | 8.84E-62 | 0.336788 | 0.224 | 0.006 | 2.98E-57 |
| LINC00426 | 8.73E-42 | 0.327193 | 0.2 | 0.009 | 2.94E-37 |
| PRKD2 | 2.37E-10 | 0.326741 | 0.282 | 0.09 | 7.97E-06 |
| EBF1 | 5.21E-56 | 0.323432 | 0.224 | 0.007 | 1.75E-51 |
| LL22NC03-2H8.5 | 6.22E-09 | 0.315906 | 0.188 | 0.049 | 0.00021 |
| ARL4C | 2.07E-09 | 0.30724 | 0.212 | 0.056 | 6.98E-05 |
| UMODL1 | 1.06E-36 | 0.288308 | 0.176 | 0.008 | 3.57E-32 |
| RHPN1 | 7.31E-09 | 0.287544 | 0.247 | 0.08 | 0.000246 |
| HLA-C | 1.56E-08 | 0.283269 | 0.965 | 0.978 | 0.000527 |
| GAPDH | 2.51E-11 | 0.279403 | 1 | 1 | 8.46E-07 |
| EIF1 | 1.80E-12 | 0.277735 | 1 | 1 | 6.05E-08 |
| MME | 6.26E-44 | 0.250838 | 0.129 | 0.002 | 2.11E-39 |
| **Cluster5** | | | | | |
| HIST1H4C | 1.36E-52 | 1.386421 | 0.945 | 0.759 | 4.58E-48 |
| TUBA1B | 9.78E-76 | 1.18954 | 0.982 | 0.955 | 3.29E-71 |
| TUBB | 5.35E-75 | 1.044876 | 0.991 | 0.864 | 1.80E-70 |
| JCHAIN | 4.72E-45 | 0.969847 | 0.333 | 0.058 | 1.59E-40 |
| RGS2 | 9.17E-18 | 0.900249 | 0.388 | 0.162 | 3.09E-13 |
| STMN1 | 8.19E-69 | 0.886491 | 0.986 | 0.859 | 2.76E-64 |
| IGHM | 3.72E-40 | 0.856201 | 0.795 | 0.38 | 1.25E-35 |
| LGALS1 | 9.27E-28 | 0.841482 | 0.863 | 0.552 | 3.12E-23 |
| CORO1A | 7.41E-61 | 0.822742 | 0.941 | 0.485 | 2.50E-56 |
| CD74 | 2.15E-34 | 0.811641 | 0.995 | 0.97 | 7.25E-30 |
| KIAA0101 | 2.80E-53 | 0.794664 | 0.922 | 0.624 | 9.45E-49 |
| HMGN2 | 2.31E-68 | 0.781542 | 0.995 | 0.992 | 7.80E-64 |
| H2AFZ | 2.19E-53 | 0.780984 | 0.982 | 0.878 | 7.38E-49 |
| TYMS | 1.76E-45 | 0.768163 | 0.858 | 0.538 | 5.92E-41 |
| HMGB2 | 4.75E-44 | 0.737636 | 0.968 | 0.873 | 1.60E-39 |
| ITM2C | 1.13E-49 | 0.731248 | 0.977 | 0.682 | 3.82E-45 |
| C12orf75 | 2.89E-52 | 0.698178 | 0.63 | 0.194 | 9.73E-48 |
| LSP1 | 3.65E-35 | 0.681364 | 0.849 | 0.534 | 1.23E-30 |
| PCNA | 3.18E-44 | 0.679175 | 0.699 | 0.296 | 1.07E-39 |
| CDKN2D | 2.77E-40 | 0.665784 | 0.712 | 0.295 | 9.35E-36 |
| IGLL1 | 3.13E-29 | 0.656806 | 0.886 | 0.631 | 1.05E-24 |
| PLP2 | 8.30E-35 | 0.628564 | 0.845 | 0.483 | 2.80E-30 |
| HLA-DRA | 5.44E-25 | 0.628311 | 0.982 | 0.896 | 1.83E-20 |
| NUSAP1 | 2.15E-66 | 0.62635 | 0.658 | 0.174 | 7.25E-62 |
| BIRC5 | 7.89E-57 | 0.621847 | 0.626 | 0.18 | 2.66E-52 |
| PLD4 | 1.60E-43 | 0.607894 | 0.479 | 0.125 | 5.38E-39 |
| HLA-DPA1 | 2.26E-29 | 0.601719 | 0.954 | 0.723 | 7.61E-25 |
| HIST2H2AC | 6.29E-25 | 0.596928 | 0.694 | 0.396 | 2.12E-20 |

**Supplemental Table 2.** Differentially expressed genes in $CD34^+$ cells in Clusters 1 through 7

| Gene | p_val | avg_logFC | pct.1 | pct.2 | p_val_adj |
|---|---|---|---|---|---|
| CYBA | 1.55E-33 | 0.574809 | 0.991 | 0.853 | 5.24E-29 |
| HLA-DPB1 | 1.05E-27 | 0.562739 | 0.95 | 0.729 | 3.55E-23 |
| NASP | 8.27E-35 | 0.551236 | 0.881 | 0.643 | 2.79E-30 |
| HERPUD1 | 8.93E-24 | 0.525383 | 0.621 | 0.304 | 3.01E-19 |
| PKM | 2.05E-35 | 0.524407 | 0.904 | 0.566 | 6.92E-31 |
| ATAD2 | 5.75E-43 | 0.515064 | 0.639 | 0.223 | 1.94E-38 |
| VIM | 1.05E-31 | 0.512745 | 1 | 0.923 | 3.53E-27 |
| ID2 | 8.37E-30 | 0.508933 | 0.717 | 0.33 | 2.82E-25 |
| CXCR4 | 1.85E-31 | 0.508182 | 0.493 | 0.167 | 6.23E-27 |
| CFL1 | 8.81E-45 | 0.503857 | 1 | 0.966 | 2.97E-40 |
| UBE2C | 1.04E-58 | 0.503495 | 0.553 | 0.126 | 3.50E-54 |
| TRA2B | 6.68E-22 | 0.499702 | 0.954 | 0.843 | 2.25E-17 |
| IER2 | 4.70E-17 | 0.498872 | 0.708 | 0.44 | 1.58E-12 |
| HLA-DRB1 | 2.05E-20 | 0.497917 | 0.913 | 0.68 | 6.90E-16 |
| TK1 | 3.73E-49 | 0.492602 | 0.584 | 0.172 | 1.26E-44 |
| CENPN | 1.40E-40 | 0.490676 | 0.584 | 0.204 | 4.71E-36 |
| DDIT4 | 7.72E-27 | 0.490023 | 0.525 | 0.215 | 2.60E-22 |
| PLAUR | 9.49E-38 | 0.489924 | 0.393 | 0.094 | 3.20E-33 |
| HELLS | 1.39E-42 | 0.488297 | 0.744 | 0.306 | 4.68E-38 |
| CRIP1 | 8.14E-20 | 0.486534 | 0.338 | 0.119 | 2.74E-15 |
| CDCA5 | 4.58E-80 | 0.485042 | 0.575 | 0.103 | 1.54E-75 |
| SIVA1 | 1.68E-26 | 0.482877 | 0.895 | 0.653 | 5.65E-22 |
| SLC3A2 | 1.92E-27 | 0.480218 | 0.689 | 0.355 | 6.46E-23 |
| CKS1B | 4.38E-34 | 0.478948 | 0.721 | 0.322 | 1.47E-29 |
| CDT1 | 1.35E-26 | 0.478451 | 0.703 | 0.39 | 4.55E-22 |
| EIF4A3 | 9.53E-41 | 0.474345 | 0.639 | 0.24 | 3.21E-36 |
| PHGDH | 6.27E-55 | 0.473654 | 0.616 | 0.17 | 2.11E-50 |
| SPNS3 | 9.18E-41 | 0.469848 | 0.744 | 0.294 | 3.09E-36 |
| HLA-C | 1.14E-30 | 0.466775 | 1 | 0.975 | 3.85E-26 |
| CENPU | 1.39E-31 | 0.461333 | 0.749 | 0.374 | 4.68E-27 |
| DUT | 6.66E-24 | 0.457943 | 0.909 | 0.747 | 2.25E-19 |
| IDH2 | 1.29E-31 | 0.455361 | 0.95 | 0.718 | 4.35E-27 |
| HLA-DRB5 | 4.53E-23 | 0.447679 | 0.831 | 0.486 | 1.53E-18 |
| SKA3 | 5.16E-64 | 0.440813 | 0.516 | 0.102 | 1.74E-59 |
| MKI67 | 8.67E-55 | 0.4393 | 0.484 | 0.103 | 2.92E-50 |
| NUCKS1 | 5.30E-21 | 0.43827 | 0.84 | 0.626 | 1.79E-16 |
| ACTG1 | 6.14E-24 | 0.434258 | 1 | 0.996 | 2.07E-19 |
| GADD45B | 2.96E-36 | 0.429271 | 0.493 | 0.146 | 9.97E-32 |
| MYBL2 | 7.27E-54 | 0.428502 | 0.543 | 0.134 | 2.45E-49 |
| RAB11FIP1 | 1.31E-14 | 0.421314 | 0.406 | 0.197 | 4.41E-10 |
| SRSF10 | 2.10E-29 | 0.413737 | 0.959 | 0.812 | 7.08E-25 |
| GMNN | 3.91E-35 | 0.412206 | 0.612 | 0.247 | 1.32E-30 |
| DBI | 1.20E-23 | 0.411322 | 0.785 | 0.498 | 4.06E-19 |
| SRSF7 | 3.18E-22 | 0.408729 | 0.799 | 0.487 | 1.07E-17 |
| ARPC5L | 1.26E-20 | 0.405226 | 0.858 | 0.62 | 4.24E-16 |
| HMGB1 | 1.09E-26 | 0.40484 | 1 | 0.989 | 3.67E-22 |
| HLA-B | 1.04E-28 | 0.404649 | 1 | 0.995 | 3.49E-24 |
| ATAD5 | 1.12E-43 | 0.404573 | 0.616 | 0.202 | 3.78E-39 |
| ARL4C | 6.82E-37 | 0.395026 | 0.256 | 0.04 | 2.30E-32 |
| DNAJC9 | 4.12E-24 | 0.392765 | 0.712 | 0.358 | 1.39E-19 |
| SLC25A5 | 1.93E-31 | 0.389661 | 0.991 | 0.927 | 6.49E-27 |
| HIST1H1E | 1.60E-18 | 0.389653 | 0.653 | 0.377 | 5.39E-14 |
| PMAIP1 | 2.87E-20 | 0.380509 | 0.594 | 0.297 | 9.67E-16 |
| RAD51AP1 | 1.02E-52 | 0.379656 | 0.543 | 0.135 | 3.45E-48 |
| DEK | 6.75E-23 | 0.379205 | 0.922 | 0.782 | 2.27E-18 |

**Supplemental Table 2.** Differentially expressed genes in CD34+ cells in Clusters 1 through 7

| Gene | p_val | avg_logFC | pct.1 | pct.2 | p_val_adj |
|---|---|---|---|---|---|
| CDCA7 | 2.30E-29 | 0.37774 | 0.626 | 0.267 | 7.76E-25 |
| TMSB10 | 1.14E-11 | 0.371523 | 1 | 0.998 | 3.83E-07 |
| CKS2 | 6.95E-22 | 0.370724 | 0.749 | 0.429 | 2.34E-17 |
| IRF8 | 3.03E-35 | 0.370296 | 0.311 | 0.063 | 1.02E-30 |
| CXXC5 | 4.26E-22 | 0.369455 | 0.653 | 0.335 | 1.44E-17 |
| HLA-DMA | 3.77E-15 | 0.367796 | 0.813 | 0.625 | 1.27E-10 |
| B2M | 1.27E-35 | 0.36625 | 1 | 1 | 4.30E-31 |
| RUNX3 | 5.36E-33 | 0.365272 | 0.443 | 0.131 | 1.80E-28 |
| SMC4 | 1.54E-20 | 0.364293 | 0.557 | 0.268 | 5.19E-16 |
| IGKC | 3.87E-28 | 0.358914 | 0.137 | 0.013 | 1.30E-23 |
| USP1 | 1.14E-22 | 0.358269 | 0.639 | 0.317 | 3.83E-18 |
| ORC6 | 3.75E-49 | 0.35767 | 0.575 | 0.157 | 1.27E-44 |
| ARPC2 | 3.19E-23 | 0.356313 | 0.973 | 0.87 | 1.07E-18 |
| CLSPN | 1.06E-49 | 0.3561 | 0.452 | 0.097 | 3.58E-45 |
| ANAPC11 | 3.69E-20 | 0.353008 | 0.84 | 0.603 | 1.24E-15 |
| TMPO | 7.63E-23 | 0.352259 | 0.575 | 0.267 | 2.57E-18 |
| NPC2 | 2.42E-21 | 0.349824 | 0.941 | 0.728 | 8.14E-17 |
| CDCA4 | 9.03E-36 | 0.349223 | 0.53 | 0.173 | 3.04E-31 |
| RP11-386I14.4 | 8.82E-14 | 0.348874 | 0.84 | 0.637 | 2.97E-09 |
| CD2AP | 1.48E-17 | 0.3483 | 0.557 | 0.276 | 5.00E-13 |
| CAPG | 3.27E-48 | 0.348021 | 0.603 | 0.16 | 1.10E-43 |
| ZFP36L2 | 9.03E-17 | 0.3464 | 0.868 | 0.583 | 3.04E-12 |
| LSM4 | 6.52E-20 | 0.344617 | 0.872 | 0.638 | 2.20E-15 |
| PRC1 | 5.98E-37 | 0.344395 | 0.53 | 0.167 | 2.02E-32 |
| RGS1 | 1.32E-12 | 0.344133 | 0.237 | 0.086 | 4.44E-08 |
| PTPRE | 1.19E-28 | 0.34363 | 0.516 | 0.183 | 4.01E-24 |
| MCM7 | 2.74E-17 | 0.343094 | 0.767 | 0.503 | 9.25E-13 |
| GGH | 2.03E-21 | 0.340763 | 0.603 | 0.325 | 6.85E-17 |
| UBC | 1.05E-12 | 0.34052 | 1 | 0.983 | 3.53E-08 |
| CTSB | 9.21E-12 | 0.338882 | 0.53 | 0.313 | 3.10E-07 |
| TOP2A | 5.48E-50 | 0.337493 | 0.42 | 0.081 | 1.85E-45 |
| LGMN | 1.65E-59 | 0.336406 | 0.237 | 0.016 | 5.57E-55 |
| SMC3 | 1.13E-18 | 0.330599 | 0.817 | 0.571 | 3.82E-14 |
| JUNB | 6.66E-10 | 0.329253 | 0.562 | 0.365 | 2.24E-05 |
| PPP1R18 | 1.49E-20 | 0.328663 | 0.589 | 0.272 | 5.02E-16 |
| LCP1 | 2.34E-21 | 0.325313 | 0.781 | 0.46 | 7.90E-17 |
| MYL6 | 1.10E-19 | 0.324842 | 0.986 | 0.969 | 3.71E-15 |
| SPINK2 | 3.61E-13 | 0.324799 | 0.872 | 0.64 | 1.22E-08 |
| CALM1 | 1.84E-11 | 0.323633 | 0.886 | 0.676 | 6.21E-07 |
| CCDC50 | 9.97E-11 | 0.322769 | 0.338 | 0.168 | 3.36E-06 |
| RFC1 | 1.44E-19 | 0.321735 | 0.721 | 0.42 | 4.85E-15 |
| IRF2BP2 | 4.75E-09 | 0.321466 | 0.799 | 0.647 | 0.00016 |
| TUBB4B | 1.65E-19 | 0.321362 | 0.589 | 0.283 | 5.54E-15 |
| PTTG1 | 1.16E-21 | 0.317353 | 0.635 | 0.326 | 3.89E-17 |
| GSTP1 | 5.05E-21 | 0.312461 | 0.986 | 0.96 | 1.70E-16 |
| PPIG | 3.37E-15 | 0.31244 | 0.9 | 0.722 | 1.14E-10 |
| TAGLN2 | 8.02E-15 | 0.310861 | 0.973 | 0.915 | 2.70E-10 |
| FUS | 1.16E-15 | 0.309903 | 0.973 | 0.871 | 3.89E-11 |
| CENPF | 4.62E-21 | 0.309418 | 0.47 | 0.191 | 1.56E-16 |
| ERP29 | 3.34E-13 | 0.309037 | 0.936 | 0.81 | 1.13E-08 |
| CYTH4 | 8.65E-38 | 0.307125 | 0.288 | 0.05 | 2.92E-33 |
| 9-Sep | 3.48E-13 | 0.306778 | 0.502 | 0.275 | 1.17E-08 |
| PTPRS | 8.79E-17 | 0.305702 | 0.11 | 0.016 | 2.96E-12 |
| NUDT1 | 5.46E-21 | 0.305668 | 0.781 | 0.453 | 1.84E-16 |

**Supplemental Table 2.** Differentially expressed genes in $CD34^+$ cells in Clusters 1 through 7

| Gene | p_val | avg_logFC | pct.1 | pct.2 | p_val_adj |
|---|---|---|---|---|---|
| ATF3 | 6.29E-22 | 0.305044 | 0.32 | 0.095 | 2.12E-17 |
| COX8A | 8.26E-14 | 0.303016 | 0.945 | 0.795 | 2.78E-09 |
| HLA-DQA1 | 5.07E-18 | 0.302141 | 0.484 | 0.213 | 1.71E-13 |
| CENPW | 9.97E-26 | 0.300175 | 0.575 | 0.237 | 3.36E-21 |
| RALY | 4.55E-16 | 0.299629 | 0.872 | 0.644 | 1.53E-11 |
| DDX39A | 1.80E-14 | 0.299406 | 0.735 | 0.467 | 6.08E-10 |
| CENPH | 6.13E-18 | 0.29843 | 0.671 | 0.395 | 2.07E-13 |
| MIS18BP1 | 4.18E-18 | 0.296507 | 0.731 | 0.441 | 1.41E-13 |
| DNAJC1 | 1.50E-11 | 0.296002 | 0.717 | 0.492 | 5.06E-07 |
| CARHSP1 | 2.04E-17 | 0.295327 | 0.639 | 0.358 | 6.87E-13 |
| MAD2L2 | 1.48E-15 | 0.291952 | 0.575 | 0.336 | 5.00E-11 |
| SRSF2 | 7.61E-14 | 0.291742 | 0.941 | 0.777 | 2.56E-09 |
| ALOX5AP | 6.25E-24 | 0.287133 | 0.215 | 0.043 | 2.11E-19 |
| FMNL1 | 8.26E-12 | 0.286594 | 0.68 | 0.451 | 2.78E-07 |
| SAE1 | 2.10E-21 | 0.285531 | 0.571 | 0.254 | 7.06E-17 |
| TMEM106C | 1.44E-19 | 0.285358 | 0.461 | 0.201 | 4.85E-15 |
| ATP6V0B | 1.63E-12 | 0.284139 | 0.831 | 0.641 | 5.49E-08 |
| MAD2L1 | 6.04E-21 | 0.279624 | 0.489 | 0.209 | 2.04E-16 |
| UBALD2 | 3.95E-13 | 0.279247 | 0.694 | 0.426 | 1.33E-08 |
| KIF22 | 1.87E-19 | 0.277998 | 0.594 | 0.295 | 6.29E-15 |
| PSME2 | 4.50E-14 | 0.276675 | 0.826 | 0.609 | 1.52E-09 |
| NAP1L1 | 5.58E-21 | 0.276659 | 1 | 0.989 | 1.88E-16 |
| MZB1 | 2.49E-19 | 0.275754 | 0.516 | 0.22 | 8.39E-15 |
| ADGRE5 | 1.79E-19 | 0.274778 | 0.411 | 0.16 | 6.05E-15 |
| ARHGDIA | 1.37E-12 | 0.269867 | 0.767 | 0.539 | 4.62E-08 |
| ANKRD11 | 3.97E-15 | 0.269822 | 0.658 | 0.377 | 1.34E-10 |
| CH17-373J23.1 | 7.45E-13 | 0.269437 | 0.461 | 0.233 | 2.51E-08 |
| UHRF1 | 6.13E-27 | 0.269067 | 0.379 | 0.114 | 2.07E-22 |
| LDLRAD4 | 1.05E-24 | 0.267583 | 0.283 | 0.072 | 3.55E-20 |
| HIST1H2AM | 5.16E-28 | 0.267369 | 0.37 | 0.109 | 1.74E-23 |
| ARF6 | 2.83E-09 | 0.266209 | 0.872 | 0.657 | 9.54E-05 |
| TACC3 | 1.42E-30 | 0.265894 | 0.438 | 0.135 | 4.79E-26 |
| UBE2J1 | 2.53E-08 | 0.26566 | 0.767 | 0.568 | 0.000852 |
| LMNB1 | 2.36E-22 | 0.265367 | 0.42 | 0.155 | 7.94E-18 |
| AURKB | 1.32E-43 | 0.26416 | 0.361 | 0.069 | 4.46E-39 |
| HMGXB4 | 3.07E-13 | 0.262621 | 0.74 | 0.511 | 1.04E-08 |
| MGST3 | 2.88E-16 | 0.262118 | 0.63 | 0.343 | 9.70E-12 |
| OAZ1 | 1.18E-10 | 0.261437 | 0.977 | 0.893 | 3.96E-06 |
| YEATS4 | 3.43E-16 | 0.261365 | 0.616 | 0.339 | 1.15E-11 |
| PHF5A | 5.01E-18 | 0.261198 | 0.694 | 0.387 | 1.69E-13 |
| ASPM | 1.59E-36 | 0.261107 | 0.338 | 0.072 | 5.36E-32 |
| APP | 1.37E-11 | 0.260746 | 0.493 | 0.252 | 4.61E-07 |
| C9orf142 | 2.49E-17 | 0.260705 | 0.63 | 0.335 | 8.40E-13 |
| BRCA2 | 3.67E-32 | 0.259611 | 0.379 | 0.101 | 1.24E-27 |
| TUBA1A | 7.67E-11 | 0.259363 | 0.932 | 0.778 | 2.58E-06 |
| GGNBP2 | 2.67E-09 | 0.258396 | 0.872 | 0.673 | 8.99E-05 |
| HSPB11 | 3.71E-15 | 0.257323 | 0.667 | 0.382 | 1.25E-10 |
| IKZF1 | 8.24E-12 | 0.25732 | 0.708 | 0.476 | 2.77E-07 |
| VAMP8 | 9.67E-10 | 0.256736 | 0.913 | 0.805 | 3.26E-05 |
| SLC9A3R1 | 1.87E-25 | 0.256449 | 0.397 | 0.127 | 6.29E-21 |
| DDAH2 | 1.78E-09 | 0.256447 | 0.868 | 0.714 | 5.98E-05 |
| DNMT1 | 1.76E-15 | 0.256278 | 0.612 | 0.344 | 5.92E-11 |
| PIM1 | 1.37E-13 | 0.253988 | 0.639 | 0.378 | 4.62E-09 |
| CD99 | 1.90E-16 | 0.251346 | 0.982 | 0.85 | 6.39E-12 |

**Supplemental Table 2.** Differentially expressed genes in $CD34^+$ cells in Clusters 1 through 7

| Gene | p_val | avg_logFC | pct.1 | pct.2 | p_val_adj |
|---|---|---|---|---|---|
| EZR | 1.11E-13 | 0.250955 | 0.662 | 0.402 | 3.73E-09 |
| TACC1 | 2.07E-14 | 0.250616 | 0.708 | 0.426 | 6.96E-10 |
| CDK1 | 4.86E-41 | 0.250182 | 0.365 | 0.075 | 1.64E-36 |
| **Cluster6** | | | | | |
| SPINK2 | 7.49E-141 | 0.875219 | 0.974 | 0.562 | 2.52E-136 |
| SMIM24 | 3.82E-91 | 0.609764 | 0.923 | 0.54 | 1.29E-86 |
| GYPC | 1.65E-75 | 0.584367 | 0.981 | 0.896 | 5.55E-71 |
| CD52 | 3.37E-79 | 0.569203 | 0.84 | 0.441 | 1.14E-74 |
| C1QTNF4 | 3.24E-70 | 0.55467 | 0.844 | 0.455 | 1.09E-65 |
| KIAA0125 | 5.12E-54 | 0.544485 | 0.801 | 0.55 | 1.73E-49 |
| ZFP36L2 | 5.36E-53 | 0.508646 | 0.836 | 0.539 | 1.81E-48 |
| CSF3R | 2.50E-70 | 0.507954 | 0.758 | 0.328 | 8.43E-66 |
| ITM2C | 3.61E-64 | 0.507053 | 0.906 | 0.649 | 1.22E-59 |
| HOPX | 2.66E-67 | 0.491702 | 0.628 | 0.222 | 8.95E-63 |
| ICAM3 | 3.77E-67 | 0.48801 | 0.923 | 0.667 | 1.27E-62 |
| PIK3R1 | 8.69E-50 | 0.43364 | 0.821 | 0.548 | 2.93E-45 |
| IGHM | 8.48E-60 | 0.412511 | 0.718 | 0.326 | 2.86E-55 |
| NPDC1 | 6.90E-50 | 0.39914 | 0.654 | 0.309 | 2.32E-45 |
| ENO1 | 1.01E-54 | 0.393016 | 0.989 | 0.956 | 3.41E-50 |
| MAP3K8 | 1.43E-34 | 0.369228 | 0.818 | 0.583 | 4.83E-30 |
| IGLL1 | 4.17E-25 | 0.366091 | 0.786 | 0.615 | 1.41E-20 |
| EGFL7 | 6.29E-36 | 0.364198 | 0.812 | 0.579 | 2.12E-31 |
| SELL | 1.06E-35 | 0.363842 | 0.534 | 0.26 | 3.59E-31 |
| TAOK3 | 3.44E-39 | 0.360642 | 0.75 | 0.492 | 1.16E-34 |
| PRDX1 | 1.26E-29 | 0.359411 | 0.835 | 0.701 | 4.25E-25 |
| AIF1 | 1.26E-38 | 0.34536 | 0.991 | 0.866 | 4.25E-34 |
| C1orf228 | 5.81E-31 | 0.340551 | 0.786 | 0.588 | 1.96E-26 |
| CD99 | 1.39E-44 | 0.338984 | 0.953 | 0.834 | 4.67E-40 |
| NAP1L1 | 1.65E-58 | 0.337055 | 0.998 | 0.987 | 5.56E-54 |
| MSI2 | 8.20E-31 | 0.319183 | 0.789 | 0.57 | 2.76E-26 |
| LDHA | 2.34E-28 | 0.318657 | 0.951 | 0.882 | 7.90E-24 |
| TKT | 3.29E-29 | 0.311865 | 0.944 | 0.875 | 1.11E-24 |
| SPNS3 | 1.73E-30 | 0.310106 | 0.543 | 0.274 | 5.84E-26 |
| GAPDH | 4.02E-66 | 0.309449 | 1 | 1 | 1.35E-61 |
| PKM | 3.02E-27 | 0.304338 | 0.789 | 0.539 | 1.02E-22 |
| PRSS2 | 5.44E-27 | 0.303514 | 0.141 | 0.023 | 1.83E-22 |
| GNA15 | 6.09E-29 | 0.301522 | 0.814 | 0.651 | 2.05E-24 |
| CD74 | 4.78E-44 | 0.290992 | 1 | 0.964 | 1.61E-39 |
| LST1 | 2.29E-23 | 0.290001 | 0.776 | 0.563 | 7.72E-19 |
| MZB1 | 1.44E-34 | 0.286126 | 0.449 | 0.185 | 4.85E-30 |
| BEX1 | 1.99E-14 | 0.281795 | 0.444 | 0.281 | 6.70E-10 |
| PRAM1 | 7.12E-38 | 0.28024 | 0.346 | 0.111 | 2.40E-33 |
| H2AFY | 4.46E-33 | 0.269617 | 0.97 | 0.919 | 1.50E-28 |
| LSP1 | 6.25E-22 | 0.26946 | 0.712 | 0.518 | 2.11E-17 |
| HLA-DPA1 | 1.04E-29 | 0.269196 | 0.889 | 0.7 | 3.51E-25 |
| CASP4 | 2.38E-20 | 0.261376 | 0.65 | 0.467 | 8.01E-16 |
| IGFBP7 | 6.14E-18 | 0.260128 | 0.746 | 0.632 | 2.07E-13 |
| HLA-A | 1.18E-32 | 0.259081 | 0.996 | 0.99 | 3.97E-28 |
| PIM2 | 1.65E-22 | 0.25249 | 0.393 | 0.194 | 5.56E-18 |
| 6-Sep | 9.44E-19 | 0.251816 | 0.823 | 0.712 | 3.18E-14 |
| **Cluster7** | | | | | |
| RP11-354E11.2 | 7.29E-114 | 0.780843 | 0.687 | 0.197 | 2.45E-109 |
| ITGA2B | 2.89E-86 | 0.771274 | 0.659 | 0.24 | 9.74E-82 |

**Supplemental Table 2.** Differentially expressed genes in CD34$^{+}$ cells in Clusters 1 through 7

| Gene | p_val | avg_logFC | pct.1 | pct.2 | p_val_adj |
|---|---|---|---|---|---|
| PDLIM1 | 8.85E-57 | 0.613631 | 0.693 | 0.358 | 2.98E-52 |
| S100A4 | 1.44E-68 | 0.589988 | 0.972 | 0.892 | 4.85E-64 |
| FCER1A | 6.03E-106 | 0.588952 | 0.494 | 0.083 | 2.03E-101 |
| PDZD8 | 2.05E-65 | 0.588485 | 0.723 | 0.375 | 6.91E-61 |
| GATA2 | 8.83E-59 | 0.535689 | 0.53 | 0.188 | 2.98E-54 |
| SLC40A1 | 1.22E-76 | 0.519833 | 0.572 | 0.183 | 4.12E-72 |
| LINC00152 | 8.95E-46 | 0.510985 | 0.811 | 0.604 | 3.01E-41 |
| PHACTR4 | 3.22E-48 | 0.505008 | 0.687 | 0.386 | 1.08E-43 |
| RNF130 | 2.06E-59 | 0.500831 | 0.89 | 0.664 | 6.95E-55 |
| SOX4 | 2.18E-42 | 0.498087 | 0.928 | 0.795 | 7.36E-38 |
| PKIG | 1.10E-58 | 0.497496 | 0.697 | 0.353 | 3.70E-54 |
| PHTF1 | 2.74E-43 | 0.476195 | 0.627 | 0.336 | 9.23E-39 |
| RP11-620J15.3 | 2.98E-56 | 0.45974 | 0.952 | 0.845 | 1.00E-51 |
| ELF1 | 4.05E-50 | 0.442993 | 0.956 | 0.772 | 1.36E-45 |
| EIF4G2 | 2.99E-45 | 0.439771 | 0.904 | 0.768 | 1.01E-40 |
| MARCKSL1 | 3.17E-41 | 0.416969 | 0.787 | 0.539 | 1.07E-36 |
| S100A6 | 2.80E-51 | 0.410048 | 0.928 | 0.745 | 9.44E-47 |
| CTNNBL1 | 3.47E-41 | 0.402142 | 0.612 | 0.329 | 1.17E-36 |
| TPM1 | 1.28E-46 | 0.401696 | 0.5 | 0.204 | 4.31E-42 |
| PRKACB | 1.21E-37 | 0.394257 | 0.588 | 0.316 | 4.09E-33 |
| ZEB2 | 2.16E-30 | 0.38424 | 0.725 | 0.523 | 7.27E-26 |
| GLUL | 2.48E-16 | 0.37647 | 0.797 | 0.658 | 8.36E-12 |
| FERMT3 | 2.73E-16 | 0.374862 | 0.462 | 0.31 | 9.21E-12 |
| H1F0 | 4.67E-17 | 0.365462 | 0.62 | 0.449 | 1.57E-12 |
| MPP1 | 1.15E-31 | 0.361393 | 0.596 | 0.357 | 3.87E-27 |
| LEPROT | 6.11E-30 | 0.356756 | 0.813 | 0.621 | 2.06E-25 |
| NAA38 | 3.49E-26 | 0.354115 | 0.886 | 0.737 | 1.18E-21 |
| EMP3 | 3.37E-28 | 0.345896 | 0.825 | 0.684 | 1.14E-23 |
| FBXO7 | 5.84E-30 | 0.343371 | 0.813 | 0.605 | 1.97E-25 |
| APOC1 | 3.16E-22 | 0.34231 | 0.562 | 0.333 | 1.07E-17 |
| MPST | 2.59E-21 | 0.334938 | 0.691 | 0.523 | 8.73E-17 |
| PSTPIP2 | 5.48E-34 | 0.334906 | 0.484 | 0.239 | 1.85E-29 |
| ACSM3 | 1.63E-31 | 0.333019 | 0.462 | 0.219 | 5.50E-27 |
| SH3BGRL | 1.58E-32 | 0.323508 | 0.918 | 0.814 | 5.33E-28 |
| CITED2 | 3.11E-23 | 0.322323 | 0.414 | 0.216 | 1.05E-18 |
| SERPINB1 | 1.36E-20 | 0.317732 | 0.936 | 0.893 | 4.59E-16 |
| CAT | 9.31E-24 | 0.316209 | 0.815 | 0.663 | 3.14E-19 |
| ATF7IP2 | 1.68E-08 | 0.315041 | 0.693 | 0.634 | 0.000566 |
| CNRIP1 | 1.22E-37 | 0.312131 | 0.428 | 0.163 | 4.12E-33 |
| CD63 | 8.53E-14 | 0.309657 | 0.865 | 0.781 | 2.87E-09 |
| STK4 | 1.57E-23 | 0.30482 | 0.586 | 0.377 | 5.28E-19 |
| PRKAR2B | 1.43E-27 | 0.302685 | 0.448 | 0.222 | 4.81E-23 |
| MLLT3 | 1.12E-32 | 0.299694 | 0.687 | 0.393 | 3.76E-28 |
| SEC11A | 6.83E-24 | 0.292422 | 0.793 | 0.627 | 2.30E-19 |
| SDCBP | 1.35E-23 | 0.291909 | 0.691 | 0.498 | 4.55E-19 |
| MIR4435-2HG | 4.82E-21 | 0.286074 | 0.55 | 0.336 | 1.63E-16 |
| TLN1 | 5.69E-21 | 0.285944 | 0.506 | 0.312 | 1.92E-16 |
| RHOG | 3.88E-20 | 0.280233 | 0.731 | 0.563 | 1.31E-15 |
| 7-Sep | 2.98E-23 | 0.27823 | 0.873 | 0.716 | 1.00E-18 |
| MPC2 | 4.19E-17 | 0.269064 | 0.811 | 0.693 | 1.41E-12 |
| LIMD2 | 3.19E-22 | 0.268165 | 0.789 | 0.624 | 1.07E-17 |
| MYL12A | 4.54E-20 | 0.266046 | 0.791 | 0.608 | 1.53E-15 |
| VASP | 1.19E-20 | 0.263944 | 0.452 | 0.262 | 4.02E-16 |
| HACD1 | 7.09E-21 | 0.261557 | 0.48 | 0.295 | 2.39E-16 |

**Supplemental Table 2.** Differentially expressed genes in $CD34^{+}$ cells in Clusters 1 through 7

| Gene | p_val | avg_logFC | pct.1 | pct.2 | p_val_adj |
|---|---|---|---|---|---|
| MALAT1 | 6.87E-28 | 0.25731 | 1 | 1 | 2.32E-23 |
| STAT5A | 8.18E-20 | 0.256875 | 0.47 | 0.281 | 2.76E-15 |
| F2R | 4.52E-19 | 0.253278 | 0.424 | 0.237 | 1.52E-14 |

**Supplemental Table 3.** Top 200 differentially expressed genes in ssBM bulk RNAseq samples

| Donor | 1 | 2 | 1 | 2 | 1 | 2 | 1 | 2 | 1 | 2 | 1 | 2 | 1 | 2 |
|---|---|---|---|---|---|---|---|---|---|---|---|---|---|---|
| **Population** | ***a*** | ***a*** | ***b*** | ***b*** | ***c*** | ***c*** | ***d*** | ***d*** | ***e*** | ***e*** | ***f*** | ***f*** | ***g*** | ***g*** |
| **Gene** | | | | | | | | | | | | | | |
| CLC | -2.27951 | -2.39081 | -0.25356 | -1.94062 | -2.26832 | -2.14076 | 7.968692 | 7.702633 | -0.90277 | 0.570894 | -2.56784 | -2.52205 | -0.37181 | 1.395811 |
| PRTN3 | -2.67668 | -2.53473 | 1.586639 | 4.585715 | 3.679756 | 5.720002 | 3.009806 | 3.079114 | -2.77907 | -2.83274 | -2.83728 | -2.80594 | -2.85852 | -2.33608 |
| IRF8 | -2.58528 | -2.28258 | 1.340121 | 1.924075 | 7.20119 | 7.045834 | -1.4114 | -0.88649 | -0.89122 | -1.02461 | -2.70457 | -2.66413 | -1.75697 | -1.30397 |
| ELANE | -2.44605 | -2.72547 | 2.471089 | 3.906982 | 4.041444 | 4.019191 | 4.0078 | 2.861747 | -2.64526 | -2.53964 | -2.51498 | -2.87309 | -2.72463 | -2.83912 |
| KCNH2 | -2.38667 | -2.34451 | -0.17213 | -0.95853 | -2.22415 | -1.89981 | 7.374522 | 6.243694 | -1.43069 | 0.470124 | -2.58595 | -2.53377 | 1.382146 | 1.065722 |
| KCNE5 | -2.11916 | -2.07413 | 1.748818 | 2.248882 | 6.317836 | 6.677164 | -0.95698 | -0.79768 | -1.91249 | -1.73251 | -2.26176 | -2.21272 | -1.71405 | -1.21123 |
| JCHAIN | -2.22659 | -2.18212 | -0.34018 | -1.23947 | 6.77419 | 7.075558 | -0.83291 | -0.58804 | 0.306636 | -1.71421 | -2.36679 | -2.31868 | 0.431583 | -0.77897 |
| CA1 | -1.9502 | -2.63252 | -0.22292 | -0.74567 | -2.62911 | -1.82219 | 7.294409 | 5.148496 | -0.01644 | -0.59604 | -2.70168 | -1.71081 | 1.787784 | 0.796893 |
| NAPSB | -1.74774 | -1.70093 | 0.272971 | 0.274594 | 7.222127 | 6.184906 | 0.009258 | -0.7214 | -1.8029 | -1.89079 | -1.25945 | -1.13923 | -1.93515 | -1.76627 |
| AZU1 | -2.64043 | -2.26704 | 2.124785 | 3.668477 | 3.854777 | 4.040325 | 2.956019 | 2.311345 | -2.67401 | -1.17657 | -2.65131 | -2.69941 | -2.73834 | -2.10861 |
| HLA-DRB3 | 3.273109 | -2.79317 | 2.796435 | -2.82473 | 3.655822 | -2.75848 | 1.125459 | -2.81951 | 2.944384 | -2.56811 | 3.086959 | -2.61848 | 2.330259 | -2.82994 |
| LY86 | -0.94144 | -1.5693 | 0.185772 | 1.963541 | 6.716764 | 5.985555 | -1.36727 | -1.474 | -1.66944 | -1.756 | -1.7636 | -1.71205 | -1.79978 | -0.79876 |
| PRG2 | -1.86955 | -1.44908 | -0.86083 | -1.43385 | -1.6585 | -1.07178 | 6.521744 | 6.675125 | -0.98316 | 0.04902 | -1.83645 | -1.67665 | -0.52309 | 0.11706 |
| MPO | -1.9372 | -2.7565 | 2.630516 | 3.295707 | 4.371664 | 3.948335 | 2.746014 | 1.361728 | -2.20362 | -2.20877 | -3.00886 | -3.13459 | -1.53863 | -1.56579 |
| FAM178B | -2.06859 | -2.02469 | -0.44662 | -0.78622 | -1.5915 | -1.93972 | 6.006939 | 5.643762 | -0.68731 | -0.51151 | -2.20746 | -1.70438 | 1.496096 | 0.82119 |
| RNASE2 | -2.15651 | -2.28752 | 2.003406 | 2.921529 | 3.531901 | 2.760084 | 3.439465 | 2.848268 | -2.39957 | -1.44478 | -2.92119 | -2.90209 | -1.71526 | -1.67774 |
| TGM2 | -1.67074 | -1.59512 | -1.25373 | -1.27514 | -1.49964 | 0.87399 | 6.916752 | 4.951895 | -0.97769 | -0.28458 | -1.68233 | -1.76592 | -0.9802 | 0.24245 |
| PKLR | -1.49936 | -1.45524 | -1.4193 | -1.43296 | -1.33309 | -1.37113 | 6.486255 | 5.210931 | -1.55146 | -0.47181 | -1.64202 | -1.59244 | 0.713702 | 1.357929 |
| LOC107984247 | -1.84626 | -1.80307 | -1.76764 | -1.78114 | -1.68176 | 0.883871 | -1.60938 | -1.69622 | -1.89684 | 4.960339 | 3.580762 | 2.206863 | 4.29973 | -1.84926 |
| CNRIP1 | -1.79637 | -1.67078 | -0.0234 | 0.291492 | -2.24769 | -1.74274 | 5.819418 | 5.36122 | 0.006173 | -0.69692 | -2.55594 | -1.92844 | -0.05354 | 1.23751 |
| RNASE3 | -2.14036 | -2.05523 | 1.894237 | 3.105323 | 3.326735 | 2.672203 | 3.580016 | 2.630117 | -2.27742 | -2.34505 | -2.35086 | -2.31104 | -2.37823 | -1.35044 |
| LOC105369205 | -1.41223 | -0.20384 | 0.036007 | -1.34675 | -0.27749 | -1.28588 | -0.87246 | -1.27862 | -1.46369 | 6.223482 | -0.99689 | -1.5042 | 5.812052 | -1.4295 |
| HBB | -1.92592 | -2.88517 | -0.39216 | -0.98568 | -2.45496 | -0.07957 | 5.629944 | 4.562087 | -0.31995 | 0.449487 | -2.7698 | -1.34567 | 1.912603 | 0.604771 |
| FABP4 | -1.13686 | -1.09566 | -1.06233 | -1.07498 | -0.98319 | 8.376089 | -0.91834 | -1.01119 | -1.18587 | 1.065445 | -1.27195 | 0.421552 | -1.30533 | 1.182617 |
| TRIB2 | -2.18972 | -2.14786 | -0.4164 | -0.7101 | -0.16812 | 1.130698 | 5.60594 | 5.24759 | -0.94092 | -0.82141 | -2.32095 | -2.27605 | -0.09843 | 0.105726 |
| ST6GALNAC1 | -1.33143 | -1.2896 | -0.24693 | -1.26851 | -1.1741 | -1.21002 | 6.115516 | 5.253374 | -1.38088 | -1.46 | -1.46695 | -1.41981 | 0.442662 | 0.43668 |
| DNTT | -1.70154 | -2.00077 | 1.137274 | 0.446065 | 5.14996 | 5.044323 | -1.69384 | -0.3979 | 0.075911 | -1.19036 | -3.04091 | -2.63221 | 0.584237 | 0.219765 |
| RNASE6 | -2.29877 | -0.95485 | 0.485569 | 0.593237 | 4.339612 | 3.766452 | 2.758075 | 2.555203 | -1.57822 | -2.4088 | -2.41453 | -2.37523 | -1.1998 | -1.26794 |
| MS4A6A | -0.74296 | -0.81692 | 0.166974 | 1.367909 | 5.603588 | 4.935273 | -0.50282 | -0.46436 | -0.94736 | -1.90434 | -1.91138 | -1.04897 | -1.94477 | -1.78986 |
| RPL36A-HNRNPH2 | -0.22187 | -2.09074 | -2.05866 | -2.0709 | -1.97988 | -0.2703 | -1.91242 | -2.00818 | 3.468482 | 3.305987 | 3.068605 | 1.687049 | 3.227533 | -2.1447 |
| HDC | -1.62984 | -1.58672 | -1.17767 | -1.56488 | -0.51117 | -1.03554 | 5.515338 | 5.116413 | -1.02665 | 0.775582 | -0.93972 | -1.72024 | -1.09895 | 0.884052 |
| MS4A3 | -2.52576 | -2.49436 | 0.804547 | 2.352151 | 2.707238 | 2.682301 | 3.228981 | 3.205011 | -1.4601 | -1.55308 | -2.62049 | -2.58873 | -0.86155 | -0.87616 |
| ECRP | -1.85991 | -1.63109 | 1.752498 | 3.095505 | 3.573991 | 2.830816 | 2.800623 | 1.371144 | -2.0194 | -2.10167 | -2.10783 | -2.06575 | -2.1369 | -1.50203 |

**Supplemental Table 3.** Top 200 differentially expressed genes in ssBM bulk RNAseq samples

| **Donor** | **1** | **2** | **1** | **2** | **1** | **2** | **1** | **2** | **1** | **2** | **1** | **2** | **1** | **2** |
|---|---|---|---|---|---|---|---|---|---|---|---|---|---|---|
| **Population** | ***a*** | ***a*** | ***b*** | ***b*** | ***c*** | ***c*** | ***d*** | ***d*** | ***e*** | ***e*** | ***f*** | ***f*** | ***g*** | ***g*** |
| **Gene** | | | | | | | | | | | | | | |
| CSF1R | -1.24275 | -0.68925 | 0.746281 | 1.275543 | 5.91502 | 4.211567 | -0.04145 | -1.17678 | -1.22124 | -1.01833 | -1.9804 | -1.49369 | -1.76832 | -1.51618 |
| CD180 | -1.28351 | -0.99506 | -0.74756 | 1.467656 | 5.318398 | 5.2096 | -1.06874 | -1.16107 | -0.17033 | -1.04714 | -1.4121 | -1.36745 | -1.4434 | -1.29929 |
| APOC1 | -1.50991 | -1.13514 | -0.43766 | -1.14662 | -1.27628 | -1.5028 | 5.634565 | 5.110984 | -1.38764 | -0.10445 | -1.25843 | -1.19126 | 0.443816 | -0.23916 |
| GSTT1 | -2.17394 | 2.915485 | -2.04794 | 1.45321 | -2.52247 | 0.186979 | -2.21629 | 3.520729 | -2.22705 | 1.889826 | -2.0116 | 2.837584 | -1.80729 | 2.202762 |
| HLA-DRB5 | -2.07833 | 2.982802 | -2.21929 | 2.071962 | -1.80086 | 2.885714 | -2.7199 | 0.882822 | -2.08018 | 1.646059 | -1.97933 | 2.744197 | -2.49837 | 2.162705 |
| EPB42 | -1.15858 | -1.12076 | -1.08998 | -1.10168 | -1.01622 | -1.04875 | 5.755878 | 4.702018 | -1.20323 | -0.36936 | -1.28078 | -1.23833 | -0.87225 | 1.04203 |
| CD2 | -1.19589 | -0.88209 | -1.12625 | -1.13813 | 5.30901 | 5.295354 | -0.66316 | -1.07794 | -0.26415 | -0.79939 | -1.31987 | -1.27682 | 0.350463 | -1.21111 |
| KLF1 | -1.50656 | -1.4458 | 0.047924 | 0.565284 | -1.97425 | -0.85051 | 4.646772 | 4.796152 | -0.53256 | -0.47191 | -2.71319 | -1.97518 | -0.06394 | 1.477762 |
| APOE | -0.40595 | -1.3676 | -0.71869 | -1.49423 | -0.9679 | 1.86375 | 4.566912 | 5.184266 | -1.46093 | -1.27291 | -1.08192 | -0.93789 | -1.87494 | -0.03197 |
| GSTM1 | 2.056226 | -2.11905 | 2.347887 | -2.31631 | 2.746918 | -2.01064 | 2.367497 | -1.79727 | 1.985549 | -2.52601 | 1.684631 | -2.04888 | 1.823798 | -2.19435 |
| GATD3A | -1.60583 | 2.078751 | -2.04339 | 2.276387 | -1.68615 | 2.91054 | -1.95257 | 2.632876 | -2.23845 | 0.976339 | -2.80732 | 1.676096 | -2.26782 | 2.05055 |
| BLNK | -1.05388 | -1.1242 | -0.17572 | -0.40972 | 4.882118 | 5.327226 | -1.24618 | -1.52129 | -0.8962 | -1.19416 | -1.05531 | -0.9164 | 0.075635 | -0.69191 |
| DNASE1L3 | -1.01675 | -0.98108 | -0.95213 | -0.96312 | -0.88303 | 6.98587 | -0.82608 | -0.90753 | -0.80178 | 1.531609 | -1.13264 | -0.43941 | -1.161 | 1.547073 |
| HBG1 | -1.21851 | -1.13378 | 0.210777 | 0.950908 | -1.07655 | -1.10908 | 4.393128 | 5.333707 | -1.35826 | 0.247061 | -1.33953 | -1.29758 | -1.36888 | -1.2334 |
| ALAS2 | -0.93239 | -0.89902 | -0.87189 | -0.88219 | -0.80706 | -0.08812 | 6.679912 | 3.04837 | -0.97185 | -1.0349 | -1.04044 | -1.00289 | -0.5235 | -0.67404 |
| SHD | -1.62896 | -1.07637 | 1.834535 | 1.791603 | 4.587411 | 4.042725 | -0.66499 | -1.50261 | -0.80645 | -1.75248 | -1.75907 | -1.71419 | -1.40164 | 0.05048 |
| KEL | -0.52763 | -0.21413 | -0.58656 | 0.598454 | -1.61679 | -1.65376 | 5.199221 | 4.189107 | -0.56758 | -0.72362 | -1.9097 | -1.57985 | -1.47931 | 0.872143 |
| UBASH3A | -0.09086 | -0.36306 | 0.028689 | -1.24767 | -1.15916 | -1.19291 | 5.606089 | 4.092902 | 0.271535 | -1.42527 | -1.43167 | -1.48949 | -0.27739 | -1.32175 |
| LOC112268313 | 3.048334 | -1.99913 | -1.96705 | 3.793539 | 2.376577 | 0.40817 | -1.82123 | 2.807109 | -0.05411 | -0.54222 | -1.52219 | -2.11778 | -0.35686 | -2.05316 |
| LOC102723630 | 1.501971 | -2.02716 | 2.366524 | -2.00853 | 2.410091 | -1.95583 | 2.480331 | -1.94944 | 2.039054 | -2.17052 | 1.688556 | -2.13785 | 1.840537 | -2.07774 |
| EPX | -0.81352 | -0.76174 | -0.99024 | -0.17519 | -0.20838 | -0.26551 | 5.752911 | 4.049782 | -0.87792 | -1.15942 | -1.16513 | -1.12636 | -1.19226 | -1.06704 |
| IFIT1B | -0.75494 | -0.96421 | -0.66619 | -0.10565 | -0.87109 | -0.20159 | 5.812178 | 3.995441 | -1.03744 | -1.1005 | -1.10602 | -0.85659 | -1.13225 | -1.01113 |
| MNDA | -0.24075 | -1.76604 | 0.359159 | 2.089789 | 3.734322 | 3.331525 | 2.014536 | 1.385211 | -1.85158 | -1.92381 | -1.93008 | -1.51505 | -1.86617 | -1.82106 |
| AHSP | -1.60421 | -1.32082 | -0.74874 | -1.60217 | -1.94772 | -0.93878 | 4.74597 | 3.832763 | -0.28311 | 0.262829 | -0.87761 | -2.00814 | 1.983804 | 0.505938 |
| TNNI2 | -0.30809 | -1.4117 | 0.43469 | -0.26996 | 4.298947 | 5.184565 | -1.23391 | -0.64995 | -1.36663 | -1.21712 | -1.5801 | -1.53571 | 0.103101 | -0.44812 |
| HBD | -1.76191 | -1.7021 | 0.644439 | 1.086924 | -2.46268 | -1.77698 | 3.687742 | 3.999695 | -0.46305 | 0.988444 | -2.52587 | -1.41494 | 0.368617 | 1.331676 |
| GATA1 | -1.29347 | -1.2859 | 0.234775 | 0.640511 | -2.15341 | -1.40843 | 4.045612 | 3.910321 | -0.30831 | -0.12474 | -2.8484 | -1.62144 | 0.150517 | 2.062366 |
| UGT3A2 | -1.99704 | -1.42284 | 1.704815 | 1.843075 | 3.671225 | 3.999622 | 0.964006 | 0.083896 | -0.55761 | -1.89128 | -1.51113 | -2.07851 | -1.45221 | -1.35601 |
| LOC107986126 | -1.44076 | -0.99382 | -1.36887 | -1.38117 | -0.52664 | -1.32533 | -0.40952 | 0.905419 | -1.48723 | 5.301932 | 0.581697 | -0.90765 | 3.832442 | -0.7805 |
| SULT1A3 | -1.68519 | -1.62887 | -1.59748 | -1.61913 | -1.52336 | -1.57063 | 4.163194 | 1.103282 | -1.75698 | 2.320675 | 2.17145 | -1.78374 | 2.026072 | 1.38071 |
| ELK2AP | 5.391483 | -0.34959 | -0.85508 | -0.68674 | -0.77638 | 0.100026 | 4.187628 | -0.79544 | -0.7663 | -1.31112 | -1.31714 | -1.11191 | -0.74786 | -0.96157 |
| KIF17 | -1.32404 | -1.42804 | 0.707096 | 1.123819 | 3.75954 | 4.976934 | -1.43988 | -0.74593 | -1.54563 | 0.041667 | -1.79272 | -1.0876 | -0.04348 | -1.20174 |
| C4A | -0.7928 | -1.0623 | -1.80357 | -1.01357 | -1.0843 | 0.055216 | 3.601973 | 5.634299 | -0.29775 | -0.16265 | -1.17807 | -0.95029 | -0.77073 | -0.17545 |

**Supplemental Table 3.** Top 200 differentially expressed genes in ssBM bulk RNAseq samples

| Donor | 1 | 2 | 1 | 2 | 1 | 2 | 1 | 2 | 1 | 2 | 1 | 2 | 1 | 2 |
|---|---|---|---|---|---|---|---|---|---|---|---|---|---|---|
| Population | *a* | *a* | *b* | *b* | *c* | *c* | *d* | *d* | *e* | *e* | *f* | *f* | *g* | *g* |
| Gene | | | | | | | | | | | | | | |
| LOC105379461 | -1.72882 | -1.47055 | 0.522528 | 0.015722 | 0.112419 | -0.31117 | 5.124692 | 3.696335 | -1.38432 | -0.60763 | -1.85928 | -1.68011 | -0.84302 | 0.413207 |
| LYPD6B | 2.781282 | -1.46225 | 1.231942 | -1.44284 | -1.35501 | -1.38862 | -1.2912 | -1.38211 | 3.274964 | -1.29801 | 3.365405 | -1.58028 | 2.062311 | -1.51559 |
| TPSAB1 | -2.19547 | -1.91022 | 0.952105 | 0.650973 | 0.889667 | 0.413352 | 3.673273 | 3.217556 | -0.47069 | -0.73192 | -2.83882 | -2.83972 | -0.02764 | 1.217559 |
| CLEC12A | -2.19664 | -1.91626 | 1.873207 | 2.547796 | 3.171936 | 2.944118 | 1.174419 | 1.017384 | -1.41537 | -1.43494 | -1.81727 | -1.28079 | -1.21646 | -1.45112 |
| RHAG | -1.91184 | -1.87092 | -0.8227 | -0.81217 | -1.27124 | -0.59726 | 4.788141 | 4.017126 | -1.02553 | -0.26037 | -0.34684 | -0.71243 | 0.719813 | 0.106214 |
| UCA1 | -1.43638 | -1.54585 | -0.76794 | -0.2946 | -1.19008 | -1.09041 | 4.356113 | 4.59462 | -0.69035 | -0.58142 | -0.83492 | -0.82784 | 0.212501 | 0.09655 |
| NID1 | -0.92279 | -0.06311 | 0.87735 | -0.87635 | -0.28234 | -0.55716 | 6.311666 | 1.24619 | -0.72658 | -1.01677 | -1.02181 | -0.98755 | -1.04573 | -0.935 |
| UBXN10 | 0.025738 | -1.18804 | -1.15865 | -1.16983 | -1.08781 | -0.07562 | 5.540968 | 2.841179 | 0.515682 | -1.33327 | -0.60077 | -0.41208 | -1.36691 | -0.53059 |
| ANK1 | -1.61179 | -1.39893 | 0.519901 | 0.75251 | -1.35783 | -1.08962 | 4.143859 | 3.995808 | -0.23432 | -0.28199 | -2.10084 | -1.50579 | 0.050096 | 0.11893 |
| SUCNR1 | -0.65843 | -1.56671 | 1.422601 | 1.929299 | 3.533134 | 2.98922 | 0.934799 | 0.866387 | -1.43029 | -1.99611 | -2.10934 | -2.0694 | -1.24594 | -0.59922 |
| PLPPR3 | -1.57694 | -1.54012 | 1.373482 | 2.636596 | 2.735853 | 2.826907 | 1.972828 | 0.745479 | -1.61989 | -1.62137 | -1.69312 | -1.65325 | -1.72077 | -0.86571 |
| SIGLEC12 | -1.12708 | -1.09379 | 0.787186 | 1.854965 | 2.637806 | 4.956475 | 0.23162 | -1.0242 | -1.16614 | -1.2279 | -1.23328 | -1.19664 | -1.25881 | -1.14022 |
| C4B | -0.61623 | -0.58601 | 0.986555 | -0.57104 | -0.50631 | -0.5305 | 6.491553 | -0.52576 | -0.65291 | -0.71372 | -0.71919 | -0.6825 | -0.74553 | -0.62842 |
| CD40LG | -1.73787 | -0.59433 | 0.484476 | 0.39281 | -1.39868 | -0.16361 | 4.28024 | 3.694921 | -0.92798 | -0.32615 | -1.91643 | -1.50851 | -1.2729 | 0.994011 |
| MPEG1 | -1.02236 | -1.14537 | -0.37758 | 0.317272 | 4.761104 | 3.982669 | -0.9391 | -1.05052 | -0.59225 | -0.93321 | -0.26467 | -0.99263 | -0.98969 | -0.75366 |
| CTSG | -1.31256 | -1.01164 | 1.330881 | 2.100297 | 2.745337 | 2.797835 | 2.151034 | 0.993091 | -1.51604 | -2.00199 | -1.35062 | -1.68058 | -1.90122 | -1.34382 |
| LEF1 | -1.65579 | -1.61673 | 0.276228 | -0.29246 | 0.191404 | 1.205906 | 4.122685 | 3.469565 | 0.006285 | 0.02371 | -1.77949 | -1.73695 | -0.54321 | -1.67117 |
| IGFBP2 | -0.78632 | -1.31234 | 2.031223 | 1.737892 | 3.269367 | 2.422221 | 1.792641 | 0.350996 | -1.37862 | -2.0633 | -1.90178 | -1.83827 | -0.96307 | -1.36062 |
| CEBPD | -1.1254 | -1.75259 | 1.163815 | 2.15814 | 3.469438 | 3.351046 | 0.489869 | 0.212244 | -1.18103 | -1.97487 | -1.07545 | -1.38943 | -0.86787 | -1.47791 |
| NMU | -0.47933 | -0.35383 | -1.57276 | -1.25178 | -1.49411 | -0.49733 | 5.229214 | 2.332361 | -0.87858 | -0.9205 | -1.1445 | -0.58413 | 1.358507 | 0.256762 |
| CSTA | -0.60501 | -1.56174 | 1.2135 | 2.041193 | 3.34897 | 2.831722 | 1.090969 | 0.963049 | -1.64195 | -1.7097 | -1.71557 | -1.67549 | -0.96663 | -1.61333 |
| HBG2 | -0.60333 | -1.07895 | -0.14778 | 1.012893 | -0.78842 | -1.43682 | 2.819929 | 4.992931 | -1.52056 | 0.233277 | -1.67304 | -0.76229 | -0.46417 | -0.58367 |
| SLC10A4 | 0.101272 | -1.04787 | -1.02099 | -0.0012 | 0.272593 | 0.404175 | -0.90199 | 5.932305 | -1.11934 | 0.438536 | -1.18577 | -1.14951 | -1.21105 | 0.488835 |
| AVP | 1.880565 | 1.661576 | 0.307844 | -0.15287 | -2.37833 | -2.4902 | -2.8811 | -2.83687 | 0.95442 | 1.195278 | 1.749845 | 1.675445 | 0.922001 | 0.392396 |
| RNASE1 | -1.61932 | -1.34257 | 0.215454 | 0.098584 | 0.135253 | 0.182282 | 4.019178 | 3.812153 | -1.28053 | -0.30345 | -1.74347 | -1.70074 | -0.75289 | 0.280069 |
| IGSF6 | -1.16989 | -1.33902 | 0.045229 | 0.506965 | 5.104348 | 2.72724 | -0.87603 | -0.64395 | -0.28671 | -0.54029 | -0.22827 | -1.45274 | -0.85299 | -0.99389 |
| ALOX5AP | -0.82491 | -1.97889 | -0.43807 | -0.21003 | 2.498463 | 2.136214 | 2.842431 | 2.95589 | -1.2526 | -0.64469 | -2.0745 | -1.32343 | -1.24276 | -0.44311 |
| LOC101928134 | -1.298 | -0.46027 | -1.23298 | -0.47156 | -1.16217 | -0.89256 | 3.353974 | 4.593999 | -1.22821 | -0.96232 | -0.86715 | 0.197605 | -0.36681 | 0.796456 |
| CR2 | -0.93973 | -0.91085 | -0.88717 | -0.89618 | -0.82986 | -0.85522 | 5.117915 | 2.543687 | -0.5784 | -1.02676 | 0.98905 | 0.277927 | -1.05328 | -0.95112 |
| TPSAB1 | -1.9321 | -0.17406 | -1.42811 | 1.478068 | -1.78772 | 1.769007 | 0.10865 | 3.781145 | -1.00186 | 1.094009 | -2.04897 | -1.21842 | -0.80304 | 2.163397 |
| CSF1 | -2.01819 | -1.31099 | 0.140837 | -0.10835 | -2.32242 | -1.47752 | 3.472911 | 3.075689 | 0.225765 | 0.449851 | -1.63829 | -0.78321 | 0.977342 | 1.316572 |
| CCL14 | -0.85454 | -0.82767 | -0.80562 | -0.81402 | -0.75226 | 5.708589 | -0.7075 | -0.7713 | -0.88596 | 1.044785 | -0.79923 | -0.67928 | 0.302553 | 0.841443 |
| AMHR2 | -1.76037 | -0.75786 | -1.08158 | 0.551038 | -1.61558 | -1.64933 | 3.210422 | 3.427141 | -0.05059 | 0.61353 | -1.87889 | -0.98527 | 0.175897 | 1.801443 |

**Supplemental Table 3.** Top 200 differentially expressed genes in ssBM bulk RNAseq samples

| **Donor** | **1** | **2** | **1** | **2** | **1** | **2** | **1** | **2** | **1** | **2** | **1** | **2** | **1** | **2** |
|---|---|---|---|---|---|---|---|---|---|---|---|---|---|---|
| **Population** | ***a*** | ***a*** | ***b*** | ***b*** | ***c*** | ***c*** | ***d*** | ***d*** | ***e*** | ***e*** | ***f*** | ***f*** | ***g*** | ***g*** |
| **Gene** | | | | | | | | | | | | | | |
| LINC01670 | -1.56919 | 1.314864 | 2.231245 | -0.3597 | 2.716332 | -1.46494 | 2.735774 | 1.400783 | -1.61043 | -1.67491 | -1.68049 | -0.51402 | -1.70686 | 0.181531 |
| ITGB7 | -0.5811 | -1.83582 | -0.12905 | -1.88706 | 2.305 | 2.167323 | 3.093081 | 2.085348 | 0.592143 | -1.8832 | -0.7537 | -1.63321 | -0.6884 | -0.85137 |
| CD36 | -0.44851 | -1.9822 | 0.019552 | -0.30997 | 0.304139 | 0.830698 | 3.974776 | 3.379566 | -1.43991 | -1.11537 | -1.34364 | -1.19678 | -0.53918 | -0.13316 |
| TPSD1 | -1.32426 | -0.1817 | -1.26171 | 1.381608 | -1.19323 | 1.328477 | -0.17315 | 4.223137 | -1.36439 | 0.320721 | -1.433 | -1.11591 | -1.45895 | 2.252347 |
| HBA1 | -1.51703 | -2.12768 | -0.11305 | -1.40673 | -2.02985 | 1.543498 | 2.290468 | 2.763454 | 0.109612 | 0.751624 | -2.26457 | -0.7617 | 1.86801 | 0.893949 |
| CPA3 | -1.96003 | -2.19937 | 1.615428 | 1.88583 | 2.015085 | 1.587659 | 1.441735 | 1.378345 | -0.73134 | -0.55435 | -2.37046 | -2.43972 | -0.41155 | 0.742739 |
| ACSM1 | -0.65697 | -1.4886 | 0.281161 | 1.120108 | -0.0236 | -0.6478 | 3.751498 | 3.32575 | -1.32321 | -0.51802 | -1.77533 | -1.88028 | -0.35576 | 0.191044 |
| MEG3 | 1.551591 | 0.780789 | -0.3824 | -1.27158 | -2.29439 | -2.17602 | -2.38515 | -2.16549 | 1.430081 | 1.197607 | 2.364312 | 1.346779 | 1.418857 | 0.585006 |
| LGALS1 | -1.24245 | -1.29928 | 0.649826 | 1.209479 | 3.597146 | 3.599959 | -0.3156 | -0.26145 | -1.54911 | -0.6877 | -1.34802 | -1.14124 | -0.41511 | -0.79645 |
| IGLL1 | -0.96515 | -1.06108 | 1.620705 | 1.927606 | 2.663134 | 3.118652 | 0.549091 | 0.040937 | -1.29084 | -1.40075 | -2.06773 | -1.71061 | -0.89726 | -0.5267 |
| LINC01115 | -1.16876 | 2.988704 | -1.10939 | 0.548556 | -1.04472 | -1.07334 | -0.99046 | -1.06779 | -1.207 | 1.52208 | -1.27264 | 2.991065 | -1.29755 | 2.181242 |
| CYP2E1 | -0.12204 | -1.27138 | -0.33168 | 0.907314 | 1.152218 | 0.639901 | 4.551117 | 1.612126 | -1.03573 | -1.40703 | -1.41243 | -0.5258 | -1.43799 | -1.3186 |
| CTSL | -0.87126 | -0.23345 | -1.10764 | -1.25096 | -1.70284 | 1.024073 | 2.850572 | 4.12012 | -0.51862 | 0.599973 | -0.56306 | -0.15839 | -1.62465 | -0.56387 |
| DLK1 | 1.570115 | 1.628969 | -0.25624 | -0.47785 | -2.41596 | -2.54332 | -2.07724 | -2.36048 | 1.383945 | 1.040836 | 1.592084 | 1.564048 | 0.762434 | 0.588657 |
| FUT7 | -0.43857 | -0.49538 | 1.63465 | 1.530964 | 3.210577 | 3.125555 | -0.43256 | -0.66407 | -0.71188 | -1.95423 | -1.28932 | -1.40567 | -1.14764 | -0.96242 |
| GSTM5 | 1.629127 | -0.44637 | 1.89152 | -2.3849 | 2.095149 | -2.53773 | 0.72568 | -2.08002 | 1.511056 | -0.99732 | 1.42568 | -0.58163 | 1.067407 | -1.31764 |
| LINC02132 | -0.70763 | -0.92355 | 0.428673 | 0.974006 | 3.412052 | 3.754342 | -1.38376 | -0.80184 | -0.35805 | -1.24412 | -0.54174 | -0.87233 | -1.51473 | -0.22132 |
| EPAS1 | 1.509349 | 1.569251 | -0.03878 | -0.51564 | -1.73636 | -2.41925 | -2.57756 | -2.49515 | 1.413553 | 1.449822 | 1.48527 | 1.439637 | 0.586146 | 0.329714 |
| SULT1A4 | -0.40992 | 0.443274 | 0.083662 | 0.364616 | -0.49012 | 0.055954 | 2.911169 | 2.966846 | -0.09475 | -1.06664 | -2.80531 | 0.154421 | -2.82248 | 0.709278 |
| RNA28SN2 | -1.65603 | -1.91257 | -0.64753 | -0.11565 | -0.47476 | -0.20585 | -1.85471 | -1.30918 | 1.055629 | 2.527953 | 2.299629 | 0.452147 | 2.981626 | -1.1407 |
| DNAAF4-CCPG1 | -1.79152 | -1.75439 | -1.7335 | 1.480386 | 1.425823 | 1.939576 | -1.60597 | -1.68962 | 0.471463 | 1.799585 | 1.345211 | -0.08274 | 2.003613 | -1.80791 |
| HES6 | -1.02589 | -1.05514 | 0.576099 | 0.384989 | 0.968432 | 1.447253 | 3.161478 | 2.808138 | -0.75707 | -1.68675 | -1.35536 | -2.18906 | -1.23079 | -0.04633 |
| ST14 | -1.04795 | 0.735249 | 0.589234 | 0.668631 | 3.472488 | 2.903192 | -0.6032 | -1.48636 | 0.799421 | -0.35488 | -1.58552 | -1.17176 | -1.74191 | -1.17663 |
| VPREB1 | -0.88408 | -1.04282 | 1.15407 | 0.843799 | 2.741605 | 3.772359 | -1.1626 | -0.52538 | -0.62918 | -0.46371 | -2.07501 | -1.50138 | -0.11096 | -0.11672 |
| TCEAL2 | 1.392014 | 1.610667 | -0.61204 | -0.41827 | -2.22568 | -2.45163 | -1.89206 | -2.32995 | 1.324091 | 0.772868 | 1.706523 | 1.746139 | 0.619466 | 0.757868 |
| CCDC169-SOHLH2 | -1.9446 | 1.771587 | -1.88387 | 1.303034 | -1.81562 | 0.430598 | -0.28147 | 2.202934 | -1.77027 | 1.400784 | -2.04646 | 1.338691 | -0.21217 | 1.50682 |
| FBLN2 | -1.72374 | -1.51865 | -0.32128 | 1.321953 | 1.22665 | 0.378871 | 2.426321 | 3.093813 | -0.61029 | -0.55647 | -1.83392 | -1.53466 | -1.43753 | 1.088937 |
| CLEC3B | 1.281491 | 1.935856 | -1.22543 | -0.38168 | -1.92914 | -1.7285 | -2.6412 | -2.04594 | 0.931634 | 1.082296 | 1.376401 | 2.009575 | 0.478051 | 0.856594 |
| HLA-DPB2 | -1.68318 | 2.098989 | -1.45315 | 1.559165 | -1.1605 | 1.859098 | -1.23274 | 0.299568 | -1.33873 | 1.259828 | -1.49548 | 1.915912 | -2.05451 | 1.425742 |
| LTBP1 | -1.00899 | -0.40735 | 0.482983 | 1.273475 | -1.83856 | -1.87389 | 1.474626 | 1.527678 | 0.128042 | 1.759154 | -2.60132 | -2.03672 | 1.138498 | 1.982373 |
| SNHG18 | -1.42934 | 1.99619 | -1.37069 | 1.753836 | -1.30562 | 1.882953 | -1.25012 | -0.48326 | -1.46656 | 1.537238 | -1.52935 | 2.384328 | -1.55281 | 0.833211 |
| VPREB3 | -1.34207 | -1.61193 | -1.2089 | -1.25199 | 1.943435 | 3.124128 | 0.336239 | 1.447809 | 0.304523 | -0.79114 | -1.75579 | -1.71858 | 1.784182 | 0.740078 |
| ITGA2B | -0.46342 | -0.31291 | 0.190296 | 0.915814 | -2.57764 | -1.51752 | 2.687571 | 2.497699 | -0.36663 | 0.798154 | -1.81642 | -1.56505 | -0.07089 | 1.600946 |

**Supplemental Table 3.** Top 200 differentially expressed genes in ssBM bulk RNAseq samples

| **Donor** | **1** | **2** | **1** | **2** | **1** | **2** | **1** | **2** | **1** | **2** | **1** | **2** | **1** | **2** |
|---|---|---|---|---|---|---|---|---|---|---|---|---|---|---|
| **Population** | ***a*** | ***a*** | ***b*** | ***b*** | ***c*** | ***c*** | ***d*** | ***d*** | ***e*** | ***e*** | ***f*** | ***f*** | ***g*** | ***g*** |
| **Gene** | | | | | | | | | | | | | | |
| XK | -1.03786 | -0.55401 | 0.823282 | 0.25433 | -1.27999 | -1.84879 | 3.177961 | 2.80249 | -1.36546 | 0.747212 | -0.66672 | -2.02727 | 0.349162 | 0.625663 |
| LOC101927879 | 0.417238 | 1.240484 | -0.73383 | -0.26287 | -2.65844 | -2.48761 | -1.75042 | -1.63893 | 0.896862 | 1.987633 | 1.129426 | 1.613167 | 1.344523 | 0.902771 |
| ACY3 | -0.31097 | -0.04648 | 0.90988 | -0.36233 | 3.611798 | 2.758742 | -1.37578 | -2.27157 | 0.593139 | -1.32169 | -0.32729 | -0.71271 | -0.17673 | -0.96801 |
| FAM83D | -0.81864 | -0.3675 | 0.418196 | 0.235778 | 0.414296 | 0.756684 | 2.98036 | 3.1209 | -0.61117 | -1.99011 | -1.30529 | -1.75845 | -1.59454 | 0.51948 |
| PMP22 | -0.69617 | -0.47331 | -0.62774 | -0.48914 | 0.707479 | -0.97573 | 3.799962 | 3.190017 | -0.56799 | -1.02724 | -0.41106 | -1.64901 | -0.26689 | -0.51318 |
| DYNLL1 | -1.18656 | -0.87576 | -0.96291 | 0.871406 | -0.45382 | -0.77551 | 3.224568 | 3.10457 | -0.3147 | 0.183799 | -1.91012 | -1.58802 | -0.26463 | 0.947676 |
| CSF2RB | -1.34092 | -0.57155 | -0.45768 | -0.60122 | -0.67539 | -1.95794 | 3.398241 | 3.415681 | -0.46123 | 0.136329 | -0.68355 | -0.90126 | 0.260472 | 0.440016 |
| BORCS7-ASMT | 1.067155 | -1.91562 | 0.061468 | -1.91012 | -1.82317 | 1.343516 | -1.77508 | -1.85711 | 1.074313 | 1.506462 | 1.677816 | 1.636959 | 1.560651 | -0.64725 |
| HCK | -0.94266 | -0.88534 | 2.165997 | 1.742523 | 3.087414 | 2.093045 | -1.11005 | -1.18873 | -1.01402 | -1.3884 | -1.39359 | -0.80593 | 0.241171 | -0.60142 |
| SPON1 | 0.287406 | -1.31271 | 1.064666 | -0.34744 | 3.345571 | 2.418285 | -1.26076 | -1.54428 | 1.119821 | -0.23245 | -1.76598 | -1.72804 | -0.27601 | 0.231923 |
| DTX4 | -1.51365 | -0.08675 | 0.479062 | 0.105679 | 3.538584 | 3.223289 | -0.6337 | -0.0655 | -1.37381 | -1.23953 | -0.17183 | -1.26826 | -0.79835 | -0.19525 |
| TFR2 | -0.78555 | -0.37113 | 0.163436 | -0.10479 | -1.12726 | -0.79764 | 3.480635 | 2.956839 | -0.61221 | 0.006549 | -2.1849 | -1.30184 | -0.30274 | 0.980592 |
| SEC14L2 | -0.33617 | -0.84987 | -1.29261 | -0.84782 | -1.64219 | -1.64049 | 3.539206 | 3.06216 | 0.320945 | 0.337331 | -0.44603 | 0.194294 | -0.59014 | 0.1914 |
| UMODL1 | -0.61951 | -0.20382 | 0.26681 | 0.305438 | 3.217274 | 3.513916 | -1.09426 | -1.32552 | -0.27773 | -0.38012 | -0.24746 | -0.6145 | -1.88983 | -0.65068 |
| P2RX5 | -0.42133 | -1.20998 | 0.534633 | 0.329947 | 1.107411 | -0.72691 | 3.835468 | 2.085558 | -0.17634 | -0.8237 | -0.95748 | -2.28629 | -0.43221 | -0.85877 |
| CCR7 | -1.38063 | -1.3457 | -1.31702 | 0.040391 | 3.298554 | 3.232084 | -0.57775 | -0.23963 | 0.854864 | -0.42569 | -1.49121 | -0.13111 | -0.11596 | -0.4012 |
| DHRS9 | -1.66022 | -1.62616 | -1.03262 | -0.94817 | -0.68749 | -0.6683 | 1.419649 | 2.588358 | 0.295464 | 1.963223 | -1.76626 | -0.57381 | 0.271199 | 2.425149 |
| DHRS3 | -1.6908 | -1.20916 | -0.05214 | 0.735409 | 0.921459 | 1.397178 | 2.398402 | 2.551754 | -1.44662 | -0.33007 | -2.15808 | -1.62048 | -0.33867 | 0.841807 |
| TST | -0.3974 | -0.79968 | 0.402592 | 0.532361 | 1.159727 | 1.263813 | 2.718987 | 2.781786 | -1.46862 | -0.98661 | -1.62832 | -1.91766 | -0.9437 | -0.71727 |
| LOC112268350 | -1.21306 | 0.921962 | -1.16066 | -1.16385 | -1.09028 | -1.12236 | -1.04642 | 2.03571 | -1.25879 | 2.948805 | -1.32593 | 1.1439 | 0.345812 | 1.985164 |
| RASSF6 | 1.131337 | 1.856241 | -0.71964 | -0.55757 | -2.21208 | -2.00268 | -1.55216 | -2.23756 | 0.988114 | 0.845165 | 1.548291 | 1.73796 | 0.15888 | 1.01571 |
| SLC1A6 | 1.529849 | 1.473842 | 0.143225 | -0.31268 | -2.69501 | -2.58083 | -1.15987 | -1.77121 | 1.681294 | -0.00756 | 1.237176 | 1.021766 | 0.493495 | 0.946524 |
| AKR1C1 | -0.6395 | -0.18894 | -0.56617 | 0.306483 | -1.50687 | -0.66727 | 2.452249 | 3.687614 | -0.81292 | -0.54388 | -1.30207 | 0.890287 | -1.54388 | 0.434859 |
| ADAMTS14 | -0.72343 | -0.73913 | 0.624967 | -0.54935 | 0.707957 | 0.702674 | 3.310926 | 2.497262 | -0.60849 | -0.74 | -1.88329 | -1.8455 | -1.10714 | 0.352553 |
| RBPMS2 | -0.11903 | -0.43166 | 0.309131 | 1.035446 | -1.17205 | -0.50041 | 2.962272 | 2.795053 | -1.39637 | 0.128011 | -2.23474 | -0.80964 | -1.14773 | 0.581713 |
| F13A1 | -0.28685 | -0.01653 | 1.197774 | 1.100117 | 2.726624 | 2.183889 | 0.800898 | 0.03185 | -0.90205 | -1.35152 | -2.34986 | -1.66588 | -1.35606 | -0.11241 |
| LINC01835 | -1.2531 | -0.80343 | -0.05556 | 1.610381 | -0.38609 | -0.78275 | 2.557134 | 3.008351 | -1.01202 | 0.472017 | -1.35308 | -1.3188 | -1.37682 | 0.693753 |
| C7orf55-LUC7L2 | -1.86788 | 0.077926 | -0.23084 | -0.4488 | -1.74045 | 2.332342 | 0.201822 | 1.133827 | 0.308915 | 2.194723 | -1.98141 | 1.346022 | -1.90367 | 0.577469 |
| FCER1A | -2.29435 | 0.898157 | -1.2801 | 0.178753 | -1.32712 | -1.44205 | 1.112313 | 1.737999 | -0.1812 | 1.191604 | -2.26507 | 0.904607 | 0.676792 | 2.089662 |
| ZAR1 | -0.03728 | -0.0991 | -0.40606 | 1.096197 | 0.598471 | -1.69129 | -1.53113 | 0.588186 | -1.88541 | 2.457958 | 1.837552 | -1.32544 | 1.925661 | -1.52831 |
| KCNG2 | -1.51618 | 0.097832 | 1.402605 | 1.79902 | 2.274127 | 2.573382 | -0.50997 | -0.19997 | -1.2474 | -0.93554 | -1.978 | -0.45673 | -1.36359 | 0.060401 |
| CLEC4G | 0.105901 | -1.15593 | 0.621843 | 0.594078 | 3.107889 | 2.987531 | 0.391697 | -1.08891 | -0.76011 | -0.3276 | -1.28894 | -0.67396 | -1.31307 | -1.20041 |
| PPP1R14A | -1.03095 | -0.12016 | -0.10179 | 0.551331 | -1.05077 | -0.76993 | 3.144893 | 3.19079 | -0.8867 | -0.41551 | -1.36369 | -0.96396 | -0.77847 | 0.594905 |

**Supplemental Table 3.** Top 200 differentially expressed genes in ssBM bulk RNAseq samples

| Donor | 1 | 2 | 1 | 2 | 1 | 2 | 1 | 2 | 1 | 2 | 1 | 2 | 1 | 2 |
|---|---|---|---|---|---|---|---|---|---|---|---|---|---|---|
| Population | *a* | *a* | *b* | *b* | *c* | *c* | *d* | *d* | *e* | *e* | *f* | *f* | *g* | *g* |
| Gene | | | | | | | | | | | | | | |
| MS4A1 | -0.73162 | -0.84784 | -1.14585 | -0.55117 | -1.49656 | -0.38805 | 2.906746 | 3.662374 | -0.27433 | 0.227996 | -0.39555 | -0.55743 | -0.36196 | -0.04676 |
| BLOC1S5-TXNDC5 | -1.39195 | -0.15256 | -1.33662 | 0.326149 | 0.870059 | -1.30234 | -0.13865 | 0.875241 | -1.42682 | 2.167173 | 1.815793 | -1.45357 | 2.55185 | -1.40376 |
| PDZRN4 | 0.973247 | 1.77383 | -0.04052 | 0.178537 | -2.28225 | -2.30895 | -1.46993 | -2.03744 | 0.515297 | 1.284494 | 0.520463 | 1.72142 | -0.07857 | 1.250376 |
| TARP | -1.18781 | -1.36433 | 1.317314 | 1.673211 | 2.290586 | 1.985161 | 1.277324 | 0.720642 | -1.42455 | -1.06378 | -1.51988 | -1.30282 | -0.71338 | -0.68769 |
| CDH7 | 0.747192 | 1.567049 | -1.38701 | 0.006664 | -2.1255 | -1.69847 | -1.8945 | -1.6411 | 0.858081 | 1.564195 | 1.08772 | 1.838223 | 0.240611 | 0.836858 |
| LIMCH1 | 1.162694 | 1.430235 | 0.034456 | -0.20539 | -2.06131 | -2.11301 | -2.00135 | -2.19201 | 1.047688 | 1.034116 | 1.14495 | 1.261511 | 0.757291 | 0.700121 |
| GPR75-ASB3 | 0.803165 | 0.743059 | 0.546741 | 0.542273 | 1.466486 | 0.901274 | 1.284792 | 0.399919 | 0.523864 | -2.56295 | -2.58304 | 0.096562 | -2.58366 | 0.42152 |
| MYL4 | -1.15891 | -1.34286 | -0.02148 | 0.071998 | -1.81736 | -0.88231 | 2.545413 | 2.33155 | -0.04052 | 0.854745 | -1.95289 | -0.62303 | 0.429972 | 1.605686 |
| CRHBP | 1.218091 | 1.727928 | 0.215652 | -0.0584 | -2.29288 | -2.36555 | -1.70432 | -1.82204 | 1.080396 | 0.494241 | 0.889812 | 1.476843 | 0.285955 | 0.854269 |
| CST7 | -1.30145 | -0.64295 | 0.931779 | 1.762921 | 2.041448 | 1.870174 | 1.493543 | 1.027604 | -1.3641 | -1.44357 | -1.6237 | -1.28663 | -1.171 | -0.29406 |
| MINOS1-NBL1 | 0.726774 | -0.25506 | -1.88889 | -0.2263 | -1.81489 | -0.85889 | -1.89699 | -0.58117 | 1.424167 | 1.629565 | 1.934975 | 0.842682 | 2.031346 | -1.06732 |
| HES1 | 1.264481 | 0.835009 | -0.6733 | -1.17217 | -2.0067 | -1.32138 | -1.9188 | -1.71648 | 1.336889 | 1.285901 | 1.861849 | 1.253126 | 1.195733 | -0.22415 |
| SELENOM | 1.162237 | 1.616763 | -0.66273 | -0.91811 | -2.11777 | -1.32259 | -2.18918 | -1.66284 | 1.007306 | 0.918561 | 1.526855 | 1.645259 | 0.570333 | 0.4259 |
| CHST2 | -0.93321 | -0.25188 | -0.00109 | 0.034108 | 1.289442 | 0.369859 | 3.06227 | 2.635501 | -0.8357 | -1.67546 | -1.16442 | -1.09593 | -1.06303 | -0.37045 |
| LOC403323 | 1.304367 | -1.71199 | 1.494877 | -1.3962 | 1.585364 | -1.12196 | 1.352641 | -1.30806 | 1.131069 | -1.35136 | 1.084101 | -0.78391 | 1.476687 | -1.75562 |
| SERPINB8 | -0.06757 | -0.43842 | 0.965604 | 1.106067 | 2.929301 | 2.515531 | 0.194696 | -1.37571 | -0.47945 | -1.952 | -0.78769 | -0.92057 | -0.66477 | -1.02502 |
| SELENBP1 | -0.25028 | 0.093714 | -0.86561 | -1.42906 | -1.47221 | -0.36026 | 3.358588 | 2.904074 | -0.1397 | -0.44329 | -0.19445 | -0.18813 | -0.13984 | -0.87356 |
| LOC107986939 | 1.338138 | 1.419951 | -0.01149 | -0.55277 | -1.35711 | -1.96152 | -2.25451 | -2.25102 | 1.023602 | 0.945596 | 1.224882 | 1.326521 | 0.702912 | 0.406828 |
| GGTA1P | 0.10117 | 1.977263 | -1.47945 | 0.893928 | -2.67023 | -0.41167 | -2.11263 | -0.19656 | -0.10437 | 1.162142 | 0.012525 | 1.782785 | -0.47787 | 1.522975 |
| NTRK1 | -0.88326 | -1.02498 | -0.86051 | 0.300981 | -1.46894 | -1.49936 | 2.804251 | 2.630479 | 0.010045 | 0.279215 | -0.97039 | -0.79778 | -0.01016 | 1.490424 |
| LOC112268246 | 0.992364 | -1.70248 | 0.855069 | -1.54712 | 0.677481 | -1.23015 | -0.06784 | -1.43728 | 1.016392 | 1.973133 | 1.303842 | -1.10746 | 1.975296 | -1.70125 |
| PLK1 | -0.53957 | -0.02103 | 0.680287 | 0.703332 | 1.988174 | 2.107834 | 1.357051 | 1.295619 | -1.8115 | -1.52533 | -1.61002 | -0.88679 | -1.8724 | 0.13435 |
| ABI3BP | 1.090228 | 1.659761 | -0.95754 | -0.37611 | -1.76834 | -1.14034 | -2.15103 | -2.22629 | 0.967884 | 0.822129 | 1.16105 | 1.63649 | 0.471746 | 0.810365 |
| SPP1 | 0.935136 | 1.924358 | -0.5284 | -0.34871 | -1.92716 | -1.75661 | -1.87206 | -1.8405 | 1.34961 | 0.564351 | 0.772092 | 1.628719 | -0.07693 | 1.176114 |
| THY1 | 1.599879 | 1.88093 | -0.60834 | -1.63361 | -1.84079 | -0.58969 | -1.78377 | -1.27821 | 0.878242 | 0.784798 | 1.606415 | 1.734808 | -0.8514 | 0.10074 |
| FCGR1A | -0.65978 | -0.6701 | 1.189767 | 1.802024 | 2.263395 | 2.07957 | 1.203988 | -0.45319 | -1.06713 | -1.60235 | -1.36588 | -0.39388 | -0.80518 | -1.52127 |
| CD3E | -0.34153 | -0.16733 | 0.936924 | 0.330888 | 2.261328 | 3.416558 | -0.82007 | -1.14727 | 0.351547 | -1.08141 | -1.34201 | -1.3085 | -0.65002 | -0.4391 |
| FXYD6 | 1.863016 | 1.54627 | 0.029334 | -0.80162 | -0.45126 | -1.13634 | -2.30826 | -2.40041 | 1.367929 | -0.28628 | 1.52724 | 1.372797 | -0.23386 | -0.08855 |
| LOC100233156 | 1.209285 | -1.41134 | 1.246733 | -0.99707 | 1.832821 | -1.35243 | 2.036069 | -1.34716 | 0.908655 | -1.42092 | 1.212828 | -0.90501 | 0.5737 | -1.58617 |
| PLIN2 | -0.95001 | -0.75744 | 0.282656 | 0.121997 | 0.68921 | 0.687346 | 3.172016 | 2.544017 | -0.97699 | -1.07487 | -1.22623 | -1.11322 | -1.05016 | -0.34834 |
| LOC339862 | -0.21163 | -0.75956 | 1.896622 | 1.203471 | 2.452781 | 2.073252 | -1.02939 | -1.69203 | 0.134989 | -1.05123 | -1.28102 | -1.35827 | -0.7066 | 0.328606 |
| LOC101928047 | -0.89233 | -0.56125 | -0.82209 | -0.32081 | -0.56225 | 0.024143 | 3.506817 | 2.591861 | -1.41751 | -1.08008 | -0.00047 | -0.66765 | 0.369975 | -0.16838 |
| TNFAIP6 | 0.392655 | -1.02951 | -0.45073 | -1.01552 | -0.71528 | 0.15889 | 3.995372 | 1.61948 | -1.08879 | 0.285035 | -0.64285 | -1.11343 | -0.59272 | 0.197411 |

**Supplemental Table 3.** Top 200 differentially expressed genes in ssBM bulk RNAseq samples

| **Donor** | **1** | **2** | **1** | **2** | **1** | **2** | **1** | **2** | **1** | **2** | **1** | **2** | **1** | **2** |
|---|---|---|---|---|---|---|---|---|---|---|---|---|---|---|
| **Population** | ***a*** | ***a*** | ***b*** | ***b*** | ***c*** | ***c*** | ***d*** | ***d*** | ***e*** | ***e*** | ***f*** | ***f*** | ***g*** | ***g*** |
| **Gene** | | | | | | | | | | | | | | |
| ADD2 | -0.18837 | -0.61622 | -0.34267 | -1.00134 | -1.28685 | -1.16398 | 3.433288 | 2.776148 | -0.52281 | 0.17101 | -0.21869 | -0.40494 | -0.20235 | -0.43222 |
| SPNS3 | -0.00988 | -0.43499 | 1.372769 | 0.820455 | 2.740354 | 2.59142 | -1.13772 | -1.75115 | -0.3251 | -1.13581 | -0.68434 | -0.83804 | -0.37479 | -0.83319 |

**Supplemental Table 4.** Top 200 differential genes in GCSF-mobilized bulk RNAseq samples

| **Donor** | **1** | **2** | **1** | **2** | **1** | **2** | **1** | **2** |
|---|---|---|---|---|---|---|---|---|
| **Population** | ***a*** | ***a*** | ***b*** | ***b*** | ***c*** | ***c*** | ***d*** | ***d*** |
| **Gene** | | | | | | | | |
| XIC | 3.314195 | -3.25311 | 3.190342 | -3.2597 | 3.343936 | -3.25174 | 3.099726 | -3.18365 |
| LOC112268313 | -1.99459 | 3.609558 | -1.98634 | -2.04813 | -2.16378 | 3.984753 | -2.161 | 2.759535 |
| DDX3Y | -2.55706 | 2.629814 | -2.55342 | 2.665592 | -2.80166 | 2.656109 | -2.78868 | 2.749315 |
| KDM5D | -2.06466 | 2.478745 | -2.2529 | 2.439492 | -2.49087 | 1.955178 | -2.30957 | 2.244583 |
| USP9Y | -2.21979 | 2.168112 | -2.21241 | 2.331804 | -2.26503 | 2.110836 | -1.89149 | 1.977969 |
| RPS4Y1 | -2.16189 | 2.136493 | -2.18827 | 2.342526 | -2.20168 | 2.091578 | -2.0387 | 2.019947 |
| HDC | -1.36804 | -1.63502 | -0.50078 | -0.32455 | -1.9356 | -1.37104 | 3.684635 | 3.450414 |
| TXLNGY | -2.05983 | 1.929994 | -2.15391 | 2.29782 | -1.54192 | 1.839596 | -2.33902 | 2.027268 |
| RNA45SN2 | -1.72909 | 1.215871 | -2.72487 | 1.39018 | -2.89706 | 2.072836 | 0.989827 | 1.682302 |
| ZFY | -1.92094 | 1.60998 | -1.91361 | 2.040046 | -1.98172 | 2.29196 | -1.83927 | 1.713565 |
| CNRIP1 | -1.51808 | -0.8909 | -1.5077 | 0.040621 | -1.53063 | -0.77494 | 2.96913 | 3.212501 |
| ANK1 | -0.85227 | -1.29269 | -0.97156 | -0.6687 | -1.41283 | -1.14584 | 3.142865 | 3.201036 |
| DNTT | -1.81667 | -0.84796 | -1.74632 | 0.052773 | 2.390273 | 2.618575 | -2.15821 | 1.507527 |
| KCNH2 | -1.15922 | -1.05978 | -1.16873 | -0.82577 | -1.30122 | -0.71501 | 2.854236 | 3.375495 |
| FCER1A | -1.56754 | -0.75305 | 0.059261 | 0.183244 | -2.00164 | -1.61713 | 2.90177 | 2.795095 |
| GATA1 | -0.88079 | -1.14669 | -0.94191 | -0.82016 | -1.3664 | -0.99849 | 3.278839 | 2.875604 |
| TRIB2 | -2.1305 | -1.51305 | -2.12003 | -1.18848 | 1.685675 | 1.324823 | 1.688248 | 2.253316 |
| IRF8 | -1.45237 | -0.82729 | -1.26784 | -0.69957 | 2.403863 | 3.250988 | -1.59829 | 0.190514 |
| MPO | -2.4371 | -0.57999 | -0.60574 | 0.093628 | 2.164229 | 3.221051 | -0.69535 | -1.16072 |
| ITGB3 | -1.11594 | -1.15515 | -1.16612 | -1.29479 | -0.34167 | -0.73114 | 3.09719 | 2.707627 |
| USP32P1 | -1.9947 | 1.774326 | -1.83514 | 1.878255 | -1.49658 | 1.333885 | -1.37405 | 1.714 |
| XK | -0.92466 | -0.96481 | -1.02775 | -0.59681 | -1.16149 | -0.97779 | 2.715997 | 2.937307 |
| HLA-DRB1 | -1.3369 | 1.963236 | -1.39693 | 1.620881 | -1.53914 | 1.95609 | -2.00797 | 0.740736 |
| MS4A1 | -1.05918 | -0.47493 | -0.89556 | -0.2546 | -1.80124 | -0.71753 | 2.872299 | 2.330741 |
| CD36 | -1.01145 | -0.74701 | -1.1168 | -0.78215 | -0.82748 | -0.84078 | 2.671087 | 2.654591 |
| EIF1AY | -1.36165 | 0.942905 | -1.49596 | 1.681976 | -1.63351 | 1.599974 | -1.55899 | 1.825244 |
| ITGA2B | -0.40164 | -0.31591 | -1.00709 | -0.955 | -1.03632 | -1.4184 | 2.378234 | 2.75613 |
| RYR3 | -0.86953 | 0.168427 | -1.08259 | -0.26453 | -1.64647 | -1.22659 | 2.232723 | 2.688563 |
| UTY | -1.34311 | 1.503974 | -1.7387 | 1.631009 | -1.52894 | 1.417719 | -1.43957 | 1.497619 |
| FREM1 | -0.33231 | 0.312093 | -0.48327 | 0.168175 | -2.15284 | -1.90325 | 2.328436 | 2.062965 |
| TFR2 | -1.23799 | -0.87552 | -0.69227 | -0.16768 | -1.31907 | -0.77334 | 2.435669 | 2.630198 |
| MS4A3 | -1.44398 | -1.31001 | -0.89003 | 0.212781 | -1.21573 | -0.07058 | 2.727499 | 1.99005 |
| LOC112268155 | -1.22059 | 2.103882 | -0.50063 | 1.854397 | -1.34266 | 1.649707 | -1.34066 | -1.20345 |

**Supplemental Table 4.** Top 200 differential genes in GCSF-mobilized bulk RNAseq samples

| **Donor** | **1** | **2** | **1** | **2** | **1** | **2** | **1** | **2** |
|---|---|---|---|---|---|---|---|---|
| **Population** | ***a*** | ***a*** | ***b*** | ***b*** | ***c*** | ***c*** | ***d*** | ***d*** |
| **Gene** | | | | | | | | |
| RNA28SN2 | 0.373334 | 1.419589 | 0.973753 | -0.26335 | 0.703107 | -1.91017 | -2.78306 | 1.486801 |
| ACSM1 | -1.25858 | -1.14071 | -0.03563 | 0.587582 | -0.9842 | -1.54525 | 2.025937 | 2.350855 |
| NOTCH3 | -0.61697 | -0.51395 | -0.85582 | -0.27071 | 2.562301 | 2.091672 | -1.23551 | -1.16101 |
| PLGLB2 | -1.42354 | 0.848175 | -1.4175 | 1.628018 | -1.54558 | 1.413642 | -0.95867 | 1.455453 |
| PRKY | -1.36213 | 1.323606 | -1.58662 | 1.442287 | -1.50874 | 1.270109 | -0.92604 | 1.347533 |
| DYNLL1 | -0.93789 | -0.84264 | -0.69891 | -0.87158 | -0.71177 | -0.58595 | 2.456071 | 2.192683 |
| LTBP1 | -0.66448 | -0.53955 | -0.82958 | -0.50282 | -1.06777 | -0.96724 | 2.124987 | 2.446456 |
| ABO | -1.32549 | 1.106513 | -1.25834 | 1.180834 | -2.21697 | 0.679277 | 0.143371 | 1.690803 |
| SPON1 | -1.14591 | 0.090778 | -0.70451 | 0.230499 | 1.791928 | 2.279786 | -1.80826 | -0.73431 |
| CSF2RB | -0.37072 | -1.03249 | -0.47754 | -0.84557 | -0.77093 | -1.0079 | 2.542573 | 1.962569 |
| JCHAIN | -1.076 | -0.8285 | -0.71986 | -0.67949 | 2.621262 | 1.666455 | -1.18882 | 0.204946 |
| TPSAB1 | -1.04638 | -1.06921 | 0.944437 | -0.94337 | 0.509199 | -1.08966 | 2.874789 | -0.17981 |
| SERF1A | -0.72867 | -0.7637 | -0.7346 | 2.263527 | -0.82929 | -0.77258 | 2.26442 | -0.69909 |
| HBD | -0.70752 | -0.95955 | -0.44396 | -0.47625 | -0.96608 | -0.90908 | 2.187928 | 2.274505 |
| PACSIN1 | -0.99147 | -0.13785 | -0.80699 | -0.44207 | 1.986427 | 2.325185 | -1.33134 | -0.60188 |
| CSF1 | -0.61028 | -0.53519 | -0.72633 | 0.085595 | -1.44196 | -0.90955 | 1.607935 | 2.529788 |
| RAB31 | -0.60821 | -0.21827 | -0.98463 | -0.11369 | 2.373709 | 1.780894 | -1.35669 | -0.8731 |
| LOC105377225 | -0.99987 | 1.172271 | -1.026 | 1.223106 | -0.7281 | -0.93872 | -1.01917 | 2.316485 |
| OVOS | -1.31513 | 1.714331 | -1.54099 | 1.437113 | -1.19779 | 0.556974 | -0.77532 | 1.12082 |
| SPTB | -0.85142 | -0.6086 | -0.81797 | -0.50304 | -0.94913 | -0.57755 | 2.313703 | 1.994011 |
| P2RX5 | -1.50005 | -0.89863 | 0.033919 | 0.584616 | -1.09175 | -0.87475 | 1.561161 | 2.18548 |
| SIK1 | -0.89346 | 1.073182 | -1.65922 | 0.544665 | -1.13687 | 1.856181 | -1.02937 | 1.244905 |
| TTTY15 | -0.98537 | 1.21993 | -1.31649 | 1.219087 | -1.79293 | 1.140911 | -0.72717 | 1.242025 |
| JAML | 1.188086 | 0.537693 | -0.21398 | -0.72902 | 1.871947 | 0.762126 | -1.75489 | -1.66196 |
| MYO16 | -0.87227 | -0.57852 | -0.79318 | -0.65344 | -0.4781 | -0.78214 | 1.933949 | 2.223706 |
| LOC112267940 | -1.34509 | -0.28658 | 1.329324 | 0.318896 | 1.769686 | -1.38815 | 0.930844 | -1.32893 |
| FAM178B | -0.5635 | -0.56048 | -0.70788 | -0.73929 | -0.79801 | -0.7444 | 2.229432 | 1.884109 |
| LOC107987136 | -1.18328 | 1.271401 | -1.09022 | 1.15882 | -1.65507 | 0.459732 | -0.51086 | 1.549472 |
| RNF4 | -0.8722 | 0.891599 | -0.86781 | 1.649707 | -0.94234 | -0.90217 | 1.908191 | -0.86498 |
| SLC2A5 | 0.091522 | 0.129545 | -0.22805 | 0.037977 | 1.74473 | 1.436127 | -2.08794 | -1.12391 |
| SCN3A | -1.22283 | -0.62315 | -0.92346 | -0.50783 | 1.867208 | 1.730861 | -0.91768 | 0.596883 |
| FUT7 | 1.089508 | -1.08458 | 0.958502 | -0.81968 | 1.72548 | 0.590499 | -0.81924 | -1.64049 |
| KEL | -0.55556 | -0.65482 | -0.40348 | -0.84793 | -0.76334 | -0.73102 | 2.058269 | 1.897889 |

**Supplemental Table 4.** Top 200 differential genes in GCSF-mobilized bulk RNAseq samples

| **Donor** | **1** | **2** | **1** | **2** | **1** | **2** | **1** | **2** |
|---|---|---|---|---|---|---|---|---|
| **Population** | ***a*** | ***a*** | ***b*** | ***b*** | ***c*** | ***c*** | ***d*** | ***d*** |
| **Gene** | | | | | | | | |
| F13A1 | -1.47941 | -0.80178 | 0.473692 | -0.28004 | 1.771841 | 1.744547 | -0.86764 | -0.56121 |
| CCR7 | -0.54479 | -0.42952 | -0.66394 | -0.84985 | 2.215548 | 1.624089 | -0.99252 | -0.35901 |
| LOC107984312 | -0.69842 | -0.89823 | -0.80266 | -0.33033 | -0.72427 | -0.37762 | 1.697987 | 2.133544 |
| ZNF385D | 0.231372 | 0.229322 | 0.518645 | 0.591358 | -2.14701 | -1.57326 | 1.039534 | 1.11004 |
| SUCNR1 | -0.96192 | -0.69242 | -0.2489 | -0.04257 | 2.052821 | 1.679961 | -0.9077 | -0.87927 |
| PIEZO2 | -0.8742 | -0.56216 | -0.80194 | -0.72563 | 0.014943 | -0.78594 | 1.5224 | 2.212533 |
| ACSM4 | -0.75347 | -0.77025 | -0.21329 | -0.44365 | -0.83747 | -0.78523 | 2.074177 | 1.729187 |
| RASSF6 | 1.534195 | 1.464854 | 0.51125 | 0.530051 | -0.2637 | -0.99675 | -1.47271 | -1.30719 |
| AFF2 | -1.19999 | -1.10051 | -0.78624 | 0.154231 | -0.58469 | -0.03046 | 1.611663 | 1.935989 |
| SPTA1 | -0.83479 | -1.10806 | -0.36321 | -0.43541 | -0.41711 | -0.57255 | 1.679025 | 2.052108 |
| MINPP1 | -0.99409 | -0.88984 | -0.32695 | -0.62469 | -0.30803 | -0.58965 | 1.975101 | 1.758156 |
| SLC40A1 | -0.51091 | -0.14526 | -0.36511 | -0.00949 | -1.15725 | -1.31222 | 1.673506 | 1.826738 |
| SLC24A3 | -0.95451 | -0.99728 | -0.44129 | 0.064152 | -0.78542 | -0.50868 | 1.687535 | 1.935497 |
| LOC101928202 | 0.980629 | 1.979313 | -0.37253 | 0.978265 | -1.28015 | -0.71465 | -1.07022 | -0.50066 |
| SPNS3 | -0.11989 | 0.083137 | 0.104355 | 0.063383 | 1.400873 | 1.56562 | -1.81608 | -1.28139 |
| NTRK1 | -0.73551 | -0.60793 | -0.73152 | -0.25823 | -0.81644 | -0.5513 | 1.960837 | 1.740074 |
| RIPOR3 | -0.01221 | 0.517564 | -0.35266 | 0.094829 | -1.75502 | -1.32639 | 1.347682 | 1.486216 |
| STXBP6 | -0.67772 | -0.52507 | -0.71519 | -0.39999 | -0.65863 | -0.72205 | 1.7503 | 1.948355 |
| HOPX | 0.866011 | 0.596295 | 0.553409 | 0.55635 | 0.970278 | 0.062498 | -1.65334 | -1.9515 |
| SPTBN2 | -1.16173 | -0.27047 | -0.69026 | -0.10437 | -1.06432 | -0.17763 | 1.581574 | 1.887205 |
| FAM215B | -1.89653 | 0.733467 | -1.42234 | 1.177185 | -0.09941 | 1.166778 | -0.06376 | 0.40461 |
| NBL1 | 1.477117 | 0.679291 | 0.832273 | 0.106134 | -2.12824 | 0.15031 | -1.06316 | -0.05373 |
| SETBP1 | 0.9435 | 0.617085 | 0.280864 | 0.0273 | 1.147055 | 0.478494 | -1.83869 | -1.6556 |
| CPA3 | -1.72363 | -1.6534 | 0.270344 | 0.348766 | 0.328662 | -0.01516 | 1.289588 | 1.154831 |
| CRYBG1 | -0.03144 | 0.380532 | 0.087373 | 0.537977 | 1.147508 | 1.148494 | -2.12559 | -1.14485 |
| PRG2 | 0.409765 | -0.74533 | -0.7677 | -1.11366 | -0.49297 | -0.43682 | 2.238392 | 0.908322 |
| LOC112268349 | -1.59623 | 0.49608 | 0.23005 | 0.383601 | -0.1788 | 1.19943 | -1.70759 | 1.173453 |
| PDZRN4 | -0.55773 | 1.990102 | -0.45592 | 1.334879 | -1.0654 | -0.84112 | -0.67799 | 0.273186 |
| MAMDC2 | 0.184867 | 0.041883 | 0.876935 | 0.825513 | 0.855116 | 0.658171 | -1.69754 | -1.74494 |
| PLGLB1 | 0.945972 | 0.008785 | 1.106954 | -1.28139 | 0.74583 | -1.28739 | 0.997676 | -1.23644 |
| CP | -1.2171 | -0.62283 | 1.185624 | 1.166588 | 1.022667 | 0.631273 | -0.97028 | -1.19594 |
| MME | -0.78953 | -0.69777 | -1.30732 | -0.27193 | 1.399987 | 1.612618 | -0.67459 | 0.728531 |
| SLC5A4-AS1 | 1.535696 | -1.00296 | 0.913888 | -1.01365 | 1.239164 | -0.84271 | 0.142651 | -0.97209 |

**Supplemental Table 4.** Top 200 differential genes in GCSF-mobilized bulk RNAseq samples

| **Donor** | **1** | **2** | **1** | **2** | **1** | **2** | **1** | **2** |
|---|---|---|---|---|---|---|---|---|
| **Population** | ***a*** | ***a*** | ***b*** | ***b*** | ***c*** | ***c*** | ***d*** | ***d*** |
| **Gene** | | | | | | | | |
| LOC105378072 | -0.06772 | 0.195957 | -0.24535 | 0.694347 | 1.025383 | 1.296435 | -2.11843 | -0.78062 |
| SDK2 | -0.37797 | 0.250552 | -0.10443 | 0.376729 | 1.355911 | 1.33369 | -1.86178 | -0.9727 |
| GPR75-ASB3 | -1.18244 | 0.291933 | 1.624032 | 0.65936 | 0.889774 | 0.166779 | -1.27958 | -1.16986 |
| LOC105379461 | -0.99709 | -0.68833 | -0.55012 | 0.099627 | -0.49826 | -0.7569 | 1.697871 | 1.693204 |
| SH3BP5 | 0.61995 | 0.010953 | -0.30561 | -0.27506 | 1.181118 | 1.570466 | -1.39408 | -1.40774 |
| NA | 1.177053 | -0.64409 | 1.101967 | -1.10283 | 1.335449 | -0.59984 | 0.120855 | -1.38856 |
| NPTX2 | -0.298 | -0.57738 | -0.01993 | -0.5417 | 1.955223 | 1.303913 | -1.32626 | -0.49586 |
| PTGS1 | 0.83361 | 0.276263 | 0.236273 | -0.34761 | -1.30812 | -1.80316 | 1.234547 | 0.878193 |
| HCK | 0.017235 | -0.64878 | 0.750197 | 0.070035 | 1.531289 | 1.021718 | -1.42995 | -1.31175 |
| PRKG2 | 0.871968 | 0.525587 | 0.580524 | 0.392705 | -1.48327 | -1.9485 | 0.601454 | 0.459537 |
| VWF | 0.341327 | -0.20152 | 0.541455 | -0.69443 | -1.12493 | -1.51915 | 1.367124 | 1.290127 |
| LIMCH1 | 1.437242 | 1.097935 | 0.641399 | 0.571941 | -0.87836 | -1.448 | -0.44265 | -0.9795 |
| SPINK2 | 0.325724 | 0.258672 | 0.943734 | 0.580327 | 0.966037 | 0.258666 | -1.52016 | -1.813 |
| CYTH4 | 0.557251 | 0.039773 | 0.804742 | 0.03348 | 1.289768 | 0.404746 | -1.17044 | -1.95932 |
| UNC5B-AS1 | 1.450601 | -0.474 | 0.088976 | -0.84772 | 1.503837 | 0.458678 | -1.04817 | -1.1322 |
| KBTBD11 | 0.127632 | -1.00879 | 0.77032 | -0.22947 | 1.750952 | 0.740387 | -0.70104 | -1.44999 |
| ADAMTS14 | -0.84467 | -0.90089 | -0.13216 | 0.014331 | -0.91476 | -0.44978 | 1.67935 | 1.548584 |
| LINC00865 | -0.64532 | -0.04279 | -0.65997 | -0.41503 | 0.860125 | 1.96199 | -1.40908 | 0.350066 |
| GSTM1 | -0.80255 | 0.832273 | -0.92665 | 0.865546 | -0.61608 | 1.162455 | -1.44588 | 0.930885 |
| SPNS2 | 0.934812 | 0.318602 | 0.415862 | 0.151896 | 0.70302 | 0.729386 | -1.40622 | -1.84736 |
| LOC112268350 | 0.741811 | 1.012595 | -1.21084 | 0.832284 | 0.28889 | -1.25351 | 0.793469 | -1.2047 |
| GSTM5 | 0.940638 | -0.91062 | 0.904484 | -1.2126 | 1.067215 | -1.03703 | 0.905411 | -0.6575 |
| DHRS3 | -0.07178 | -0.79459 | -0.8855 | -1.01913 | -0.05348 | -0.28971 | 1.713817 | 1.400375 |
| RNA28SN5 | 0.814476 | -0.50671 | 0.28411 | -1.82209 | -0.30444 | -0.66127 | 0.804773 | 1.391147 |
| BLOC1S5-TXNDC5 | 1.296534 | -0.37446 | 0.504422 | 0.36931 | -1.54269 | -1.42834 | 0.801677 | 0.373551 |
| MICAL2 | -0.76488 | -0.83822 | -0.07958 | -0.67485 | -0.41338 | -0.47419 | 1.597736 | 1.647347 |
| MMP2 | 0.695599 | 0.370651 | 0.84848 | 0.088823 | 1.075598 | 0.014911 | -1.40141 | -1.69266 |
| MIR8485 | -0.98871 | -0.36666 | -0.63839 | -0.4866 | 1.930676 | -0.33607 | -0.33664 | 1.222394 |
| LOC107984427 | -0.31485 | 1.317403 | -1.24526 | 0.891416 | -1.32974 | 0.612043 | -0.62137 | 0.690371 |
| HLA-DQA2 | 0.914159 | -0.06243 | 1.142278 | -0.46075 | 1.21942 | -0.4415 | -0.67376 | -1.63742 |
| ARHGEF12 | 0.471931 | 0.419538 | -0.06992 | -0.08812 | -1.45061 | -1.48344 | 1.124699 | 1.075917 |
| ARL4C | 0.505835 | -0.40284 | 0.555491 | -0.23916 | 1.220649 | 1.075345 | -1.447 | -1.26832 |
| EPPK1 | 1.866131 | 1.239733 | 0.07257 | -0.58135 | -0.45754 | -0.80209 | -0.74315 | -0.5943 |

**Supplemental Table 4.** Top 200 differential genes in GCSF-mobilized bulk RNAseq samples

| **Donor** | **1** | **2** | **1** | **2** | **1** | **2** | **1** | **2** |
|---|---|---|---|---|---|---|---|---|
| **Population** | ***a*** | ***a*** | ***b*** | ***b*** | ***c*** | ***c*** | ***d*** | ***d*** |
| **Gene** | | | | | | | | |
| PDE3A | -0.08887 | -0.04991 | -0.37153 | 0.051105 | -1.23214 | -1.16643 | 1.460742 | 1.397028 |
| HLA-DRB3 | -0.81756 | 1.038228 | -0.75211 | 1.181796 | -0.91837 | 1.208137 | -1.02697 | 0.086854 |
| LOC112267939 | 0.747648 | 1.177519 | -1.12907 | -1.10778 | -1.22513 | 0.875364 | 0.120328 | 0.541118 |
| STAB1 | 0.647993 | 0.407916 | -0.81669 | -0.61436 | 0.713361 | 1.680969 | -0.99352 | -1.02567 |
| TIMP3 | -0.61908 | -0.24541 | -0.85895 | -0.4052 | -0.92116 | 0.001686 | 1.750773 | 1.29735 |
| AKAP12 | -0.45218 | -0.47136 | -0.53856 | -0.147 | -0.8503 | 0.087444 | 0.055844 | 2.316109 |
| NPIPA3 | -1.19253 | 0.191683 | -0.9758 | 0.667737 | 0.467227 | 1.774826 | -0.69978 | -0.23337 |
| ST6GAL2 | 0.385738 | 0.24194 | -0.04943 | -0.42458 | -1.12368 | -1.46516 | 1.257004 | 1.178155 |
| HNRNPLL | 0.433216 | -0.04233 | 0.448732 | 0.131582 | -1.263 | -1.66956 | 1.058893 | 0.902467 |
| MEG3 | 1.349487 | 1.408629 | 0.143697 | 0.360292 | -0.94337 | -0.80152 | -0.55393 | -0.96329 |
| HOXA10-HOXA9 | -0.91612 | -0.42736 | 1.434086 | -0.94234 | 1.267308 | -0.94722 | 0.064671 | 0.466983 |
| TSPOAP1 | -0.92118 | -0.25304 | -0.30844 | -0.09884 | 1.185515 | 1.787159 | -0.88171 | -0.50946 |
| LOC100130264 | -0.68702 | -0.60505 | -0.46322 | -0.13754 | -0.49491 | -0.6794 | 1.744901 | 1.322236 |
| TCEAL2 | 1.887009 | 0.869674 | 0.124708 | -0.20698 | -0.83962 | -0.98206 | -0.14905 | -0.70368 |
| ID3 | 1.413329 | 0.815491 | 0.893528 | 0.133242 | -0.53996 | -1.2611 | -0.45973 | -0.9948 |
| FYB1 | -1.15191 | -0.86401 | -1.00576 | -0.5049 | 0.728222 | 0.807556 | 1.154734 | 0.836073 |
| NEGR1 | 0.347185 | 0.35994 | 0.05595 | 0.110531 | 1.106632 | 0.8899 | -1.71246 | -1.15768 |
| MICALCL | -0.52517 | -0.88642 | -0.39836 | -0.33406 | -0.65612 | -0.25568 | 1.519641 | 1.536182 |
| AVP | 1.398494 | 1.138341 | 0.549391 | -0.07223 | -0.13695 | -0.82281 | -0.81933 | -1.23491 |
| SELP | 0.823054 | 0.625494 | -0.05679 | -0.2616 | -1.31314 | -1.47166 | 0.982548 | 0.672093 |
| CDH2 | 0.402368 | 0.614045 | -0.1772 | 0.700737 | 0.586134 | 0.789202 | -1.67226 | -1.24303 |
| LOC112268061 | 1.029674 | -0.34859 | 0.592291 | -0.63797 | 1.281384 | 0.369751 | -1.02914 | -1.25739 |
| FNTB | -1.59468 | 0.60329 | -0.09651 | 0.200609 | -1.2234 | 0.259216 | 1.173005 | 0.678478 |
| GPRIN3 | 0.278154 | 0.531476 | -0.26915 | 0.474045 | 0.877926 | 0.946123 | -1.46564 | -1.37294 |
| COBLL1 | 0.157503 | -0.10024 | 0.088631 | 0.132454 | 1.254821 | 1.108314 | -1.28698 | -1.3545 |
| PCDH17 | 1.711322 | 0.672051 | 0.479788 | -0.46594 | -1.06972 | -0.99617 | 0.233439 | -0.56477 |
| KIAA0087 | -0.44685 | 0.040402 | -0.24202 | -0.04379 | 1.259526 | 1.514977 | -1.1275 | -0.95475 |
| NA | -0.50606 | -0.51742 | -0.50304 | -0.52487 | 1.523475 | -0.5279 | 1.555203 | -0.49939 |
| RGS18 | 0.022899 | -0.15922 | 0.096405 | -0.14218 | -1.38195 | -0.97702 | 0.983387 | 1.557687 |
| GRAMD1C | 0.665039 | 0.335771 | 0.471461 | 0.324009 | 0.810498 | 0.372089 | -1.75132 | -1.22754 |
| ZBTB16 | -0.2721 | -0.4842 | 0.46044 | 0.212262 | -1.1853 | -1.1535 | 1.274786 | 1.14761 |
| EPAS1 | 1.387005 | 1.263302 | 0.201107 | 0.272066 | -0.45543 | -0.84685 | -0.81019 | -1.01101 |
| LTB | 0.095009 | 0.206057 | -0.63469 | -0.21055 | 1.240319 | 1.31735 | -1.45732 | -0.55618 |

**Supplemental Table 4.** Top 200 differential genes in GCSF-mobilized bulk RNAseq samples

| **Donor** | **1** | **2** | **1** | **2** | **1** | **2** | **1** | **2** |
|---|---|---|---|---|---|---|---|---|
| **Population** | ***a*** | ***a*** | ***b*** | ***b*** | ***c*** | ***c*** | ***d*** | ***d*** |
| **Gene** | | | | | | | | |
| LEF1 | -0.34624 | -0.37703 | -0.49595 | -0.4629 | -0.6984 | -0.50932 | 0.916201 | 1.973629 |
| RNA45SN5 | -0.95124 | 0.904127 | -0.01545 | -0.61675 | -0.05573 | -1.20613 | 0.354777 | 1.586397 |
| IQSEC3 | 1.314141 | -0.7592 | 0.854741 | -1.03251 | 0.547514 | -0.77427 | 0.696314 | -0.84673 |
| KALRN | -0.46751 | -0.25402 | -0.79228 | -0.44727 | -0.39557 | -0.6454 | 1.46191 | 1.540135 |
| LOC100996709 | -0.11352 | 0.774715 | -1.45157 | 0.779265 | -0.10983 | 1.008753 | -1.28931 | 0.401488 |
| AGTR1 | -0.50074 | -0.51173 | -0.49802 | -0.51824 | -0.55521 | -0.43973 | 1.373121 | 1.650542 |
| TMEM72 | -0.37977 | 0.133088 | -0.49515 | -0.11893 | -0.2844 | -0.32532 | -0.75511 | 2.225603 |
| THBS1 | 0.022038 | -1.00517 | -0.64086 | -0.81412 | -0.41921 | 0.093195 | 1.241866 | 1.522264 |
| FIGN | -0.78944 | 1.372705 | -0.76276 | 1.212149 | -0.98248 | 0.13934 | -0.62673 | 0.437216 |
| TNFAIP2 | 0.291258 | 0.582202 | -0.08382 | 0.125128 | 0.853006 | 1.01973 | -1.38459 | -1.40291 |
| MYO5C | 0.538276 | 0.364259 | 0.574284 | 0.503634 | 0.560181 | 0.472503 | -1.49038 | -1.52275 |
| LOC107985787 | 1.540374 | 0.924221 | 0.241843 | -0.07339 | -1.11259 | -1.19349 | -0.07946 | -0.24751 |
| LOC105378305 | 0.543928 | 0.039541 | 0.281977 | 0.677332 | 0.884152 | 0.463348 | -1.43012 | -1.46016 |
| LRP1 | 1.490136 | 1.299807 | -0.15317 | 0.137134 | -0.89026 | -0.70081 | -0.77881 | -0.40402 |
| FCMR | 0.591449 | -0.09034 | -0.24741 | -0.08689 | 1.378314 | 0.796038 | -1.53118 | -0.80999 |
| LOC105374809 | 1.018116 | -0.8433 | 0.418436 | -0.85191 | 1.031083 | -0.85622 | 0.902102 | -0.8183 |
| ZFPM1 | 0.118863 | 0.077221 | -0.34137 | -0.0401 | -1.28361 | -1.0168 | 1.174298 | 1.311499 |
| MYCT1 | 0.78626 | 0.528502 | 0.368411 | 0.26518 | -1.23999 | -1.67244 | 0.539792 | 0.424289 |
| GIMAP1-GIMAP5 | 1.195392 | 0.728211 | 0.82313 | -1.0191 | 0.395559 | -0.09558 | -1.05961 | -0.96799 |
| FAM83D | -0.59063 | -0.51025 | -0.81102 | -0.19386 | -0.4199 | -0.37332 | 1.637323 | 1.261651 |
| FSBP | -0.2607 | 0.759779 | 0.931491 | 0.820775 | -1.37373 | -0.07347 | 0.461096 | -1.26524 |
| GATA2 | 0.24215 | -0.39201 | 0.248592 | -0.33094 | -0.75987 | -1.39841 | 1.438319 | 0.952173 |
| LOC107986265 | 0.966322 | -0.58027 | 0.299032 | -1.82085 | 1.09429 | -0.07049 | 0.029977 | 0.081988 |
| LOC107987373 | -0.79482 | 0.502942 | -0.79128 | 0.643854 | -0.86507 | 1.296263 | -0.86396 | 0.87208 |
| TTTY10 | -0.78205 | 1.06724 | -0.77855 | 1.139131 | -0.85148 | 0.22748 | -0.85037 | 0.828602 |
| CTNNBL1 | -0.49804 | -0.87134 | -0.0207 | -0.18752 | -0.58453 | -0.64602 | 1.60546 | 1.202691 |
| ITGAL | -0.37088 | -0.25738 | 0.184555 | 0.082067 | 1.198333 | 1.332391 | -1.00415 | -1.16494 |
| SIGLEC17P | -0.72145 | -0.31104 | -0.16037 | -0.0722 | 1.659992 | 1.098307 | -0.573 | -0.92024 |
| PREX2 | 1.363727 | 1.219626 | 0.33524 | 0.060045 | -0.99591 | -0.60374 | -0.60763 | -0.77136 |
| FGD5 | 1.103393 | 0.840337 | 0.87704 | 0.450192 | -0.62903 | -0.87389 | -0.6002 | -1.16785 |
| RTEL1-TNFRSF6B | 0.461359 | 0.685096 | 1.16472 | -0.59328 | 0.810399 | -1.12471 | -1.17619 | -0.22739 |
| MN1 | 0.335467 | 0.282844 | 0.394402 | 0.054714 | 1.073282 | 0.629183 | -1.40763 | -1.36226 |
| C10orf105 | -0.21408 | 0.009205 | -0.60145 | -0.27215 | 1.375575 | 1.387218 | -0.88346 | -0.80086 |

**Supplemental Table 4.** Top 200 differential genes in GCSF-mobilized bulk RNAseq samples

| **Donor** | **1** | **2** | **1** | **2** | **1** | **2** | **1** | **2** |
|---|---|---|---|---|---|---|---|---|
| **Population** | ***a*** | ***a*** | ***b*** | ***b*** | ***c*** | ***c*** | ***d*** | ***d*** |
| **Gene** | | | | | | | | |
| SEPT5-GP1BB | 0.172402 | 0.365396 | -0.0481 | -0.1582 | -1.33376 | -1.16487 | 0.941865 | 1.225272 |
| MFSD2B | -0.90678 | -0.78637 | 0.075083 | 0.072034 | -0.78371 | -0.31473 | 1.261499 | 1.382977 |

**Supplemental Table 5.** Genes upregulated in ssBM cells from Population *a*

| Gene | baseMean | log2FoldChange | pvalue | padj |
|---|---|---|---|---|
| MX1 | 494.9739 | -1.00059 | 0.00011 | 0.005876 |
| SWAP70 | 1695.026 | -1.00516 | 0.000263 | 0.011562 |
| LOC100128108 | 394.85 | -1.00781 | 1.53E-05 | 0.001281 |
| NRIP1 | 10229.62 | -1.01157 | 0.000693 | 0.024322 |
| RAB29 | 585.3915 | -1.02144 | 0.001366 | 0.039716 |
| STX3 | 463.4232 | -1.02365 | 9.94E-05 | 0.005453 |
| YES1 | 326.9988 | -1.03125 | 0.001784 | 0.047559 |
| ELMO1 | 4476.965 | -1.03125 | 0.000663 | 0.023531 |
| LRRC70 | 1523.709 | -1.03374 | 0.000809 | 0.027297 |
| IL4R | 180.2649 | -1.03733 | 0.000472 | 0.018068 |
| ZC2HC1A | 355.8907 | -1.03842 | 0.001823 | 0.047849 |
| CD109 | 1587.164 | -1.0417 | 2.30E-05 | 0.001836 |
| JUND | 24583.97 | -1.04277 | 5.14E-06 | 0.000557 |
| VIM | 24350.85 | -1.04514 | 1.02E-06 | 0.000144 |
| PNPLA8 | 632.7647 | -1.05394 | 0.000809 | 0.027297 |
| LOC102724765 | 659.8099 | -1.05476 | 0.000153 | 0.007603 |
| NFAT5 | 988.4713 | -1.05822 | 1.24E-05 | 0.00108 |
| PHTF1 | 2629.221 | -1.06274 | 5.02E-06 | 0.000548 |
| NFATC2 | 988.6014 | -1.06412 | 2.86E-05 | 0.002133 |
| ITPKB | 590.1405 | -1.06489 | 0.000168 | 0.008073 |
| LINC01122 | 563.7154 | -1.09004 | 3.41E-05 | 0.002433 |
| TRIM8 | 853.5996 | -1.09707 | 7.59E-06 | 0.000744 |
| LOC105378645 | 489.4958 | -1.10043 | 0.000694 | 0.024322 |
| NEDD4L | 897.8868 | -1.10229 | 0.000217 | 0.00995 |
| MMP28 | 374.8082 | -1.10399 | 0.001431 | 0.041202 |
| GBP5 | 253.9217 | -1.10458 | 0.001087 | 0.033761 |
| LOC105375304 | 523.8434 | -1.1046 | 0.001802 | 0.047627 |
| NAMPT | 1612.35 | -1.10539 | 0.000166 | 0.008047 |
| GCHFR | 1091.951 | -1.10858 | 0.000181 | 0.008481 |
| CALN1 | 1219.665 | -1.10867 | 7.52E-07 | 0.000116 |
| SORBS3 | 735.406 | -1.11083 | 0.001758 | 0.047231 |
| GABARAPL1 | 342.5218 | -1.11132 | 0.000636 | 0.022842 |
| CHMP1B | 7120.283 | -1.11134 | 0.000152 | 0.007588 |
| C3orf80 | 921.1874 | -1.11415 | 0.000264 | 0.0116 |
| HOPX | 4900.006 | -1.1144 | 0.001462 | 0.041589 |
| LINC-PINT | 644.1453 | -1.11475 | 0.000194 | 0.008969 |
| DST | 1388.623 | -1.11724 | 1.58E-06 | 0.000212 |
| BMPR2 | 296.5071 | -1.11791 | 0.000936 | 0.030133 |
| GATA2-AS1 | 314.0654 | -1.12577 | 0.000533 | 0.019843 |
| HSPA1B | 337.4805 | -1.12968 | 0.000362 | 0.014716 |
| LOC100507507 | 537.8228 | -1.13592 | 0.000177 | 0.008348 |
| UBOX5 | 375.1842 | -1.13951 | 3.68E-05 | 0.002543 |
| INPP4B | 758.1114 | -1.1402 | 0.001269 | 0.037789 |
| EVA1C | 283.4484 | -1.14174 | 0.000748 | 0.025547 |
| STK17B | 4280.73 | -1.14304 | 1.90E-05 | 0.001557 |
| RNF19B | 305.595 | -1.15395 | 0.001791 | 0.047627 |
| RANBP2 | 2437.924 | -1.15717 | 1.07E-05 | 0.00098 |
| HLA-E | 12294.85 | -1.15833 | 0.001804 | 0.047627 |
| CEBPB | 604.3898 | -1.15873 | 0.001362 | 0.039716 |
| GSTO2 | 612.8316 | -1.15995 | 0.001126 | 0.034731 |
| TNFAIP3 | 2709.294 | -1.16039 | 1.66E-05 | 0.001384 |
| YPEL5 | 10498.52 | -1.16157 | 0.000172 | 0.00819 |
| NOXA1 | 181.4893 | -1.17536 | 0.000179 | 0.008363 |
| PRKCH | 1707.82 | -1.17933 | 0.001844 | 0.048329 |

**Supplemental Table 5.** Genes upregulated in ssBM cells from Population *a*

| Gene | baseMean | log2FoldChange | pvalue | padj |
|---|---|---|---|---|
| ARHGEF40 | 283.8867 | -1.18677 | 5.50E-05 | 0.003403 |
| HMGA2 | 513.254 | -1.18805 | 2.05E-06 | 0.000265 |
| VPS37B | 2237.233 | -1.19358 | 0.001451 | 0.041452 |
| ANKRD42 | 233.6819 | -1.19399 | 0.001064 | 0.0332 |
| FTH1 | 147380.8 | -1.19638 | 3.87E-07 | 6.57E-05 |
| TK2 | 213.6409 | -1.2 | 0.000589 | 0.021495 |
| MPPED2 | 236.694 | -1.20613 | 0.001298 | 0.038335 |
| VWA2 | 254.6622 | -1.20687 | 0.001448 | 0.041449 |
| RHOC | 767.2825 | -1.20768 | 0.000143 | 0.00719 |
| LDOC1 | 615.5366 | -1.21206 | 4.12E-06 | 0.000484 |
| PRDM16 | 205.8275 | -1.2148 | 0.001741 | 0.046894 |
| IDS | 3484.561 | -1.21505 | 0.001133 | 0.034783 |
| TIMP3 | 810.5746 | -1.21723 | 0.000161 | 0.007909 |
| TSPYL2 | 454.4889 | -1.21757 | 3.33E-06 | 0.000403 |
| SNAPC1 | 905.6843 | -1.2268 | 8.79E-05 | 0.004909 |
| LOC105377458 | 124.7842 | -1.22857 | 0.001857 | 0.048607 |
| HEMGN | 1259.552 | -1.23134 | 6.88E-06 | 0.000681 |
| FOSL2 | 3806.124 | -1.23237 | 1.10E-05 | 0.000992 |
| SNHG14 | 164.7249 | -1.23668 | 6.18E-05 | 0.00376 |
| EHD2 | 745.9003 | -1.24887 | 0.00113 | 0.034783 |
| NFIL3 | 1937.509 | -1.2516 | 0.000202 | 0.009311 |
| KLHL3 | 205.1869 | -1.25654 | 0.000726 | 0.0251 |
| FZD1 | 204.8696 | -1.26143 | 0.000676 | 0.023838 |
| CAVIN1 | 1759.025 | -1.26476 | 4.15E-05 | 0.002767 |
| LOC101929709 | 379.9723 | -1.26889 | 0.001206 | 0.036522 |
| LOC107984658 | 218.7394 | -1.27046 | 0.000437 | 0.017106 |
| MYH10 | 926.3862 | -1.27172 | 0.000165 | 0.008019 |
| PPP1R9A | 379.9274 | -1.27235 | 0.001764 | 0.047248 |
| NKAIN2 | 466.6337 | -1.27338 | 4.77E-05 | 0.003065 |
| GCH1 | 484.7873 | -1.28454 | 0.000648 | 0.02319 |
| H1F0 | 6554.085 | -1.28929 | 0.000302 | 0.012885 |
| ADGRG6 | 1003.141 | -1.30231 | 2.24E-07 | 4.06E-05 |
| PXDC1 | 154.1464 | -1.30421 | 0.000615 | 0.022369 |
| RIN2 | 111.4428 | -1.30489 | 0.001399 | 0.040501 |
| NFIA | 1541.033 | -1.30615 | 7.62E-05 | 0.004388 |
| FBXO44 | 109.8309 | -1.30759 | 0.00052 | 0.019435 |
| LOC105369593 | 983.6716 | -1.30809 | 4.42E-06 | 0.000509 |
| MIR155HG | 289.1947 | -1.32019 | 3.96E-05 | 0.002669 |
| RRN3P1 | 81.70608 | -1.32093 | 0.00147 | 0.041761 |
| CREM | 1291.098 | -1.32601 | 5.28E-05 | 0.003327 |
| LOC107985939 | 127.5152 | -1.32979 | 0.001475 | 0.041825 |
| SNX21 | 127.0076 | -1.3306 | 0.00182 | 0.047838 |
| FAM133A | 186.3463 | -1.33188 | 0.001271 | 0.037789 |
| BEX5 | 481.6012 | -1.33343 | 0.000354 | 0.014563 |
| LOC105371227 | 389.2573 | -1.33936 | 8.85E-07 | 0.000132 |
| VEGFA | 400.5256 | -1.34238 | 0.000252 | 0.011165 |
| SNX9 | 515.0679 | -1.34693 | 7.51E-05 | 0.004351 |
| CXCL3 | 1211.923 | -1.3567 | 5.67E-08 | 1.23E-05 |
| ABCA13 | 215.3193 | -1.36217 | 0.000944 | 0.03027 |
| TNFRSF1B | 540.4958 | -1.36423 | 5.72E-08 | 1.23E-05 |
| FERMT1 | 141.1021 | -1.36618 | 8.48E-05 | 0.004838 |
| RABGEF1 | 1110.413 | -1.3663 | 0.000327 | 0.013565 |
| HIST1H2BK | 1553.965 | -1.36644 | 2.99E-05 | 0.002212 |
| SGIP1 | 99.91713 | -1.37246 | 0.001497 | 0.042196 |

**Supplemental Table 5.** Genes upregulated in ssBM cells from Population *a*

| Gene | baseMean | log2FoldChange | pvalue | padj |
|---|---|---|---|---|
| PHF1 | 327.5009 | -1.37558 | 0.00011 | 0.005872 |
| ST6GAL2 | 589.8345 | -1.38018 | 0.000163 | 0.007944 |
| HCG4 | 147.0901 | -1.38125 | 0.000717 | 0.02489 |
| NR4A3 | 2236.945 | -1.38234 | 0.000124 | 0.006393 |
| SHTN1 | 232.4562 | -1.38615 | 0.000267 | 0.011707 |
| ATP2B1-AS1 | 1077.006 | -1.3867 | 3.51E-09 | 1.01E-06 |
| LINC02265 | 121.8383 | -1.38957 | 0.001439 | 0.041393 |
| ZNF589 | 190.2328 | -1.40478 | 0.00182 | 0.047838 |
| LRP10 | 555.1695 | -1.40486 | 1.42E-05 | 0.001203 |
| CSRNP1 | 1948.591 | -1.40699 | 1.43E-05 | 0.001203 |
| RBPMS | 1174.029 | -1.41199 | 3.59E-07 | 6.20E-05 |
| SCN9A | 792.4347 | -1.41636 | 2.00E-05 | 0.001616 |
| SERPINI1 | 472.1499 | -1.41824 | 0.000171 | 0.00819 |
| BIRC3 | 1303.024 | -1.42029 | 4.11E-11 | 1.88E-08 |
| STARD9 | 325.7854 | -1.42139 | 5.77E-06 | 0.000597 |
| ADAM28 | 1719.033 | -1.42905 | 0.000396 | 0.015776 |
| ROBO4 | 389.2318 | -1.42986 | 0.00022 | 0.010056 |
| BEST1 | 154.64 | -1.43627 | 0.000427 | 0.016756 |
| LOC284454 | 2061.464 | -1.44026 | 5.28E-09 | 1.47E-06 |
| MYCT1 | 972.639 | -1.44618 | 5.53E-06 | 0.000583 |
| CXCL2 | 4955.58 | -1.44788 | 4.84E-09 | 1.36E-06 |
| CXorf40A | 695.809 | -1.44978 | 9.99E-06 | 0.000929 |
| DUBR | 163.005 | -1.45581 | 0.001718 | 0.0467 |
| LOC729291 | 85.08995 | -1.45854 | 0.001546 | 0.043099 |
| MXRA7 | 237.646 | -1.46318 | 0.000745 | 0.025547 |
| HIST1H1C | 3203.13 | -1.46322 | 3.30E-09 | 9.61E-07 |
| ABCB1 | 543.9984 | -1.46333 | 0.000361 | 0.014716 |
| SLC2A3 | 3750.683 | -1.48391 | 0.000299 | 0.012814 |
| SLC22A17 | 158.4224 | -1.49368 | 0.000309 | 0.013147 |
| ZC3H12A | 1196.588 | -1.49494 | 4.68E-06 | 0.000522 |
| ESAM | 381.9372 | -1.49951 | 0.000802 | 0.02717 |
| DCUN1D3 | 162.016 | -1.50068 | 0.00026 | 0.011497 |
| C16orf45 | 384.4037 | -1.51012 | 0.000104 | 0.005616 |
| PRKG2 | 1050.742 | -1.5134 | 3.71E-08 | 8.18E-06 |
| ITGAM | 125.8974 | -1.51345 | 0.000318 | 0.013333 |
| AREG | 22159.91 | -1.51493 | 8.14E-06 | 0.000785 |
| LOC100506282 | 96.81404 | -1.51911 | 0.000287 | 0.012479 |
| GADD45A | 2369.762 | -1.51987 | 1.33E-05 | 0.001139 |
| TNFSF8 | 277.8283 | -1.52146 | 0.001262 | 0.037704 |
| PLAG1 | 201.9516 | -1.52443 | 0.001692 | 0.046105 |
| CCL4L2 | 113.6835 | -1.52466 | 0.001275 | 0.037839 |
| NEIL1 | 136.1296 | -1.52534 | 0.000601 | 0.0219 |
| NFIB | 514.8109 | -1.52702 | 2.59E-05 | 0.001991 |
| FREM1 | 512.0902 | -1.53228 | 0.000213 | 0.009802 |
| NR4A1 | 12509.1 | -1.53326 | 4.72E-06 | 0.000524 |
| ZNF165 | 165.1319 | -1.54053 | 0.001594 | 0.04404 |
| TIPARP | 762.6504 | -1.54629 | 2.14E-07 | 3.92E-05 |
| EGFEM1P | 735.0891 | -1.54686 | 0.000312 | 0.013248 |
| TUBA4A | 1245.658 | -1.54714 | 0.000644 | 0.023088 |
| ANK3 | 156.0673 | -1.54847 | 9.10E-05 | 0.005067 |
| NBPF20 | 139.5187 | -1.5486 | 0.000131 | 0.006678 |
| CPED1 | 452.5243 | -1.55068 | 0.001189 | 0.03613 |
| ZNF543 | 112.2468 | -1.55771 | 0.001914 | 0.049805 |
| MAFF | 4585.117 | -1.56011 | 0.000119 | 0.006238 |

**Supplemental Table 5.** Genes upregulated in ssBM cells from Population *a*

| Gene | baseMean | log2FoldChange | pvalue | padj |
|---|---|---|---|---|
| RHCE | 206.3335 | -1.56223 | 0.00108 | 0.033587 |
| MECOM | 788.3901 | -1.56426 | 3.45E-05 | 0.002443 |
| PTK2 | 350.579 | -1.56815 | 3.40E-05 | 0.002433 |
| ADAM8 | 657.2073 | -1.5695 | 0.000185 | 0.008593 |
| RAPGEF2 | 1329.885 | -1.57592 | 2.79E-09 | 8.37E-07 |
| MANSC1 | 213.3018 | -1.58134 | 1.76E-06 | 0.00023 |
| TSC22D1 | 12922.59 | -1.58237 | 2.72E-05 | 0.002059 |
| LOC107986148 | 56.13932 | -1.58444 | 0.000729 | 0.025132 |
| HIST2H2AA3 | 7143.567 | -1.58444 | 9.98E-08 | 2.01E-05 |
| HIST2H2AA4 | 7143.567 | -1.58444 | 9.98E-08 | 2.01E-05 |
| CFAP70 | 66.19763 | -1.59034 | 0.001295 | 0.038315 |
| CD40 | 311.2899 | -1.59142 | 0.000289 | 0.01252 |
| MLLT3 | 2620.545 | -1.59487 | 3.10E-10 | 1.21E-07 |
| GATA3 | 322.6659 | -1.5982 | 0.000555 | 0.020437 |
| SOCS2 | 3449.74 | -1.60213 | 1.11E-05 | 0.000996 |
| PTPRM | 142.5247 | -1.60245 | 0.00089 | 0.029169 |
| FKBP9 | 273.129 | -1.61014 | 2.10E-06 | 0.000269 |
| HIST1H2BD | 173.8863 | -1.61972 | 6.30E-05 | 0.003793 |
| DOK2 | 154.1238 | -1.62293 | 0.000293 | 0.012606 |
| EPHX2 | 627.2246 | -1.62433 | 0.000365 | 0.014808 |
| FGD4 | 157.1963 | -1.62784 | 5.06E-05 | 0.003232 |
| ALDH1A1 | 3706.4 | -1.63445 | 1.91E-07 | 3.60E-05 |
| MIPOL1 | 274.4249 | -1.65211 | 0.000278 | 0.012123 |
| HIST2H2BE | 2447.134 | -1.65865 | 1.21E-06 | 0.000171 |
| HABP4 | 103.4618 | -1.65956 | 0.001188 | 0.03613 |
| LOC101927745 | 1607.696 | -1.66772 | 1.38E-06 | 0.000188 |
| ZNF467 | 170.3667 | -1.67342 | 0.00039 | 0.015572 |
| STAT4 | 531.802 | -1.6926 | 4.58E-07 | 7.56E-05 |
| UPP1 | 167.7785 | -1.69562 | 0.000349 | 0.014408 |
| LOC107986589 | 240.1177 | -1.70061 | 6.62E-06 | 0.000667 |
| CD83 | 2436.032 | -1.70395 | 1.25E-06 | 0.000175 |
| HIST1H2AC | 483.0348 | -1.70808 | 0.000681 | 0.023935 |
| ZNF204P | 111.9789 | -1.71669 | 0.000127 | 0.006498 |
| EGR3 | 547.4314 | -1.73013 | 3.30E-05 | 0.002397 |
| GRASP | 1936.824 | -1.74119 | 5.42E-10 | 1.91E-07 |
| LOC440895 | 203.5476 | -1.74289 | 0.001194 | 0.036205 |
| GFOD1 | 352.3678 | -1.74405 | 9.67E-07 | 0.000141 |
| DDN-AS1 | 98.83744 | -1.7557 | 4.92E-05 | 0.003154 |
| FOXO1 | 1665.05 | -1.75712 | 2.22E-10 | 9.02E-08 |
| KLF2 | 2734.372 | -1.78378 | 1.33E-07 | 2.66E-05 |
| PCDH9 | 5367.67 | -1.78867 | 1.31E-10 | 5.44E-08 |
| PTGS1 | 2694.26 | -1.79722 | 4.81E-06 | 0.000527 |
| SYTL4 | 457.0231 | -1.80032 | 1.79E-07 | 3.46E-05 |
| MAGI2 | 143.674 | -1.80598 | 6.87E-06 | 0.000681 |
| CCDC42 | 599.0117 | -1.81036 | 8.43E-13 | 5.15E-10 |
| OXT | 65.59233 | -1.81548 | 0.000632 | 0.02284 |
| PDZRN4 | 207.3959 | -1.81784 | 0.001733 | 0.046815 |
| RASSF9 | 103.0175 | -1.81849 | 0.001366 | 0.039716 |
| SETBP1 | 341.3717 | -1.83625 | 0.000934 | 0.030133 |
| TMEM92 | 99.99546 | -1.83662 | 5.58E-05 | 0.003441 |
| CEACAM1 | 66.45265 | -1.84261 | 0.000925 | 0.029973 |
| ACE | 117.8297 | -1.84575 | 0.000101 | 0.005484 |
| GPAT3 | 434.3041 | -1.84632 | 2.97E-08 | 6.87E-06 |
| CLU | 5177.858 | -1.85443 | 0.000316 | 0.013323 |

**Supplemental Table 5.** Genes upregulated in ssBM cells from Population *a*

| Gene | baseMean | log2FoldChange | pvalue | padj |
|---|---|---|---|---|
| PTGER4 | 1821.675 | -1.8581 | 1.38E-06 | 0.000188 |
| GASAL1 | 179.3755 | -1.86324 | 6.43E-05 | 0.003835 |
| ZNF331 | 1961.39 | -1.86736 | 7.44E-08 | 1.55E-05 |
| SERTAD1 | 644.691 | -1.86936 | 5.68E-06 | 0.000591 |
| HOXB7 | 37.40394 | -1.87796 | 0.001793 | 0.047627 |
| PGM5 | 270.6725 | -1.87934 | 9.97E-07 | 0.000144 |
| MLF1 | 467.2907 | -1.88005 | 0.000247 | 0.010967 |
| CXCL8 | 3895.985 | -1.8929 | 1.04E-05 | 0.00096 |
| IL15 | 75.88096 | -1.89851 | 0.001211 | 0.036555 |
| ANXA3 | 177.5677 | -1.90313 | 0.000131 | 0.006678 |
| CDC14B | 146.0723 | -1.90406 | 5.88E-05 | 0.003591 |
| HIST1H2BG | 81.47813 | -1.90407 | 0.000789 | 0.026799 |
| CRHBP | 7058.931 | -1.91836 | 2.02E-06 | 0.000262 |
| THRB | 265.6067 | -1.92016 | 1.71E-06 | 0.000228 |
| LIMCH1 | 583.8056 | -1.94048 | 8.60E-12 | 4.77E-09 |
| LOC101927879 | 1159.694 | -1.94271 | 5.88E-06 | 0.000605 |
| PREX2 | 722.9604 | -1.94731 | 0.000164 | 0.007987 |
| LOC101928489 | 136.304 | -1.95643 | 9.37E-07 | 0.000138 |
| LOC100507103 | 154.9398 | -1.95711 | 2.46E-05 | 0.001927 |
| MAP1LC3A | 79.73896 | -1.96423 | 2.77E-05 | 0.002091 |
| CATIP | 53.09469 | -1.96843 | 0.000932 | 0.030109 |
| MIR22HG | 355.6716 | -1.97097 | 1.32E-06 | 0.000181 |
| LILRB2 | 92.96849 | -1.97224 | 0.000677 | 0.023838 |
| CLEC14A | 157.5006 | -1.97225 | 0.00169 | 0.046105 |
| SH3BP5 | 157.6521 | -1.97305 | 6.22E-05 | 0.003761 |
| NCF1 | 85.16844 | -1.97576 | 0.000345 | 0.014252 |
| MIAT | 105.86 | -1.982 | 3.34E-05 | 0.002415 |
| KLF5 | 360.3383 | -1.98775 | 0.000126 | 0.006479 |
| RRAS | 393.0431 | -1.99973 | 4.08E-10 | 1.56E-07 |
| ETV3 | 2362.906 | -2.01253 | 2.41E-05 | 0.001907 |
| ELL2 | 512.3782 | -2.01731 | 4.05E-11 | 1.88E-08 |
| NPM2 | 137.9675 | -2.02893 | 1.08E-05 | 0.000988 |
| EFNB2 | 104.6937 | -2.02959 | 0.000491 | 0.018632 |
| PTGS2 | 1736.408 | -2.03723 | 1.42E-08 | 3.52E-06 |
| UST | 113.107 | -2.04832 | 8.57E-06 | 0.000822 |
| HIST1H2BC | 317.665 | -2.0495 | 3.97E-06 | 0.000469 |
| PLEC | 275.6216 | -2.05183 | 5.60E-07 | 8.99E-05 |
| LOC105369378 | 74.75662 | -2.05337 | 1.15E-05 | 0.001021 |
| SRPX2 | 87.81358 | -2.05617 | 0.000229 | 0.010376 |
| NR4A2 | 10887.87 | -2.07743 | 1.62E-13 | 1.19E-10 |
| HIST1H2AE | 235.653 | -2.08518 | 2.41E-11 | 1.23E-08 |
| ID2-AS1 | 68.88717 | -2.10008 | 0.000325 | 0.013539 |
| FUT6 | 59.21774 | -2.10406 | 0.000124 | 0.006393 |
| NDRG2 | 357.4437 | -2.1116 | 2.44E-05 | 0.001921 |
| DNM3 | 503.1717 | -2.11164 | 0.000355 | 0.014566 |
| LOC105376637 | 137.8765 | -2.11626 | 0.000538 | 0.019937 |
| MFAP2 | 587.9104 | -2.11733 | 1.11E-09 | 3.56E-07 |
| LOC105374381 | 41.44771 | -2.15098 | 0.000534 | 0.019854 |
| PDZK1IP1 | 121.4362 | -2.15742 | 3.39E-06 | 0.000408 |
| LOC101928202 | 181.097 | -2.1625 | 5.25E-06 | 0.000562 |
| CCL3L1 | 129.4678 | -2.1696 | 1.24E-05 | 0.001079 |
| HSD17B14 | 112.0564 | -2.17942 | 0.000268 | 0.011715 |
| MFAP3L | 42.67948 | -2.18798 | 0.000798 | 0.027078 |
| BHLHE41 | 106.5576 | -2.21272 | 2.86E-05 | 0.002133 |

**Supplemental Table 5.** Genes upregulated in ssBM cells from Population *a*

| Gene | baseMean | log2FoldChange | pvalue | padj |
|---|---|---|---|---|
| TNNC2 | 61.62121 | -2.21818 | 0.000158 | 0.007781 |
| SLC1A6 | 493.3005 | -2.23254 | 9.52E-05 | 0.005253 |
| NLRP12 | 37.07093 | -2.23466 | 0.000811 | 0.027297 |
| FAM241B | 80.30219 | -2.24315 | 4.53E-06 | 0.000512 |
| LOC399900 | 130.7603 | -2.25862 | 0.000165 | 0.008019 |
| AVP | 3349.397 | -2.2913 | 3.02E-13 | 1.98E-10 |
| CNMD | 100.0894 | -2.29575 | 4.00E-05 | 0.002683 |
| LMNA | 5448.089 | -2.326 | 6.15E-06 | 0.000626 |
| SUCLG2 | 371.5444 | -2.33196 | 7.90E-07 | 0.00012 |
| LOC107986939 | 799.3344 | -2.33681 | 5.29E-11 | 2.36E-08 |
| CCL4L1 | 173.7886 | -2.34322 | 4.27E-05 | 0.002811 |
| LINC00891 | 73.00568 | -2.35206 | 1.23E-05 | 0.001079 |
| LOC100132741 | 73.00568 | -2.35206 | 1.23E-05 | 0.001079 |
| AFDN | 393.4498 | -2.35801 | 7.12E-09 | 1.92E-06 |
| JAML | 1729.374 | -2.37699 | 1.78E-07 | 3.46E-05 |
| EMCN | 676.6464 | -2.37941 | 4.33E-14 | 3.96E-11 |
| PHLDB2 | 666.9446 | -2.37967 | 5.04E-14 | 4.39E-11 |
| BRE-AS1 | 773.3593 | -2.39457 | 2.72E-17 | 7.12E-14 |
| LOC107984362 | 42.65862 | -2.42234 | 0.001134 | 0.034783 |
| LZTS3 | 213.9512 | -2.43252 | 7.30E-05 | 0.004247 |
| NR3C2 | 46.28584 | -2.45673 | 0.000175 | 0.008289 |
| EPPK1 | 105.7249 | -2.46024 | 0.000237 | 0.010661 |
| DEPP1 | 97.244 | -2.46399 | 0.000173 | 0.008231 |
| LINC02160 | 67.4482 | -2.46713 | 4.60E-05 | 0.00301 |
| STARD13 | 47.61821 | -2.46981 | 0.000389 | 0.015572 |
| DMKN | 83.81865 | -2.47152 | 1.07E-05 | 0.000981 |
| MPZL2 | 207.2009 | -2.47439 | 1.22E-08 | 3.14E-06 |
| CD4 | 510.5293 | -2.49066 | 8.01E-09 | 2.13E-06 |
| NOS1AP | 28.21995 | -2.50699 | 0.001066 | 0.033221 |
| MYL9 | 93.53567 | -2.50727 | 1.27E-08 | 3.18E-06 |
| PHLDA2 | 48.88345 | -2.52375 | 8.64E-05 | 0.004869 |
| SNED1 | 128.4598 | -2.53757 | 0.001551 | 0.043099 |
| HLF | 1171.426 | -2.54031 | 3.21E-11 | 1.55E-08 |
| SETDB2-PHF11 | 147.5157 | -2.54049 | 0.001902 | 0.049628 |
| ANKRD20A2 | 47.32234 | -2.60145 | 0.00191 | 0.049769 |
| FGF9 | 66.56326 | -2.64487 | 0.000159 | 0.007793 |
| HIST2H3PS2 | 58.99151 | -2.66489 | 7.07E-06 | 0.000696 |
| SPP1 | 920.3645 | -2.67028 | 4.53E-07 | 7.56E-05 |
| FAM198B | 168.1024 | -2.67526 | 1.10E-05 | 0.000992 |
| CDH7 | 296.8143 | -2.68024 | 7.58E-11 | 3.23E-08 |
| WWTR1 | 43.59968 | -2.68525 | 0.000458 | 0.017763 |
| PCDH8 | 29.63688 | -2.7062 | 0.001503 | 0.042217 |
| LOC105378604 | 90.12459 | -2.72061 | 7.27E-05 | 0.00424 |
| BMP6 | 75.40507 | -2.73818 | 2.50E-05 | 0.001952 |
| DIRAS3 | 80.74026 | -2.78779 | 0.001446 | 0.041448 |
| DLK1 | 383.0275 | -2.79465 | 2.61E-16 | 5.96E-13 |
| MCF2L | 30.34498 | -2.79646 | 0.000405 | 0.016025 |
| THRB-AS1 | 25.54178 | -2.80434 | 0.000471 | 0.018068 |
| MDGA2 | 74.50015 | -2.82636 | 7.58E-05 | 0.004381 |
| PCDH17 | 90.6553 | -2.89544 | 4.75E-06 | 0.000524 |
| LOC105374314 | 49.22219 | -2.908 | 3.11E-06 | 0.000382 |
| TCEAL2 | 304.8223 | -2.90852 | 1.85E-13 | 1.30E-10 |
| ADAMTS1 | 132.9171 | -2.91052 | 1.76E-06 | 0.00023 |
| MEG3 | 553.3765 | -2.92508 | 4.18E-10 | 1.56E-07 |

**Supplemental Table 5.** Genes upregulated in ssBM cells from Population *a*

| Gene | baseMean | log2FoldChange | pvalue | padj |
|---|---|---|---|---|
| LOC105374296 | 54.13247 | -2.92808 | 4.73E-05 | 0.003049 |
| MTMR11 | 41.67739 | -2.94433 | 0.000707 | 0.024707 |
| LRP1 | 50.43354 | -2.95119 | 0.000133 | 0.00673 |
| FXYD6 | 383.6558 | -2.96475 | 2.17E-08 | 5.16E-06 |
| FILIP1L | 42.97703 | -3.01296 | 2.79E-05 | 0.002091 |
| RTL5 | 62.38507 | -3.01465 | 6.79E-06 | 0.000679 |
| CXCL1 | 30.72107 | -3.03408 | 0.000318 | 0.013333 |
| ABI3BP | 173.5027 | -3.05433 | 1.14E-11 | 5.99E-09 |
| HES1 | 188.8038 | -3.06801 | 4.20E-06 | 0.00049 |
| HAR1B | 20.57462 | -3.07089 | 0.00159 | 0.043995 |
| RASSF6 | 243.2049 | -3.14791 | 6.16E-11 | 2.69E-08 |
| SELENOM | 273.0694 | -3.18995 | 8.29E-09 | 2.17E-06 |
| LOC101929538 | 43.79003 | -3.23696 | 0.000113 | 0.005977 |
| COL6A2 | 74.10501 | -3.31354 | 2.63E-05 | 0.002006 |
| FAM198B-AS1 | 45.56566 | -3.32526 | 3.21E-05 | 0.002351 |
| ARHGAP20 | 29.16205 | -3.35183 | 0.001762 | 0.047248 |
| HAS2 | 50.99963 | -3.40934 | 8.04E-06 | 0.000779 |
| CCDC184 | 91.52447 | -3.42179 | 2.56E-08 | 6.01E-06 |
| ADCY4 | 27.34296 | -3.44098 | 4.25E-05 | 0.002811 |
| LOC107987347 | 26.46104 | -3.45997 | 3.90E-05 | 0.002656 |
| CLEC3B | 598.3781 | -3.47507 | 3.29E-16 | 6.02E-13 |
| SCG5 | 45.83243 | -3.48202 | 3.15E-06 | 0.000385 |
| LOC105374657 | 26.98195 | -3.62815 | 0.001005 | 0.031675 |
| KCTD7 | 27.93318 | -3.63992 | 0.000482 | 0.018345 |
| RHOD | 24.17149 | -3.71534 | 0.000327 | 0.013565 |
| LDHD | 45.32148 | -3.74473 | 0.000375 | 0.015127 |
| LIF | 29.88115 | -3.75009 | 0.00165 | 0.045527 |
| LRRC4C | 30.7751 | -3.83589 | 0.000117 | 0.006152 |
| CUEDC1 | 23.06678 | -3.95 | 3.79E-05 | 0.002603 |
| MEIS1-AS3 | 36.58593 | -3.96697 | 4.67E-05 | 0.003034 |
| AMOTL2 | 31.705 | -4.10494 | 8.39E-05 | 0.004802 |
| DOK5 | 18.54238 | -4.12628 | 0.001314 | 0.038691 |
| SOGA3 | 28.14087 | -4.18379 | 0.000735 | 0.025272 |
| TCEA3 | 19.68283 | -4.19593 | 0.00046 | 0.017822 |
| TNS2 | 14.85093 | -4.32697 | 0.000951 | 0.030406 |
| SPAG17 | 21.09394 | -4.39854 | 0.001333 | 0.039135 |
| AKAP3 | 13.90518 | -4.58011 | 0.0014 | 0.040501 |
| DBNDD1 | 24.39196 | -4.62588 | 0.000971 | 0.030826 |
| KCNJ8 | 24.83811 | -4.8033 | 3.43E-05 | 0.002433 |
| LOC105379499 | 83.02463 | -5.28672 | 2.15E-06 | 0.000274 |
| CCL2 | 19.35472 | -5.34504 | 0.000711 | 0.024749 |
| LOC107987433 | 12.75724 | -5.81218 | 0.00136 | 0.039716 |
| SLCO1C1 | 18.0267 | -6.68772 | 0.000222 | 0.010056 |
| LOC105371406 | 14.60163 | -7.10165 | 0.000103 | 0.005558 |
| CFB | 12.54505 | -7.93245 | 0.000244 | 0.010857 |
| WIF1 | 19.67862 | -8.14895 | 3.63E-05 | 0.002543 |
| NUDT4P2 | 28.82727 | -8.28338 | 5.92E-07 | 9.27E-05 |
| C15orf38-AP3S2 | 133.3133 | -10.2109 | 3.64E-05 | 0.002543 |
| PLGLB1 | 36.95608 | -24.4727 | 1.24E-08 | 3.15E-06 |

**Supplemental Table 6**. Genes upregulated in ssBM cells from Population *b*

| Gene | baseMean | log2FoldChange | pvalue | padj |
|---|---|---|---|---|
| MS4A3 | 607.2288 | 11.64698 | 2.72E-14 | 2.62E-11 |
| ELANE | 3338.447 | 11.01888 | 8.09E-14 | 6.44E-11 |
| PRTN3 | 1793.53 | 10.71206 | 5.29E-14 | 4.41E-11 |
| AZU1 | 1340.101 | 10.49691 | 2.46E-07 | 4.42E-05 |
| KCNE5 | 468.52 | 10.14964 | 7.57E-12 | 4.33E-09 |
| PLPPR3 | 75.26939 | 9.549211 | 2.27E-05 | 0.001818 |
| RNASE3 | 381.9253 | 9.273937 | 3.48E-06 | 0.000414 |
| CLEC12A | 239.7496 | 8.876084 | 1.62E-15 | 2.28E-12 |
| MPO | 24481.47 | 8.572393 | 1.11E-15 | 1.69E-12 |
| IRF8 | 1581.016 | 8.542745 | 1.09E-14 | 1.25E-11 |
| CLEC5A | 14.92012 | 8.204835 | 8.70E-05 | 0.004886 |
| ECRP | 207.42 | 7.998945 | 5.13E-07 | 8.39E-05 |
| TERT | 19.56845 | 7.747428 | 6.41E-05 | 0.003835 |
| RNASE2 | 2188.5 | 7.606182 | 1.68E-31 | 1.54E-27 |
| KCNH2 | 930.6124 | 7.58153 | 6.20E-05 | 0.00376 |
| LPO | 22.13128 | 7.430327 | 9.30E-05 | 0.005144 |
| TRIB2 | 382.4477 | 6.992845 | 5.57E-06 | 0.000583 |
| CEBPE | 12.07256 | 6.960807 | 0.000499 | 0.018882 |
| NAPSB | 237.2018 | 6.920238 | 5.39E-06 | 0.000571 |
| LEF1 | 96.25195 | 6.693554 | 5.34E-05 | 0.003351 |
| FBLN2 | 81.45378 | 6.540263 | 7.22E-05 | 0.004226 |
| JCHAIN | 660.8638 | 6.476391 | 0.000465 | 0.017942 |
| FAM178B | 334.0733 | 6.45272 | 2.52E-05 | 0.001955 |
| UGT3A2 | 212.0576 | 6.373831 | 1.97E-05 | 0.001595 |
| LOC284600 | 18.40594 | 6.312331 | 0.000119 | 0.006228 |
| MICALCL | 16.6594 | 6.247752 | 0.001569 | 0.043473 |
| CPA3 | 2192.082 | 6.070702 | 1.28E-38 | 2.35E-34 |
| LOC105379461 | 119.7916 | 5.799834 | 0.000422 | 0.016645 |
| RHAG | 171.4568 | 5.772701 | 0.00029 | 0.01252 |
| HCK | 41.62468 | 5.492035 | 2.83E-07 | 5.03E-05 |
| RNASE1 | 91.01385 | 5.383341 | 0.000519 | 0.019435 |
| AFF2 | 25.52514 | 5.361965 | 6.87E-05 | 0.004072 |
| CEBPD | 270.2662 | 5.284004 | 1.92E-14 | 1.96E-11 |
| IGSF6 | 51.60483 | 5.232535 | 0.000939 | 0.030179 |
| LINC01835 | 37.86629 | 5.030152 | 0.001726 | 0.046779 |
| LOC554249 | 33.45514 | 4.979561 | 9.98E-05 | 0.005457 |
| TPSAB1 | 1127.254 | 4.956247 | 7.41E-18 | 2.26E-14 |
| CPB1 | 34.83877 | 4.744842 | 0.001804 | 0.047627 |
| SUCNR1 | 163.0882 | 4.389531 | 1.41E-09 | 4.46E-07 |
| TARP | 218.4038 | 4.370905 | 4.50E-21 | 1.65E-17 |
| CTSG | 1023.798 | 4.291867 | 1.73E-14 | 1.87E-11 |
| IGFBP2 | 1092.244 | 4.280073 | 6.41E-24 | 2.94E-20 |
| DNTT | 6867.661 | 4.242558 | 3.41E-05 | 0.002433 |
| ANK1 | 325.0213 | 4.129885 | 5.18E-10 | 1.86E-07 |
| TREM1 | 98.91869 | 4.124762 | 2.03E-07 | 3.76E-05 |
| LINC01971 | 32.86015 | 4.085641 | 7.22E-05 | 0.004226 |
| HBD | 4271.737 | 4.072499 | 5.42E-15 | 6.62E-12 |
| NETO2 | 23.47577 | 3.991747 | 0.000732 | 0.025193 |
| IGLL1 | 11801.95 | 3.981441 | 1.95E-28 | 1.19E-24 |
| PROK2 | 18.63179 | 3.891693 | 0.000656 | 0.023324 |
| FCGR1A | 49.23802 | 3.873212 | 0.000849 | 0.028223 |
| HGF | 157.2729 | 3.867161 | 6.49E-06 | 0.000657 |
| MIR181A1HG | 258.7451 | 3.854662 | 1.05E-15 | 1.69E-12 |
| RAB44 | 71.83456 | 3.825197 | 1.05E-09 | 3.44E-07 |

**Supplemental Table 6**. Genes upregulated in ssBM cells from Population *b*

| Gene | baseMean | log2FoldChange | pvalue | padj |
|---|---|---|---|---|
| SPON2 | 54.8989 | 3.644636 | 0.000222 | 0.010056 |
| TRPC6 | 35.4843 | 3.513891 | 0.00087 | 0.028654 |
| KCNG2 | 189.2824 | 3.482407 | 9.51E-07 | 0.000139 |
| LOC105379749 | 23.81137 | 3.427595 | 0.00058 | 0.021245 |
| ACPP | 15.90212 | 3.423181 | 0.001778 | 0.047472 |
| CST7 | 731.4084 | 3.422599 | 1.33E-12 | 7.89E-10 |
| PDE1A | 36.73978 | 3.410226 | 0.000101 | 0.005484 |
| LGALS1 | 1583.29 | 3.368345 | 4.68E-15 | 6.12E-12 |
| HPGDS | 636.485 | 3.318455 | 5.17E-13 | 3.26E-10 |
| CD40LG | 162.5394 | 3.312942 | 0.000652 | 0.023289 |
| DHRS3 | 187.7447 | 3.308511 | 1.49E-08 | 3.65E-06 |
| TSPOAP1 | 207.4253 | 3.271188 | 1.12E-11 | 5.99E-09 |
| MARS2 | 52.86662 | 3.251556 | 1.30E-06 | 0.00018 |
| CSF1R | 180.661 | 3.232254 | 5.40E-05 | 0.003372 |
| RFX8 | 26.00452 | 3.216624 | 0.000474 | 0.018103 |
| CNRIP1 | 1305.972 | 3.213938 | 5.92E-07 | 9.27E-05 |
| CFD | 1202.782 | 3.145938 | 2.17E-13 | 1.47E-10 |
| CD96 | 142.9638 | 3.135199 | 3.32E-08 | 7.42E-06 |
| LOC339862 | 100.929 | 3.09102 | 3.23E-05 | 0.002358 |
| VPREB1 | 534.7383 | 2.994454 | 2.91E-10 | 1.16E-07 |
| FUT7 | 209.1783 | 2.991896 | 3.28E-08 | 7.42E-06 |
| HOMER3 | 47.72576 | 2.991725 | 8.58E-05 | 0.004869 |
| NKG7 | 395.9929 | 2.938181 | 6.53E-10 | 2.26E-07 |
| CSF1 | 1494.019 | 2.930322 | 1.88E-07 | 3.58E-05 |
| CD1D | 20.58097 | 2.921299 | 0.000847 | 0.028195 |
| KLF1 | 1091.762 | 2.884433 | 4.50E-06 | 0.000512 |
| ACSM1 | 270.7297 | 2.881232 | 1.96E-05 | 0.001593 |
| STXBP6 | 162.5321 | 2.740636 | 2.38E-06 | 0.000299 |
| LOC101927497 | 930.7049 | 2.709917 | 1.03E-09 | 3.43E-07 |
| S100B | 42.74109 | 2.696185 | 0.000424 | 0.016686 |
| TESC | 654.2503 | 2.616938 | 2.93E-16 | 5.96E-13 |
| TUBB6 | 355.8487 | 2.593785 | 3.40E-07 | 5.93E-05 |
| CEBPA | 471.8078 | 2.559527 | 9.12E-14 | 6.96E-11 |
| STAR | 95.7509 | 2.552679 | 1.79E-05 | 0.001477 |
| LOC100130872 | 115.8596 | 2.493392 | 1.54E-06 | 0.000208 |
| IL1RAP | 97.00157 | 2.4871 | 3.31E-09 | 9.61E-07 |
| C16orf74 | 192.4399 | 2.482242 | 3.84E-05 | 0.002624 |
| GPT2 | 245.5668 | 2.468931 | 4.53E-10 | 1.66E-07 |
| VSTM1 | 103.3851 | 2.447183 | 8.92E-06 | 0.000848 |
| ZNF442 | 30.68266 | 2.421433 | 0.00051 | 0.019206 |
| AKAP2 | 280.447 | 2.40638 | 4.55E-07 | 7.56E-05 |
| PAG1 | 215.9837 | 2.387848 | 3.66E-07 | 6.26E-05 |
| MYC | 1854.732 | 2.380399 | 3.13E-11 | 1.55E-08 |
| LTBP1 | 390.4288 | 2.376862 | 2.79E-06 | 0.000347 |
| ASCL2 | 48.80689 | 2.376829 | 0.000121 | 0.006287 |
| DRICH1 | 43.11086 | 2.339524 | 0.00024 | 0.010711 |
| RAB7B | 79.12822 | 2.271861 | 4.52E-06 | 0.000512 |
| SIRPB2 | 107.3983 | 2.259526 | 0.000869 | 0.028654 |
| CRYM | 193.6112 | 2.243122 | 0.000152 | 0.007588 |
| MRC2 | 45.62938 | 2.238982 | 0.00146 | 0.041589 |
| P2RX5 | 221.2233 | 2.234379 | 0.001555 | 0.043159 |
| NPW | 434.6643 | 2.217578 | 5.51E-07 | 8.94E-05 |
| LRRC26 | 140.6155 | 2.160469 | 6.36E-05 | 0.003819 |
| LOC105372233 | 73.30996 | 2.151351 | 9.36E-06 | 0.000876 |

**Supplemental Table 6**. Genes upregulated in ssBM cells from Population *b*

| Gene | baseMean | log2FoldChange | pvalue | padj |
|---|---|---|---|---|
| UCA1 | 260.3523 | 2.145708 | 0.000385 | 0.01549 |
| SLCO5A1 | 102.3807 | 2.139775 | 5.82E-05 | 0.003564 |
| KBTBD11 | 265.486 | 2.122802 | 1.26E-05 | 0.001088 |
| ZNF695 | 85.25526 | 2.105754 | 0.000107 | 0.005722 |
| MINDY4 | 82.25269 | 2.063057 | 0.000461 | 0.017823 |
| TRH | 194.6018 | 2.060779 | 7.69E-08 | 1.58E-05 |
| BEND6 | 81.85684 | 2.039589 | 5.21E-06 | 0.000561 |
| LMNB1 | 526.2213 | 2.022465 | 2.60E-05 | 0.001992 |
| SERPINB8 | 651.7298 | 1.943168 | 0.001311 | 0.038673 |
| TRPM2 | 73.18995 | 1.933043 | 0.0004 | 0.015857 |
| ANXA2 | 1723.198 | 1.898142 | 6.56E-07 | 0.000102 |
| RHEX | 759.3652 | 1.897984 | 0.000257 | 0.011365 |
| PDSS1 | 238.9514 | 1.892147 | 1.45E-05 | 0.00122 |
| PDK1 | 283.3398 | 1.887257 | 9.64E-05 | 0.005304 |
| MZB1 | 1896.12 | 1.885126 | 6.85E-10 | 2.33E-07 |
| IL17RA | 520.6521 | 1.879891 | 1.92E-08 | 4.62E-06 |
| SPNS3 | 2472.1 | 1.85919 | 2.28E-09 | 6.96E-07 |
| CDCA7 | 1924.989 | 1.859051 | 1.36E-05 | 0.001159 |
| LOC101928834 | 1097.315 | 1.853409 | 1.37E-07 | 2.70E-05 |
| RTN4R | 103.8926 | 1.851772 | 0.00055 | 0.020297 |
| LOC107984120 | 84.38488 | 1.818159 | 0.001693 | 0.046105 |
| SLC45A3 | 151.8236 | 1.815311 | 5.12E-05 | 0.003257 |
| TTK | 172.8823 | 1.805955 | 0.000159 | 0.007793 |
| FAM107B | 371.3967 | 1.804149 | 3.94E-05 | 0.002668 |
| TTC7A | 602.9927 | 1.790046 | 2.38E-06 | 0.000299 |
| MT1X | 198.1518 | 1.752106 | 1.17E-05 | 0.001033 |
| LOC105370401 | 178.5539 | 1.728139 | 4.33E-05 | 0.002842 |
| FAM129A | 403.8657 | 1.725051 | 2.02E-07 | 3.76E-05 |
| LDLRAD3 | 131.2002 | 1.721564 | 0.000747 | 0.025547 |
| SPARC | 3306.608 | 1.71001 | 0.001679 | 0.046105 |
| P2RY2 | 140.5264 | 1.695108 | 0.000838 | 0.028069 |
| SDK2 | 312.6588 | 1.688676 | 0.000926 | 0.029973 |
| LOC105372857 | 77.66211 | 1.677036 | 8.62E-05 | 0.004869 |
| SLC16A10 | 159.2667 | 1.654852 | 0.000174 | 0.008272 |
| C1QTNF4 | 2381.782 | 1.632465 | 1.33E-05 | 0.001139 |
| ARHGAP10 | 120.3269 | 1.628189 | 0.000471 | 0.018068 |
| MYCN | 239.5675 | 1.627962 | 0.000183 | 0.008542 |
| PRAM1 | 765.5684 | 1.627 | 0.000237 | 0.010661 |
| CD38 | 1386.06 | 1.625371 | 6.75E-09 | 1.85E-06 |
| PLIN2 | 2695.201 | 1.610381 | 2.54E-05 | 0.00196 |
| ABHD17C | 267.8162 | 1.572601 | 4.62E-06 | 0.00052 |
| FYB1 | 367.0019 | 1.570121 | 0.000316 | 0.013323 |
| TST | 495.7916 | 1.56798 | 3.11E-05 | 0.002285 |
| MT2A | 729.4244 | 1.553333 | 0.000828 | 0.027843 |
| IGFBP7 | 5311.886 | 1.552907 | 5.93E-08 | 1.26E-05 |
| CDC7 | 244.6241 | 1.542489 | 0.000842 | 0.028091 |
| ANLN | 153.2057 | 1.541104 | 5.15E-05 | 0.003266 |
| PPIF | 1495.653 | 1.523318 | 0.000386 | 0.015512 |
| FABP5 | 4235.355 | 1.522576 | 3.22E-07 | 5.67E-05 |
| NT5DC2 | 771.2156 | 1.515173 | 0.000221 | 0.010056 |
| CITED4 | 344.7245 | 1.503432 | 1.93E-05 | 0.001581 |
| ADGRG5 | 134.485 | 1.495678 | 0.000899 | 0.029319 |
| ZNF385C | 101.6618 | 1.489025 | 0.000297 | 0.01273 |
| TNFSF13B | 10619.57 | 1.458911 | 5.68E-05 | 0.00349 |

**Supplemental Table 6**. Genes upregulated in ssBM cells from Population *b*

| Gene | baseMean | log2FoldChange | pvalue | padj |
|---|---|---|---|---|
| EEF1AKMT4 | 397.9827 | 1.441077 | 0.000989 | 0.031287 |
| LOC105373444 | 1779.056 | 1.433777 | 1.01E-06 | 0.000144 |
| FARSA | 2129.909 | 1.429512 | 0.001255 | 0.037597 |
| TSPOAP1-AS1 | 194.0967 | 1.426105 | 7.99E-06 | 0.000778 |
| CD48 | 233.0326 | 1.423637 | 5.19E-05 | 0.003281 |
| ZNF724 | 147.7298 | 1.419989 | 0.001526 | 0.042611 |
| EFCAB2 | 480.3331 | 1.413263 | 6.08E-08 | 1.28E-05 |
| GRPEL1 | 700.9984 | 1.404142 | 8.55E-07 | 0.000128 |
| DUSP10 | 1095.337 | 1.40237 | 5.80E-07 | 9.24E-05 |
| BAHCC1 | 187.5737 | 1.39776 | 0.000112 | 0.005962 |
| POLE2 | 589.5123 | 1.389554 | 0.000114 | 0.00602 |
| ZAP70 | 131.5138 | 1.381731 | 0.00012 | 0.006268 |
| GGA2 | 734.5146 | 1.375794 | 0.000669 | 0.023656 |
| NUP210 | 1449.691 | 1.363709 | 7.82E-07 | 0.000119 |
| ARMH1 | 1408.331 | 1.358346 | 2.12E-09 | 6.60E-07 |
| UHRF1 | 1275.159 | 1.357351 | 0.000168 | 0.008073 |
| SKA3 | 640.9667 | 1.356826 | 0.000624 | 0.022635 |
| LRR1 | 505.6524 | 1.350397 | 3.62E-05 | 0.002543 |
| LOC107984868 | 126.6638 | 1.32373 | 0.001055 | 0.032969 |
| MGST1 | 4696.518 | 1.31443 | 9.33E-06 | 0.000876 |
| CHEK1 | 701.3012 | 1.31129 | 0.000112 | 0.005954 |
| TYROBP | 377.1557 | 1.30777 | 9.37E-06 | 0.000876 |
| CRYBG1 | 207.8736 | 1.307472 | 0.001001 | 0.031617 |
| MAD2L1 | 2102.308 | 1.29758 | 0.000239 | 0.010686 |
| SMIM24 | 4900.651 | 1.295646 | 3.00E-08 | 6.87E-06 |
| BLNK | 1122.722 | 1.286992 | 0.001109 | 0.034345 |
| MRPS23 | 1001.604 | 1.285256 | 0.000317 | 0.013333 |
| ST3GAL6 | 344.7999 | 1.282841 | 4.36E-06 | 0.000506 |
| RAB32 | 2710.813 | 1.279598 | 0.0005 | 0.018882 |
| ARHGEF18 | 256.7524 | 1.275331 | 8.27E-05 | 0.004749 |
| NUP85 | 612.6285 | 1.270616 | 0.001883 | 0.049208 |
| DNAAF3 | 359.7196 | 1.265551 | 0.001803 | 0.047627 |
| MRPS12 | 1457.201 | 1.263786 | 4.25E-05 | 0.002811 |
| SLC35F2 | 463.763 | 1.261346 | 0.000398 | 0.015797 |
| ABHD3 | 344.5468 | 1.259439 | 0.00076 | 0.025925 |
| MCM4 | 2529.492 | 1.221955 | 8.79E-05 | 0.004909 |
| HCST | 1348.958 | 1.218707 | 0.000835 | 0.028025 |
| PLAC8 | 5580.424 | 1.200975 | 0.000139 | 0.007005 |
| LYZ | 2881.135 | 1.190353 | 0.000863 | 0.028592 |
| C8orf76 | 211.4913 | 1.188032 | 0.000286 | 0.012447 |
| GMNN | 1011.428 | 1.186997 | 0.000104 | 0.005609 |
| NRG4 | 533.0171 | 1.182366 | 0.001689 | 0.046105 |
| MRPL58 | 604.0773 | 1.181921 | 0.000589 | 0.021495 |
| METRNL | 391.5022 | 1.176907 | 2.67E-05 | 0.002027 |
| OIP5 | 562.5605 | 1.174102 | 0.000367 | 0.014845 |
| ANKLE1 | 252.0194 | 1.172693 | 0.000472 | 0.018068 |
| MRPL13 | 1551.42 | 1.169444 | 2.36E-05 | 0.001874 |
| RFWD3 | 428.5924 | 1.158209 | 0.001119 | 0.034571 |
| PPAT | 270.247 | 1.156627 | 0.00045 | 0.017492 |
| CHCHD1 | 643.1064 | 1.153341 | 2.04E-05 | 0.001643 |
| GEMIN2 | 371.9574 | 1.1523 | 0.0009 | 0.029319 |
| TFRC | 2773.061 | 1.14686 | 0.00049 | 0.018632 |
| GYPC | 5729.818 | 1.143697 | 5.38E-06 | 0.000571 |
| POLE3 | 3493.003 | 1.142647 | 3.45E-06 | 0.000413 |

**Supplemental Table 6**. Genes upregulated in ssBM cells from Population *b*

| Gene | baseMean | log2FoldChange | pvalue | padj |
|---|---|---|---|---|
| DPCD | 431.1146 | 1.134572 | 0.000636 | 0.022842 |
| AIMP2 | 1275.279 | 1.133522 | 8.61E-05 | 0.004869 |
| MRPL36 | 714.3469 | 1.132929 | 2.95E-06 | 0.000365 |
| CLDN10 | 1618.661 | 1.129737 | 4.06E-05 | 0.002713 |
| XRCC2 | 354.914 | 1.115781 | 0.001345 | 0.039418 |
| BCL2 | 392.3883 | 1.112685 | 0.00017 | 0.008137 |
| NDUFAB1 | 2740.897 | 1.110825 | 7.14E-05 | 0.004207 |
| CKS2 | 3014.415 | 1.107042 | 0.000427 | 0.016756 |
| COLGALT1 | 792.7393 | 1.100381 | 3.95E-05 | 0.002668 |
| CD33 | 465.3276 | 1.094388 | 0.001694 | 0.046105 |
| CDT1 | 1060.923 | 1.093038 | 4.62E-05 | 0.003014 |
| POLR3K | 982.3258 | 1.076698 | 0.000357 | 0.014633 |
| MEF2A | 419.799 | 1.075965 | 6.65E-05 | 0.003953 |
| C15orf61 | 478.9456 | 1.066835 | 0.000321 | 0.013377 |
| TIPIN | 552.2371 | 1.064203 | 0.000319 | 0.013337 |
| CENPW | 1080.57 | 1.060986 | 0.000529 | 0.019731 |
| MLEC | 2816.976 | 1.059908 | 1.14E-05 | 0.001019 |
| USP3 | 532.0931 | 1.059105 | 0.000969 | 0.030826 |
| NDUFB3 | 2106.104 | 1.056119 | 0.000153 | 0.007603 |
| TMIGD2 | 690.6633 | 1.047661 | 0.000361 | 0.014716 |
| OPN3 | 1067.18 | 1.039465 | 3.36E-05 | 0.002425 |
| FOXRED2 | 485.8497 | 1.034912 | 0.00087 | 0.028654 |
| ATOX1 | 886.7135 | 1.034758 | 0.001169 | 0.035703 |
| FAM45A | 1590.409 | 1.030316 | 0.000708 | 0.024707 |
| KPTN | 371.3055 | 1.018284 | 0.00054 | 0.019986 |
| MXD1 | 754.7275 | 1.008042 | 0.00044 | 0.017179 |

**Supplemental Table 7**. Genes upregulated in GCSF-mobilized cells from Population *a*

| Gene | baseMean | log2FoldChange | pvalue | padj |
|---|---|---|---|---|
| EPPK1 | 73.89057 | -3.0174 | 1.35E-09 | 7.15E-07 |
| STAB1 | 116.8942 | -2.33686 | 8.67E-07 | 0.000196 |
| PRG2 | 84.50356 | -2.33287 | 0.000625 | 0.027761 |
| LOC112268342 | 64.50085 | -2.32152 | 0.000119 | 0.008576 |
| TCEAL2 | 81.40736 | -2.28447 | 3.05E-07 | 7.62E-05 |
| OTUD7B | 53.3713 | -2.26405 | 8.34E-05 | 0.006731 |
| LRP1 | 104.4242 | -2.19666 | 4.55E-08 | 1.52E-05 |
| LOC112268284 | 77.94825 | -2.11867 | 1.82E-05 | 0.002148 |
| JAML | 238.4582 | -2.07656 | 4.79E-11 | 4.70E-08 |
| PCDH17 | 58.78369 | -2.07385 | 0.000116 | 0.008498 |
| PHLDB1 | 75.21931 | -2.05308 | 3.05E-06 | 0.000532 |
| OR2H2 | 57.11326 | -2.04934 | 0.001264 | 0.046884 |
| LOC101928202 | 70.52762 | -1.99069 | 0.000389 | 0.020088 |
| PDE1C | 88.3771 | -1.96495 | 1.69E-06 | 0.000336 |
| SUCLG2 | 65.69795 | -1.90505 | 0.000254 | 0.014479 |
| LOC107985787 | 77.63033 | -1.85902 | 1.93E-05 | 0.002177 |
| COL6A1 | 72.57931 | -1.74887 | 0.000101 | 0.007614 |
| AVP | 76.40189 | -1.68317 | 0.000337 | 0.017784 |
| BRSK2 | 92.76892 | -1.66682 | 2.05E-05 | 0.002264 |
| CD4 | 65.97593 | -1.66038 | 0.000654 | 0.02856 |
| PLEKHA6 | 71.01237 | -1.65409 | 0.000335 | 0.017735 |
| NRBP2 | 136.8077 | -1.5805 | 4.43E-06 | 0.000706 |
| EZR-AS1 | 70.3506 | -1.57537 | 0.001079 | 0.042485 |
| MEG3 | 295.2694 | -1.57422 | 1.79E-08 | 7.01E-06 |
| RNF103-CHMP3 | 95.74023 | -1.5557 | 0.000381 | 0.01975 |
| PREX2 | 394.2795 | -1.52343 | 3.71E-09 | 1.70E-06 |
| LZTS3 | 181.5105 | -1.50854 | 2.26E-06 | 0.000419 |
| MIR29B2CHG | 127.4679 | -1.49839 | 3.55E-05 | 0.003426 |
| COL6A2 | 92.8827 | -1.47465 | 0.000489 | 0.023863 |
| HLF | 1162.746 | -1.47031 | 7.64E-16 | 2.62E-12 |
| KLF2 | 963.0647 | -1.43323 | 2.78E-13 | 4.76E-10 |
| SELP | 123.7321 | -1.43018 | 6.26E-05 | 0.005345 |
| ABI3BP | 102.6122 | -1.39117 | 0.000197 | 0.012277 |
| RASSF6 | 169.2947 | -1.37693 | 0.000171 | 0.01109 |
| LOC148696 | 360.1981 | -1.33721 | 2.57E-08 | 9.28E-06 |
| SNED1 | 244.4504 | -1.29861 | 2.19E-06 | 0.000418 |
| ACOT11 | 100.7004 | -1.27487 | 0.001408 | 0.049955 |
| AGAP3 | 294.9761 | -1.25232 | 2.83E-07 | 7.20E-05 |
| LOC105369748 | 134.0892 | -1.22186 | 0.000228 | 0.013294 |
| DOCK3 | 159.4469 | -1.21699 | 0.000157 | 0.01041 |
| LOC105378604 | 143.1471 | -1.2162 | 0.000315 | 0.016918 |
| MIAT | 952.9127 | -1.21189 | 4.95E-08 | 1.62E-05 |
| IL4R | 178.1058 | -1.2041 | 5.12E-05 | 0.004561 |
| RTL5 | 180.4075 | -1.18581 | 5.28E-05 | 0.004675 |
| AFDN | 451.9378 | -1.1683 | 7.26E-08 | 2.21E-05 |
| RAPGEF3 | 151.9928 | -1.16556 | 0.000193 | 0.012173 |
| ITGA2B | 382.0294 | -1.14449 | 0.000574 | 0.026328 |
| SIK1 | 531.9297 | -1.1304 | 0.000159 | 0.010472 |
| PRDM16 | 245.5592 | -1.11417 | 1.31E-05 | 0.001631 |
| ABCA13 | 254.6232 | -1.10487 | 0.001361 | 0.049488 |
| PTK2 | 218.982 | -1.09202 | 6.94E-05 | 0.005879 |
| LOC107984427 | 288.7695 | -1.08005 | 0.000763 | 0.032616 |
| PRR5 | 177.5193 | -1.07903 | 0.001344 | 0.049174 |
| ARHGEF40 | 388.5902 | -1.0238 | 0.000121 | 0.008662 |

**Supplemental Table 7**. Genes upregulated in GCSF-mobilized cells from Population *a*

| Gene | baseMean | log2FoldChange | pvalue | padj |
|---|---|---|---|---|
| PNP | 389.2648 | -1.0173 | 7.03E-06 | 0.001005 |
| PEAR1 | 380.487 | -1.01194 | 2.16E-05 | 0.002334 |
| ULK1 | 371.2976 | -1.0108 | 2.46E-05 | 0.002599 |
| HIST2H3PS2 | 285.7964 | -1.00565 | 7.56E-05 | 0.00621 |

**Supplemental Table 8.** Genes upregulated in GCSF-mobilized cells from Population *b*

| Gene | baseMean | log2FoldChange | p-value | padj |
|---|---|---|---|---|
| LMNB1 | 554.0249 | 1.003174 | 1.67E-06 | 0.000336 |
| CDK6 | 24456.56 | 1.004854 | 2.06E-18 | 1.41E-14 |
| SPARC | 946.2339 | 1.013527 | 2.77E-08 | 9.73E-06 |
| RNF130 | 1738.676 | 1.015676 | 2.31E-07 | 6.23E-05 |
| RPSAP58 | 232.565 | 1.020366 | 0.00064 | 0.028137 |
| MAMDC2 | 886.728 | 1.030157 | 6.00E-10 | 3.43E-07 |
| ZBTB16 | 1324.212 | 1.036293 | 1.41E-10 | 1.08E-07 |
| SNHG19 | 281.3569 | 1.038539 | 2.66E-05 | 0.002747 |
| TFRC | 4387.958 | 1.040528 | 2.54E-09 | 1.29E-06 |
| PTPN14 | 282.7238 | 1.04524 | 8.17E-05 | 0.006634 |
| HPGD | 317.6832 | 1.047418 | 5.24E-06 | 0.000808 |
| EXOSC1 | 195.0903 | 1.050955 | 0.000351 | 0.018332 |
| SELPLG | 177.9986 | 1.057197 | 0.001286 | 0.047264 |
| PLEK | 2010.876 | 1.070104 | 1.85E-10 | 1.34E-07 |
| PIK3R3 | 254.6517 | 1.077499 | 3.94E-05 | 0.003649 |
| PCNA | 339.6969 | 1.080337 | 0.000148 | 0.010011 |
| KBTBD11 | 371.4965 | 1.100214 | 6.47E-06 | 0.000953 |
| FAM129A | 519.471 | 1.11022 | 1.59E-07 | 4.55E-05 |
| DNAJC6 | 315.4325 | 1.117944 | 3.70E-05 | 0.003478 |
| LOC107985855 | 261.9556 | 1.139032 | 3.09E-05 | 0.003028 |
| BLM | 210.8141 | 1.140574 | 0.000316 | 0.016918 |
| ZNF675 | 156.9312 | 1.143483 | 0.000304 | 0.016507 |
| CEBPA | 253.0219 | 1.150087 | 4.91E-05 | 0.004496 |
| TIMELESS | 381.1154 | 1.156667 | 5.02E-05 | 0.004536 |
| DTL | 335.2747 | 1.157131 | 0.00011 | 0.008232 |
| PRIM1 | 178.6158 | 1.160009 | 0.000317 | 0.016942 |
| CD38 | 961.0658 | 1.180657 | 4.81E-12 | 5.08E-09 |
| LOC105372857 | 161.7886 | 1.186601 | 0.000922 | 0.037669 |
| FAM107B | 137.4953 | 1.22731 | 0.001385 | 0.049716 |
| ZCCHC18 | 140.9743 | 1.237721 | 0.000425 | 0.021599 |
| LOC101928834 | 145.1367 | 1.239707 | 0.000828 | 0.034762 |
| CDC6 | 150.9873 | 1.250052 | 0.001084 | 0.042485 |
| KCNQ5 | 412.8885 | 1.255548 | 6.77E-08 | 2.11E-05 |
| RAB44 | 198.3617 | 1.257678 | 0.000219 | 0.013048 |
| POLQ | 184.4625 | 1.291719 | 0.000287 | 0.015882 |
| IGFBP7 | 436.4087 | 1.298038 | 4.92E-09 | 2.18E-06 |
| ASPM | 142.9632 | 1.299962 | 0.000446 | 0.022443 |
| LOC107984120 | 333.8278 | 1.312838 | 2.42E-08 | 8.99E-06 |
| LOC105370401 | 498.3724 | 1.31335 | 8.77E-11 | 7.08E-08 |
| HGF | 170.0861 | 1.313625 | 2.77E-05 | 0.002815 |
| CD69 | 2211.931 | 1.336401 | 7.89E-15 | 1.80E-11 |
| CENPU | 148.0365 | 1.35876 | 0.000191 | 0.012115 |
| PLD1 | 156.9739 | 1.400327 | 0.00019 | 0.012111 |
| NCAPG | 93.9426 | 1.431009 | 0.001264 | 0.046884 |
| CDCA7 | 1006.75 | 1.47136 | 1.89E-12 | 2.16E-09 |
| KCNE3 | 147.3216 | 1.487753 | 5.51E-05 | 0.004847 |
| KCNK5 | 192.0261 | 1.488315 | 1.23E-05 | 0.001558 |
| LOC105373444 | 934.6957 | 1.506691 | 5.12E-14 | 1.00E-10 |
| RHEX | 176.6823 | 1.513286 | 0.000119 | 0.008576 |
| SLC27A2 | 241.4406 | 1.539782 | 2.84E-08 | 9.73E-06 |
| SGK1 | 313.493 | 1.650604 | 7.32E-11 | 6.27E-08 |
| MRC2 | 87.3865 | 1.652304 | 0.001032 | 0.04135 |
| PLAU | 118.7647 | 1.68301 | 4.65E-06 | 0.000724 |
| NTNG2 | 60.6708 | 1.699732 | 0.000701 | 0.03046 |

**Supplemental Table 8.** Genes upregulated in GCSF-mobilized cells from Population *b*

| Gene | baseMean | log2FoldChange | p-value | padj |
|---|---|---|---|---|
| AKAP2 | 703.6098 | 1.727609 | 5.90E-11 | 5.40E-08 |
| LGALS1 | 102.4688 | 1.780569 | 2.81E-05 | 0.002833 |
| SLC24A3 | 125.6366 | 1.841557 | 7.37E-05 | 0.006089 |
| ARHGAP10 | 109.3418 | 1.873864 | 3.72E-06 | 0.000623 |
| ANGPTL6 | 52.3411 | 2.070017 | 0.001047 | 0.041768 |
| TK1 | 55.49215 | 2.189955 | 0.001401 | 0.049925 |
| LOC101927497 | 176.7117 | 2.216348 | 1.66E-12 | 2.16E-09 |
| AFF2 | 134.0453 | 2.227865 | 1.93E-05 | 0.002177 |
| IGLL1 | 216.4632 | 2.391489 | 6.64E-13 | 1.01E-09 |
| MIR181A1HG | 210.1168 | 2.51486 | 1.85E-12 | 2.16E-09 |
| HPGDS | 125.47 | 2.718785 | 8.61E-10 | 4.73E-07 |
| ADAMTS14 | 54.90454 | 2.858097 | 0.000206 | 0.012618 |
| MFSD2B | 56.12507 | 2.90642 | 4.10E-05 | 0.003775 |
| HDC | 478.375 | 3.070035 | 0.001362 | 0.049488 |
| ACSM1 | 195.2779 | 3.107868 | 0.00013 | 0.009081 |
| CP | 172.1422 | 3.396736 | 1.44E-16 | 6.59E-13 |
| CPA3 | 615.8863 | 3.448173 | 1.33E-42 | 1.83E-38 |
| P2RX5 | 138.7401 | 3.455304 | 2.44E-10 | 1.60E-07 |
| TPSAB1 | 72.42388 | 6.084714 | 6.88E-06 | 0.000994 |

**Supplemental Table 9.** Summary of mobilization, leukapheresis and CD34 enrichment parameters

| Donor | GCSF dose | # collections | WBC count | CD34 count | CD34 purity [%] | Comment |
|---|---|---|---|---|---|---|
| 1 | 5mg/kg | 2 | 4.74e10 | 2.68e8 | 99.00 | Cryopreserved |
| 2 | 5mg/kg | 2 | 9.70e10 | 2.88e8 | 92.00 | Cryopreserved |
| 3 | 5mg/kg | 2 | 7.72e10 | 2.87e8 | 91.80 | Cryopreserved |
| 4 | 5mg/kg | 2 | 2.07e10 | 5.92e6 | 35.10 | Discontinued |
| 5 | 5mg/kg | 1 | 3.00E+10 | 1.30e8 | 97.00 | Fresh processing |
| 6 | 7.5mg/kg | 1 | 5.98e10 | 3.70e8 | 87.90 | Fresh processing |

All donors were selected for adjusted body weight >120% of ideal body weight.

**Supplemental Table 10.** Antibodies

| Antigen | Provider | Catalog Number | Clone Name | Lot Number | Fluorochrome | Application |
|---|---|---|---|---|---|---|
| CD3 | BD | 552851 | SP34-2 | 4346516 6092584 6336728 | Brilliant Violet 786 | Mouse BM, PB, Spleen, Thymus |
| CD4 | BioLegend | 300526 | RPA-T4 | B188454 B226779 B261177 B274110 | Alexa Fluor 700 | Mouse BM, PB, Spleen, Thymus |
| CD8 | BioLegend | 344742 | SK1 | B234630 B242754 B263073 | Brilliant Violet 605 | Mouse BM, PB, Spleen, Thymus |
| CD14 | eBioscience | 25-0149-42 | 61D3 | 4306573 E10277-1635 E10278-1637 | PE-Cy7 | Mouse BM, PB, Spleen, Thymus |
| CD15 | BioLegend | 323006 | W6D3 | B199835 B230031 | PE | Mouse BM, PB, Spleen, Thymus |
| CD16 | BD | 557758 | 3G8 | 4136850 5023818 6077649 6280745 7026993 7130902 7166692 8054938 8215752 | APC-Cy7 | Mouse BM, PB, Spleen, Thymus |
| CD19 | BD | 347544 | 4G7 | 5320803 6354963 7354554 | PerCP | Mouse BM, PB, Spleen, Thymus |
| CD20 | BioLegend | 302324 | 2H7 | 6294531 6214617 | PerCP | Mouse BM, PB, Spleen, Thymus |
| CD34 | BD | 562449 | 563 | 5070925 6027596 7053641 7166684 7348681 8238572 8242838 | PE-CF594 | Human HSPCs, Mouse BM |
| CD34 | BD | 561209 | 563 | 3151510 5070925 6027586 7110925 7348681 8087674 8242838 8283572 | APC | Mouse BM |
| CD38 | BioLegend | 303522 | HIT2 | B229737 B245218 | PerCP-Cy5.5 | Human HSPCs, Mouse BM |
| CD45 (hu) | BD | 560367 | HI30 | 6184678 7096590 | V450 | Human HSPCs, Mouse BM, PB, Spleen, Thymus |
| CD45 (mu) | BD | 562420 | 30-F11 | 6036634 6196640 7039832 7096986 7180875 8032908 | PE-CF594 | Mouse BM, PB, Spleen, Thymus |

**Supplemental Table 10.** Antibodies

| Antigen | Provider | Catalog Number | Clone Name | Lot Number | Fluorochrome | Application |
|---|---|---|---|---|---|---|
| | | | | 8281746<br>8294561 | | |
| CD45RA | BD | 561212 | 5H9 | 3046621<br>4220811<br>5093523<br>5239872<br>6091801<br>6343868<br>7082838<br>7110636<br>7222952<br>8236712 | APC-H7 | Human HSPCs, Mouse BM |
| CD56 | BioLegend | 318332 | HCD56 | B207590<br>B225062<br>B233132<br>B241283<br>B252245<br>B267124 | APC-Cy7 | Mouse BM, PB, Spleen, Thymus |
| CD90 | BioLegend | 328110 | 5E10 | B206722<br>B234526<br>B236754 | PE-Cy7 | Human HSPCs, Mouse BM |
| CD133 | Miltenyi Biotech | 130-080-801 | AC133 | 5140630511<br>5150126020<br>5161207109 | PE | Human HSPCs, Mouse BM |

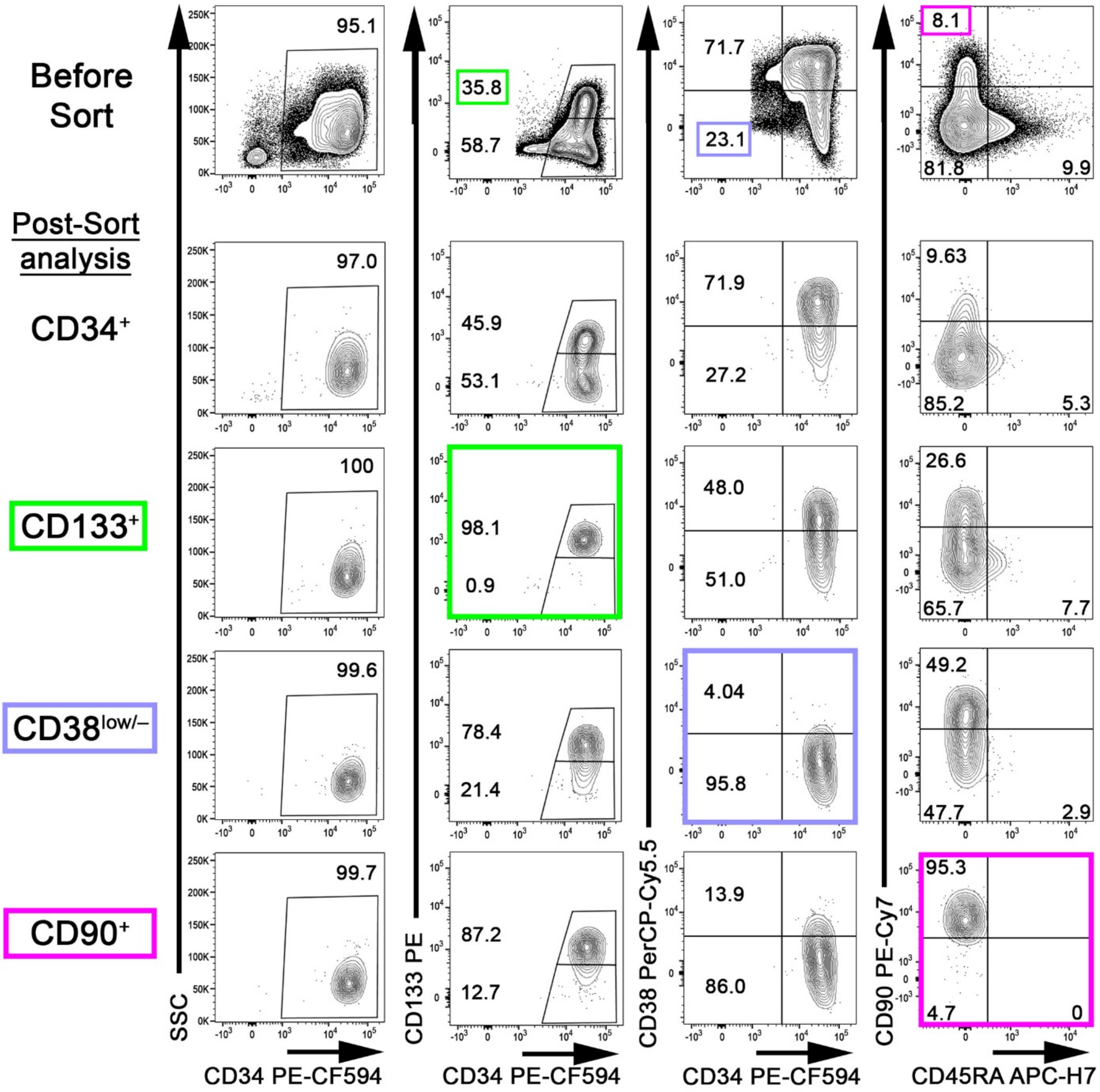


**Supplemental Figure 1. Quality control of sort-purified CD34-subpopulations.** Flow-cytometric quality control of bulk CD34$^+$ cells (top row, Before Sort) and sort-purified CD34$^+$ (2$^{nd}$ row), CD133$^+$ (3$^{rd}$ row), CD38$^{low/-}$ (4$^{th}$ row) and CD90$^+$ (5$^{th}$ row) HSPCs (Post-Sort analysis). Sorted target cell fractions are framed and color-coded. Numbers indicate frequency of gated population.

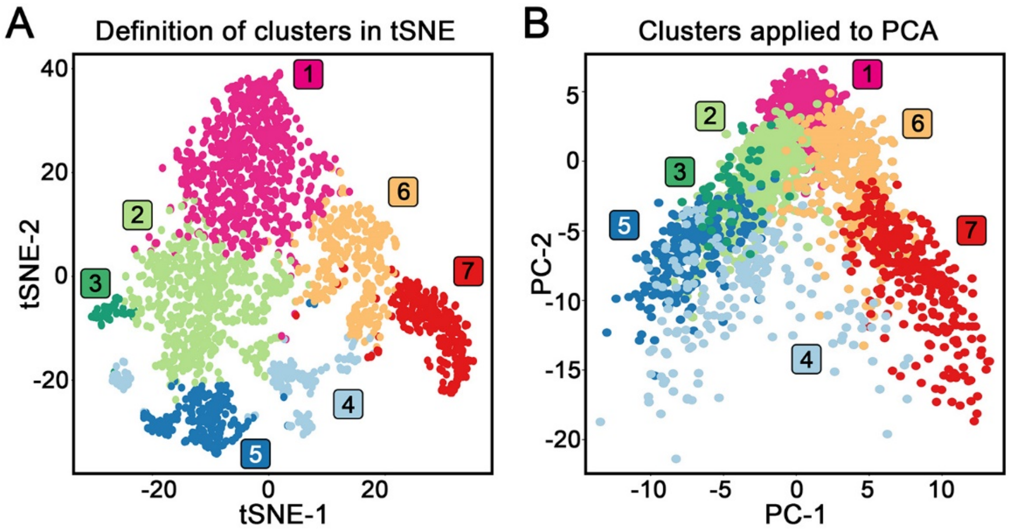


**Supplemental Figure 2. Transcriptionally distinct ssBM CD34 clusters in a second donor.** (**A**) Graph-based clustering of ssBM-derived CD34$^{+}$ cells. Transcriptionally distinct CD34 clusters were color-coded and numbered. (**B**) Clusters defined in A projected onto the PCA analysis.

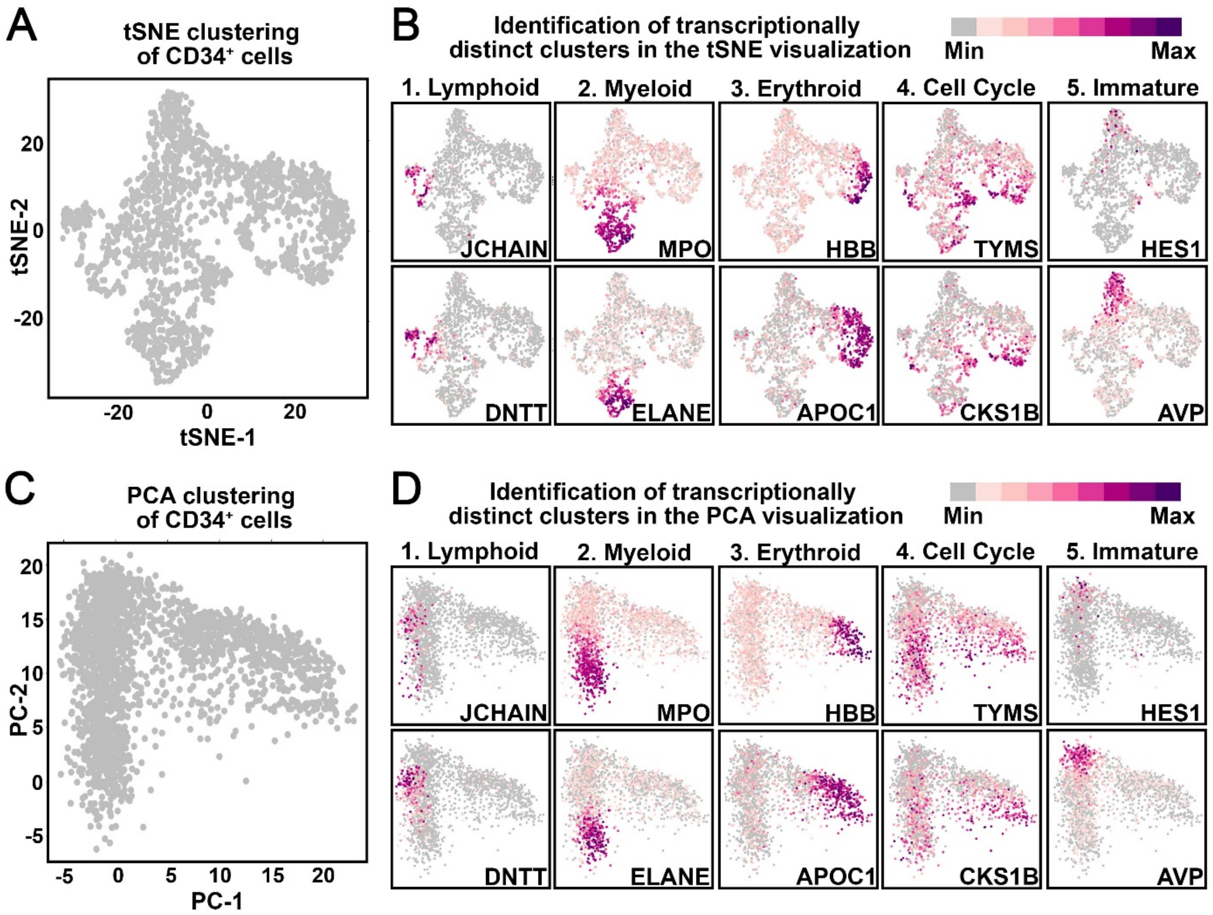


**Supplemental Figure 3. ScRNAseq of ssBM-derived CD34$^+$ HSPCs and sort-purified CD34 subsets.** (**A**) Dimensional reduction (tSNE) of scRNAseq data from ssBM-derived CD34$^+$ cells. (**B**) Feature plots showing the expression of representative genes associated with lymphoid-, myeloid-, erythroid-primed, proliferating, and immature HSPCs. Level of expression is color coded as shown in the legend. (**C**) PCA based transformation and (**D**) expression of representative genes for the same dataset shown in panel A.

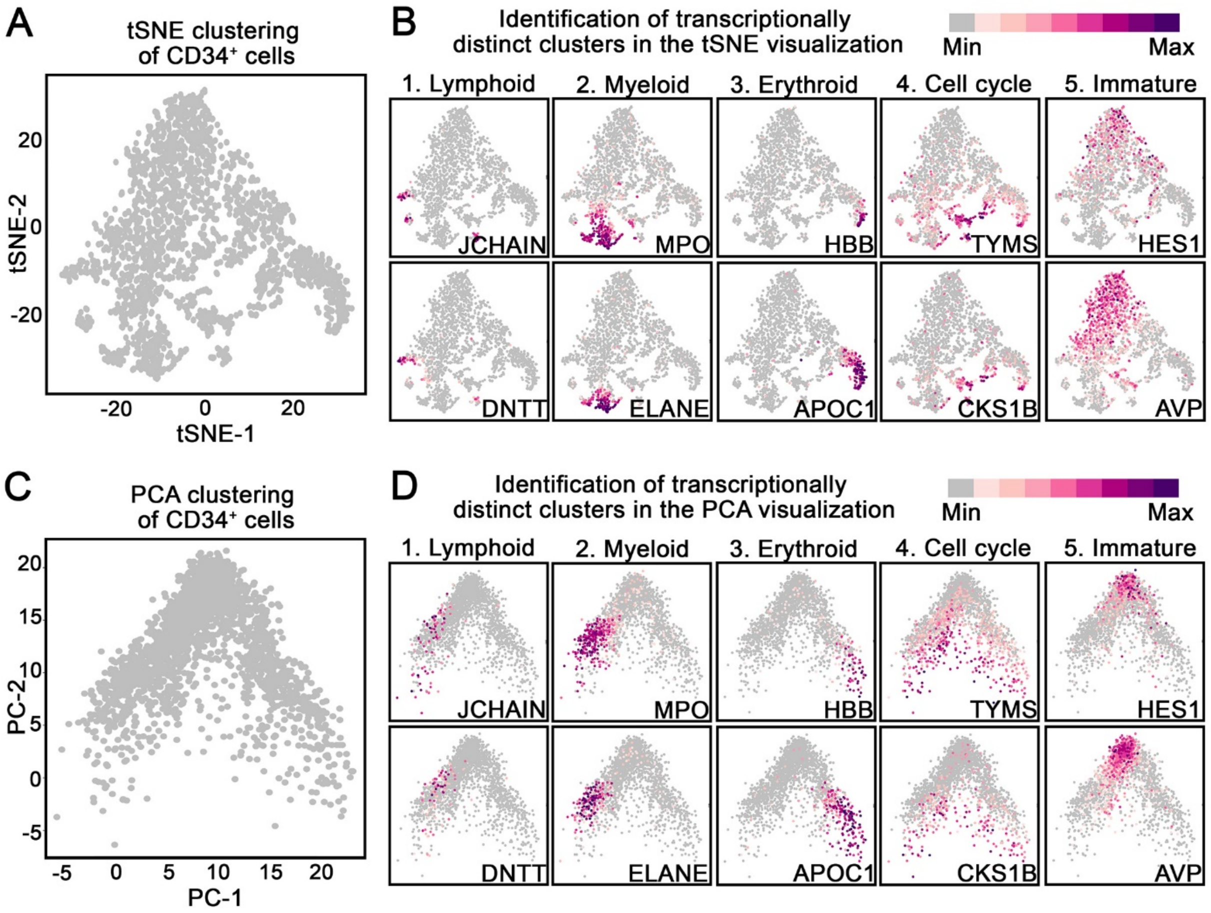


**Supplemental Figure 4. Donor-independent reproducibility of the scRNAseq ssBM reference map.** (**A**) tSNE and (**C**) PCA clustering of scRNAseq data from ssBM-derived CD34$^+$ cells from a second donor. (**B** and **D**) Feature plots showing the expression of representative genes associated with lymphoid-, myeloid-, erythroid-primed, proliferating, and immature HSPCs. Level of expression is color coded as shown in the legend.

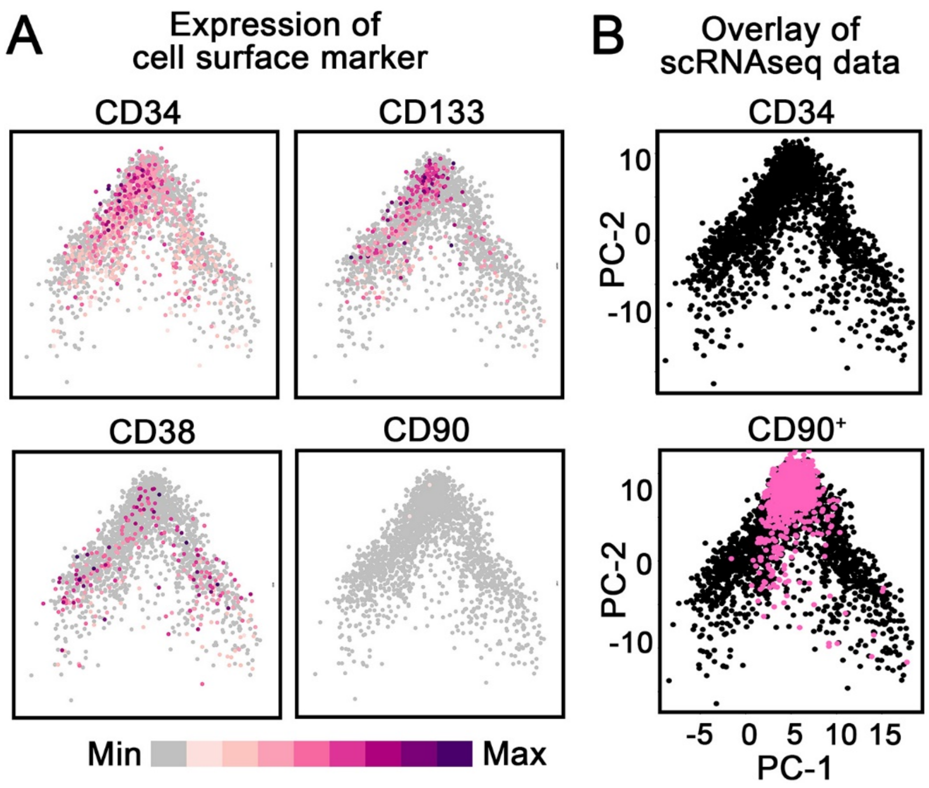


**Supplemental Figure 5: Transcriptional mapping of sort-purified CD34 subsets from a second donor.** (**A**) Expression of CD34, CD133, CD38 and CD90 in ssBM-derived CD34$^+$ cells. Level of expression is color coded as shown in the legend. (**B**) Overlay of scRNAseq data from CD34$^+$ cells (black, top plot) with sort-purified CD90$^+$ (pink, lower plot) HSPCs.

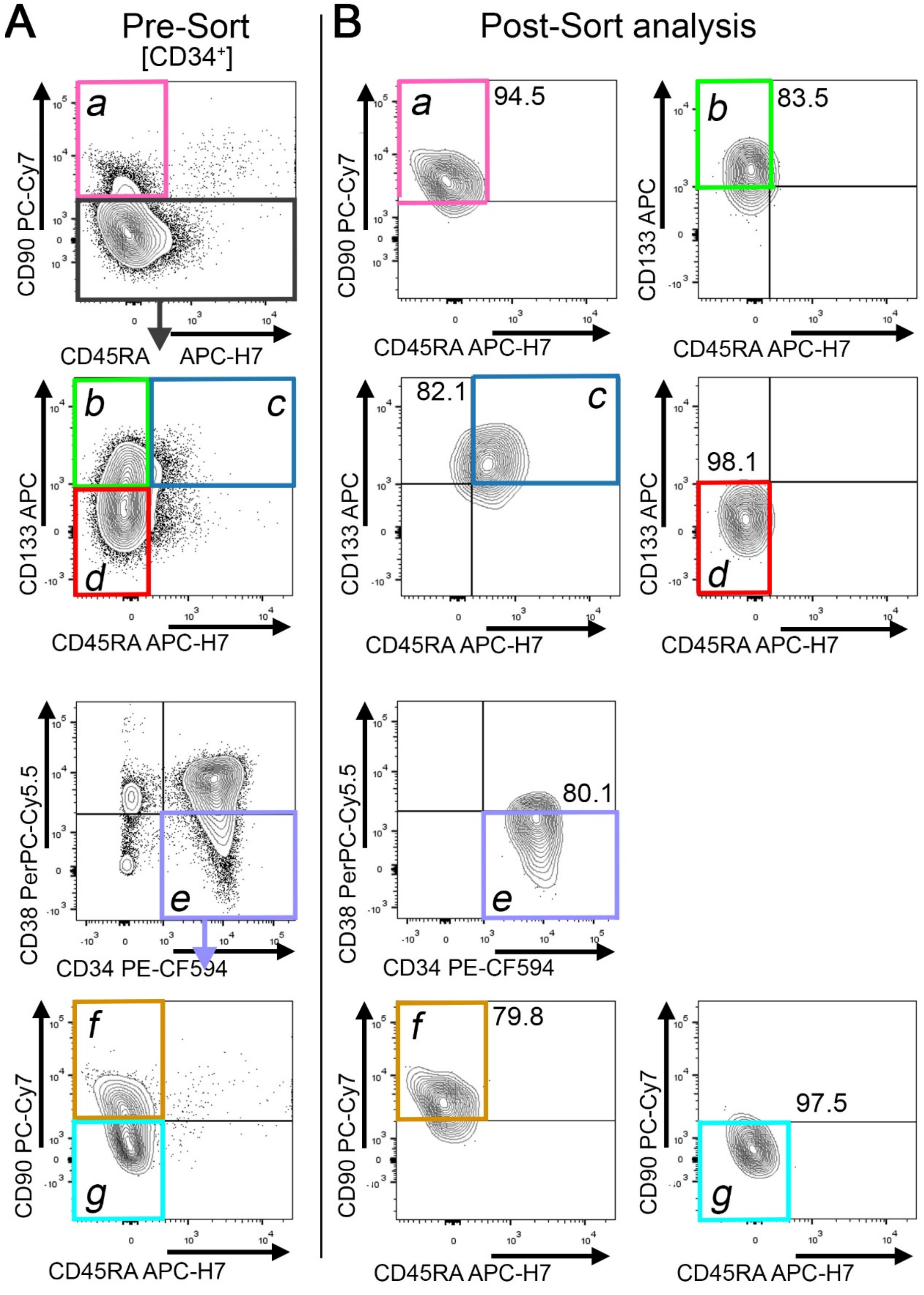


**Supplemental Figure 6. Quality control of sort-purified CD34-subpopulations for bulk RNAseq.** (**A**) Gating of ssBM-derived CD34 subpopulation defined in Figure 3A. (**B**) Flow-cytometric quality control of sort-purified CD34$^+$ subsets for bulk RNAseq. Sorted cell fractions are framed and color-coded. Numbers indicate frequency of gated population.

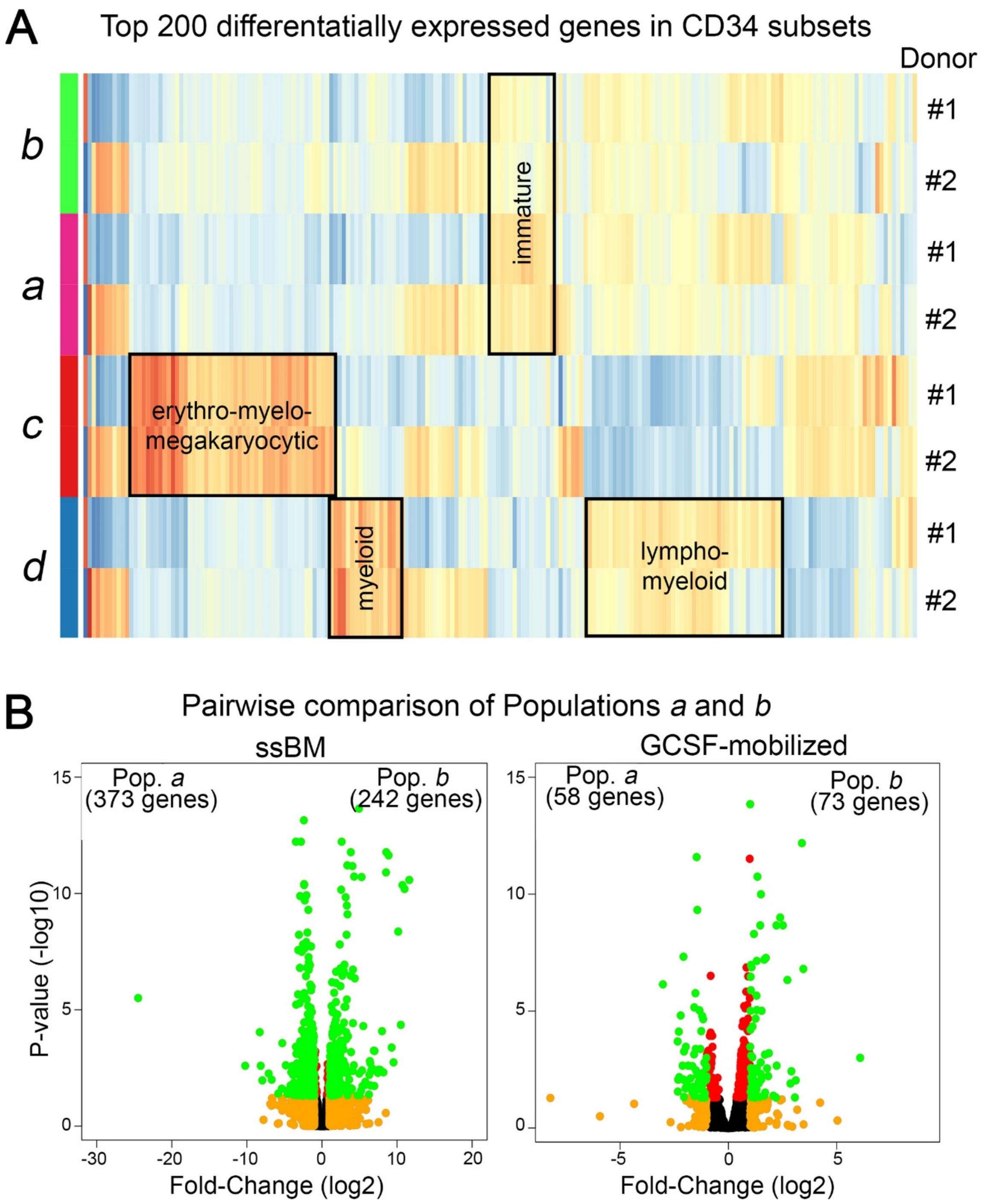


**Supplemental Figure 7. Differentially expressed genes in GCSF-mobilized bulk CD34 subsets.** (**A**) Heat map of the Top 200 differentially expressed genes in phenotypically-defined GCSF-mobilized CD34 subpopulations *a*–*d* from two independent human donors. A detailed list of the Top 200 genes can be found in **Table S4**. (**B**) Pair-wise comparison of the gene-expression in the ssBM and GCSF-mobilized subpopulations *a* and *b*. Differentially expressed genes are color coded according to the figure legend in the top left. A detailed list of all differentially expressed genes (green dots) can be found in **Tables S5, S6, S7 and S8**. Color-code: green = p-value < 0.05 and fold-change (FC) >1; red = p-value > 0.05 and FC >1; yellow = p-value > 0.05 and FC >1; black = p-value > 0.05 and FC < 1.

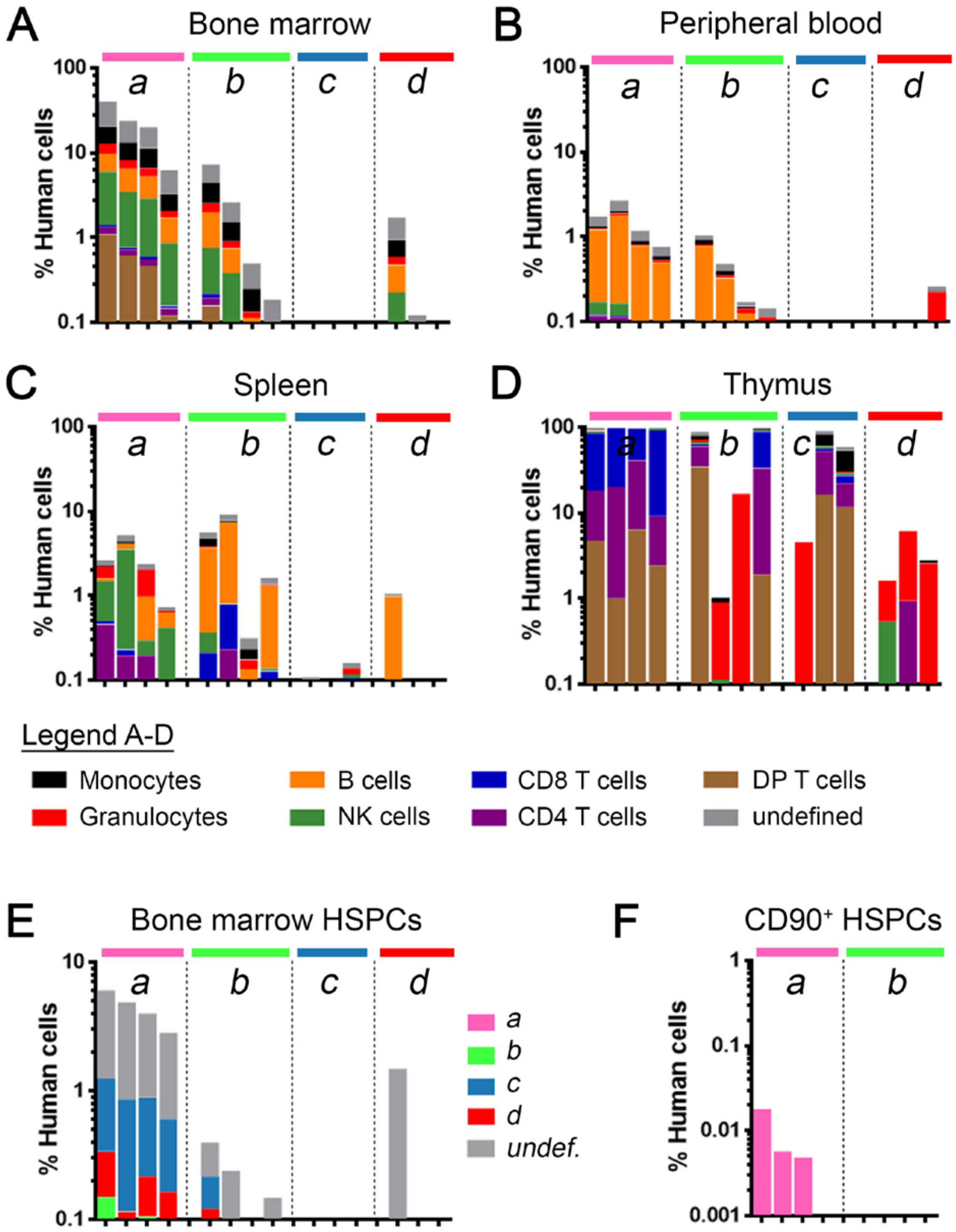


**Supplemental Figure 8. Multilineage engraftment potential of human CD34 subpopulations.** Human multilineage engraftment in the (**A**) BM, (**B**) PB, (**C**) spleen and (**D**) thymus after transplantation of sort-purified CD34 subpopulations (1e5 cells per mouse). Mice in all graphs and within each group are organized from the highest to the lowest engraftment level in the BM (A). (**E**) Frequency of human CD34$^+$ cells (total height of bars) and CD34$^+$ subpopulations (color-coded, as defined in Figure 3A). (**F**) Frequency of human CD90$^+$ HSPCs in the BM of mice transplanted with populations *a* and *b* only.

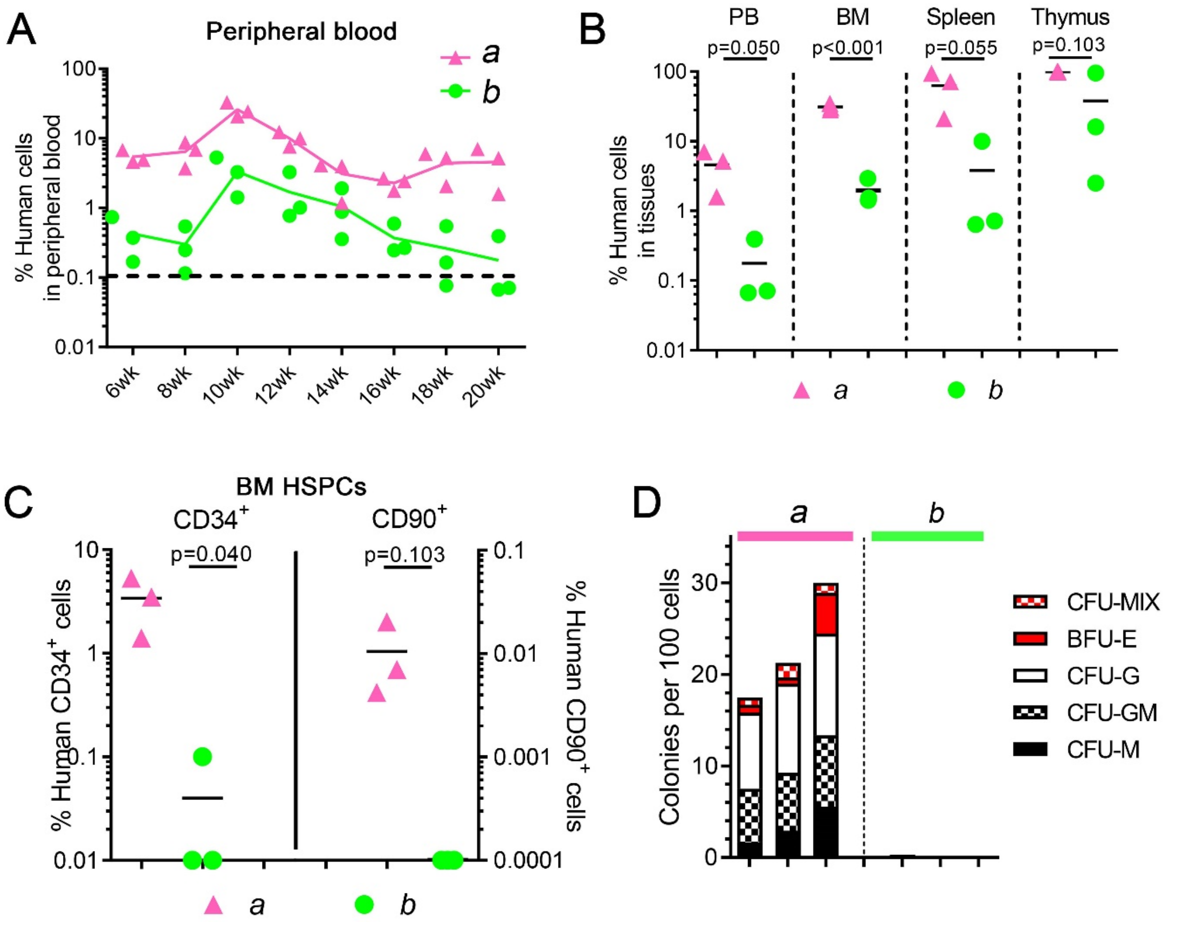


**Supplemental Figure 9. Engraftment potential of human CD34 subsets.** (**A**) Longitudinal tracking of human CD45+ engraftment in the PB of mice transplanted with $2.5\times10^5$ HSPCs cells per mouse from Population *a* or Population *b*. (**B**) Side by side comparison of human CD45+ engraftment in the PB, BM, spleen and thymus. PB and BM use left y-axis, spleen and thymus right y-axis. (**C**) Frequency of human CD34+ cells (left y-axis) and CD90+ HSPCs (right y-axis) in the BM of engrafted mice. (**D**) Erythroid, myeloid and erythro-myeloid colony-forming potential of engrafted human HSPCs.

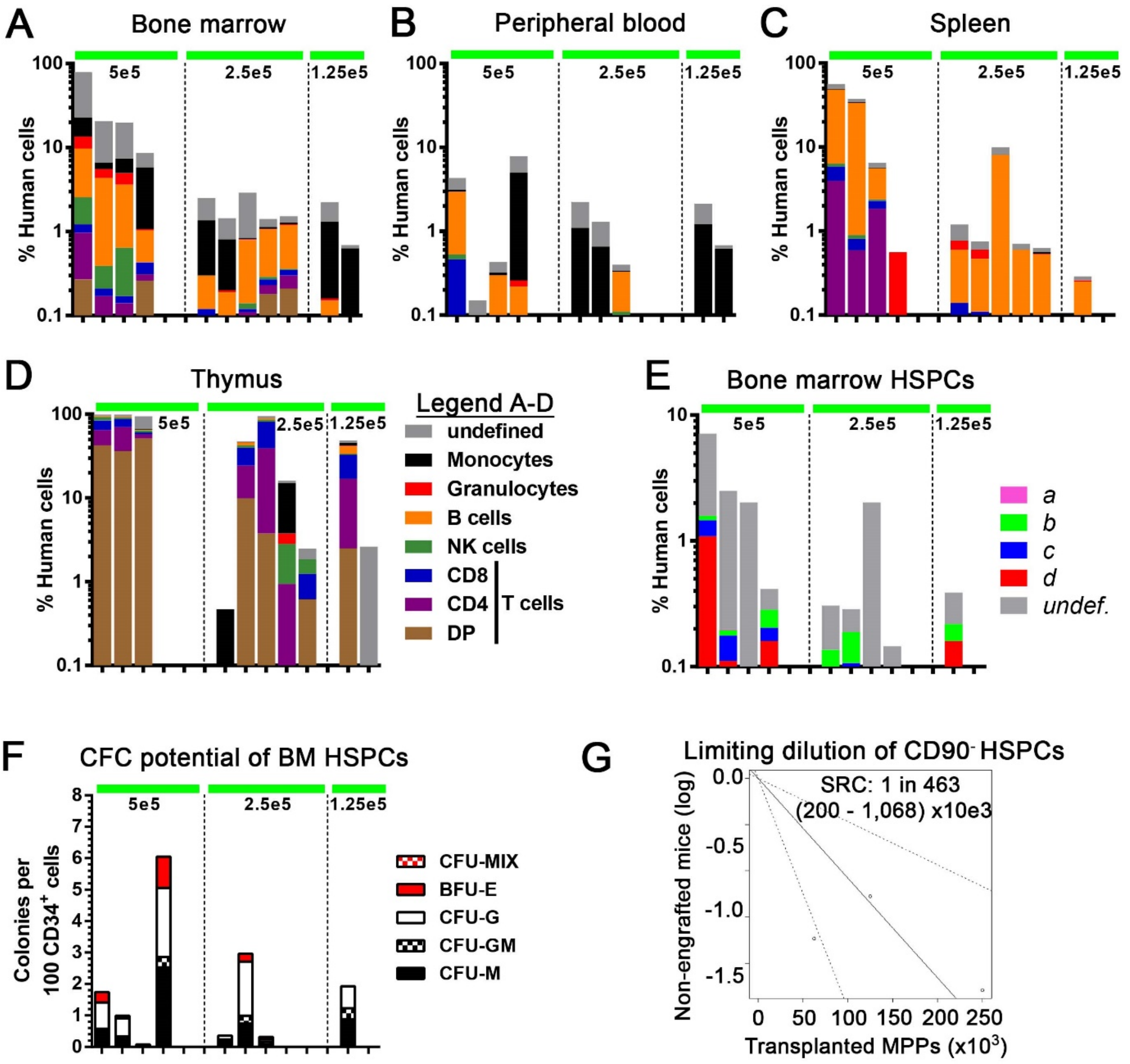


**Supplemental Figure 10. Engraftment potential of human HSPCs from Population *b*.** Human multilineage engraftment in the (**A**) BM, (**B**) PB, (**C**) spleen and (**D**) thymus. Mice in all graphs and within each group are organized from the highest to the lowest engraftment level in the BM (A). (**E**) Frequency of human $CD34^+$ cells (total height of bars) and $CD34^+$ subpopulations in the BM of transplanted mice. (**F**) Erythroid, myeloid and erythro-myeloid colony-forming potential of engrafted human HSPCs. (**G**) Calculation of human HSPCs from population *b* with SRC potential using a limiting dilution approach as previously described(51).

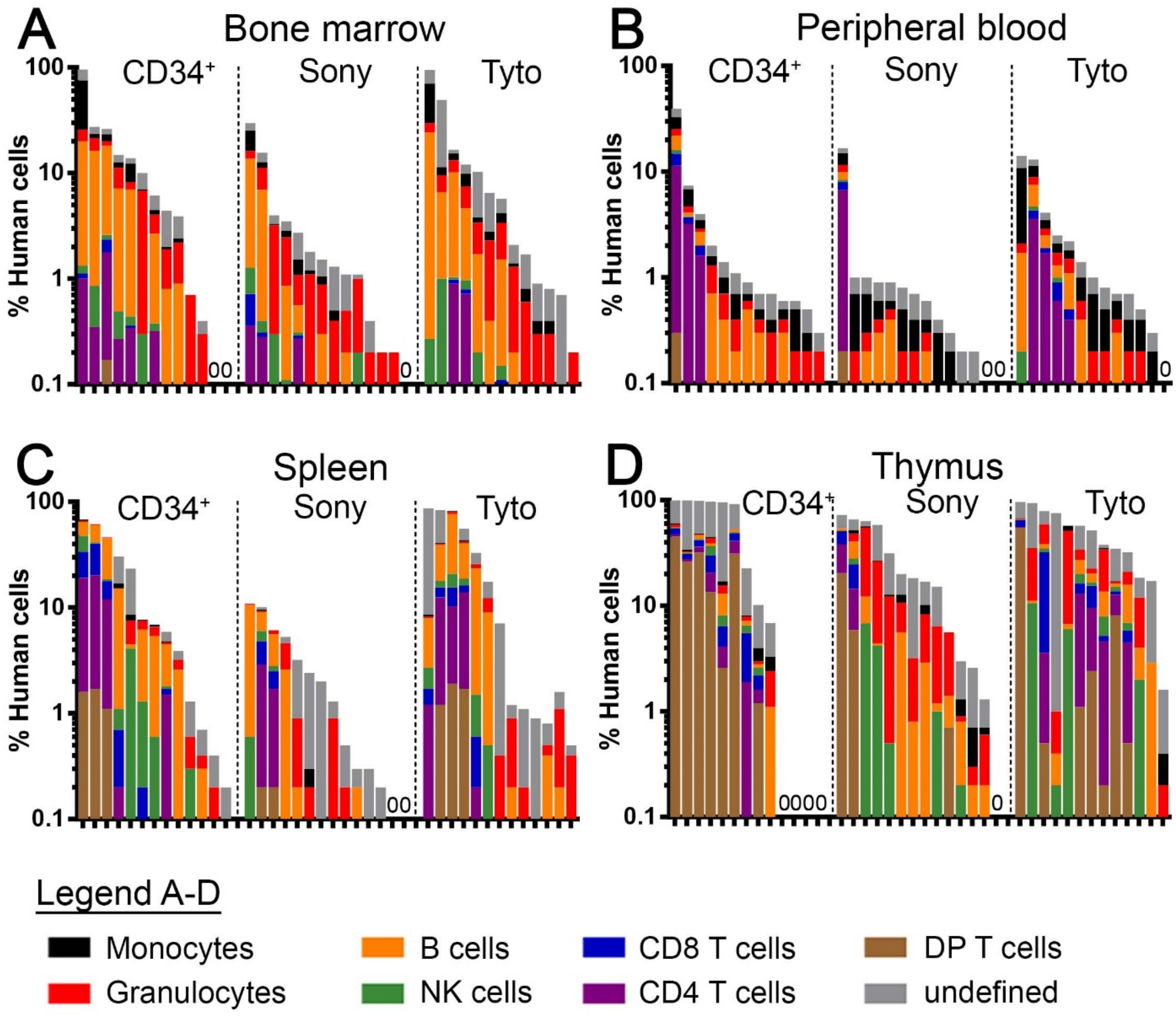


**Supplemental Figure 11. Engraftment potential of gene-modified human bulk CD34$^+$ and sort-purified CD34$^+$CD90$^+$ cells.** Human multilineage engraftment in the (**A**) BM, (**B**) PB, (**C**) spleen and (**D**) thymus of transplanted mice at 20 weeks post-transplant.
